# Endocardial primary cilia and blood flow are required for regulation of EndoMT during endocardial cushion development

**DOI:** 10.1101/2024.05.15.594405

**Authors:** Kathryn Berg, Joshua Gorham, Faith Lundt, Jonathan Seidman, Martina Brueckner

## Abstract

Blood flow is critical for heart valve formation, and cellular mechanosensors are essential to translate flow into transcriptional regulation of development. Here, we identify a role for primary cilia *in vivo* in the spatial regulation of cushion formation, the first stage of valve development, by regionally controlling endothelial to mesenchymal transition (EndoMT) via modulation of *Kruppel-like Factor 4 (Klf4)*. We find that high shear stress intracardiac regions decrease endocardial ciliation over cushion development, correlating with KLF4 downregulation and EndoMT progression. Mouse embryos constitutively lacking cilia exhibit a blood-flow dependent accumulation of KLF4 in these regions, independent of upstream left-right abnormalities, resulting in impaired cushion cellularization. snRNA-seq revealed that cilia KO endocardium fails to progress to late-EndoMT, retains endothelial markers and has reduced EndoMT/mesenchymal genes that KLF4 antagonizes. Together, these data identify a mechanosensory role for endocardial primary cilia in cushion development through regional regulation of KLF4.

## Introduction

Cardiac valves play a critical role in heart function and health by controlling blood flow directionality in the heart. Valve development happens in stages (reviewed in Geng et al. 2017^1^); briefly, small pockets of cardiac jelly are created, called endocardial cushions (ECC), that initially become cellularized through endothelial-to-mesenchymal transition (EndoMT). In this process, the endocardial luminal lining of the heart will differentiate into mesenchyme and migrate into the cardiac jelly, populating the ECC with interstitial cells. The cushion will then go through stages of remodeling, including mesenchymal proliferation and condensation, resulting in functional valve leaflets. In mice these two processes of cushion formation and remodeling begin at roughly e9 and e12.5, respectively^2^. Congenital heart disease (CHD) is a leading cause of birth defects^3^. Many CHDs have valve or septal defects linked to improper ECC formation, remodeling, or fusion during development^4,5^, an example being Hypoplastic Left Heart Syndrome (HLHS), a CHD characterized by extremely defective aortic and mitral valves directly tied to defective EndoMT^6^.

ECCs form at the atrio-ventricular and ventriculo-arterial junctions, areas of high wall shear stress (WSS) due to blood flow through a small luminal diameter^7^. Several lines of evidence link blood flow and mechanical forces to cardiogenesis, including ECC development (reviewed in^2,8–10^). Lack of contractility results in an inability to form an endocardial ring (the critical first step in ECC formation) in zebrafish and a complete loss of cushion cellularization in mice, indicating EndoMT requires mechanical stimuli^11,12^. Recently, Fukui et al. (2021)^13^ demonstrated that shear stress is both necessary and instructive for valve development in zebrafish. However, the direct mechanism by which blood flow is sensed and translated into genetic regulation of cushion formation during early heart valvulogenesis remains poorly understood, particulary in 4-chambered mammalian hearts.

Primary cilia are found on many cell types including endocardium and cushion mesenchyme^14–16^. Cilia in cardiac development are best known for their role in development of left-right asymmetry, where motile primary cilia function to generate extraembryonic fluid flow at the left-right organizer, and immotile primary cilia function as mechanosensors to detect flow and instruct the direction of cardiac looping ^17^. Immotile primary cilia are also essential during valve remodeling on cushion mesenchyme, however, whether they have a role in cushion formation remains unknown^18–20^. Even so, loss of cilia or ciliary signaling has been implicated in ECC defects irrespective of heterotaxy^17^, suggesting an important role for cilia during ECC development.

Work by Iomini et al. (2004)^21^ demonstrated shear stress-induced loss of primary cilia (deciliation) on cultured endothelial cells. Interestingly, the presence and subsequent loss of primary cilia has been shown to “prime” EndoMT in cultured HUVECs (human umbilical vein endothelial cells) through modulation of the mechanosensitive transcription factors *Kruppel-like Factors 2* and *4* (*Klf2, Klf4*, respectively)^22^. An *in vivo* link between intracardiac primary cilia and *Klf2a* expression has been shown in zebrafish cardiac regeneration^23^, but whether primary cilia are endocardial mechanosensors integrating mechanical signals and transcriptional regulation of murine cushion formation *in vivo* is currently unknown.

Here we demonstrate a novel role for primary cilia in murine heart cushion formation. Flow-dependent, dynamic ciliation correlates with spatially-selective regulation of KLF4 and subsequent EndoMT. Single-nuc RNAseq data shows transcriptional consequences of constitutive cilia loss during early heart development, including dysregulation of KLF4 targets and EndoMT pathways. EndoMT initiates, but does not progress in cilia KO embryos, and the cushions are decellularized in both embryos lacking cilia, and embryos lacking flow. This highlights a requirement for primary cilia as potential flow sensors guiding EndoMT during heart formation independent of left-right patterning, and upstream of valve remodeling.

## Results

### Endocardial ciliation of the ECCs is spatiotemporally dynamic during cushion EndoMT

Primary cilia can be lost (‘deciliation’) in response to high laminar flow^21,22^. Since the ECC centers experience high shear stress, we first evaluated if ECC endocardial cells possess cilia during cushion formation (e8.5 to e10). Primary cilia are present on the endocardial lining in both the OFT and the AVC **(Fig. 1A, Fig. S1A)** throughout cushion development. These cilia are anywhere from 0.3-1 uM in length and can be found extending into the lumen **(Fig. 1B)**. Between e8.5 and e9.0, we observed a decrease in total endocardial ciliation (percentage of cells with a cilium) in both ECCs, suggesting a deciliation event following the onset of laminar flow at e8.25 **(Fig. 1D)**. However, total endocardial ciliation in the ECCs remains constant after this initial event until the end of cushion EndoMT, even as shear stress intensity increases. This initial loss and subsequent plateau of endocardial ciliation appears to be evolutionarily conserved, as we observed similar results in live zebrafish embryo AVCs during developmentally equivalent time points **(Fig. S1B, S1C)**.

**Figure 1.**
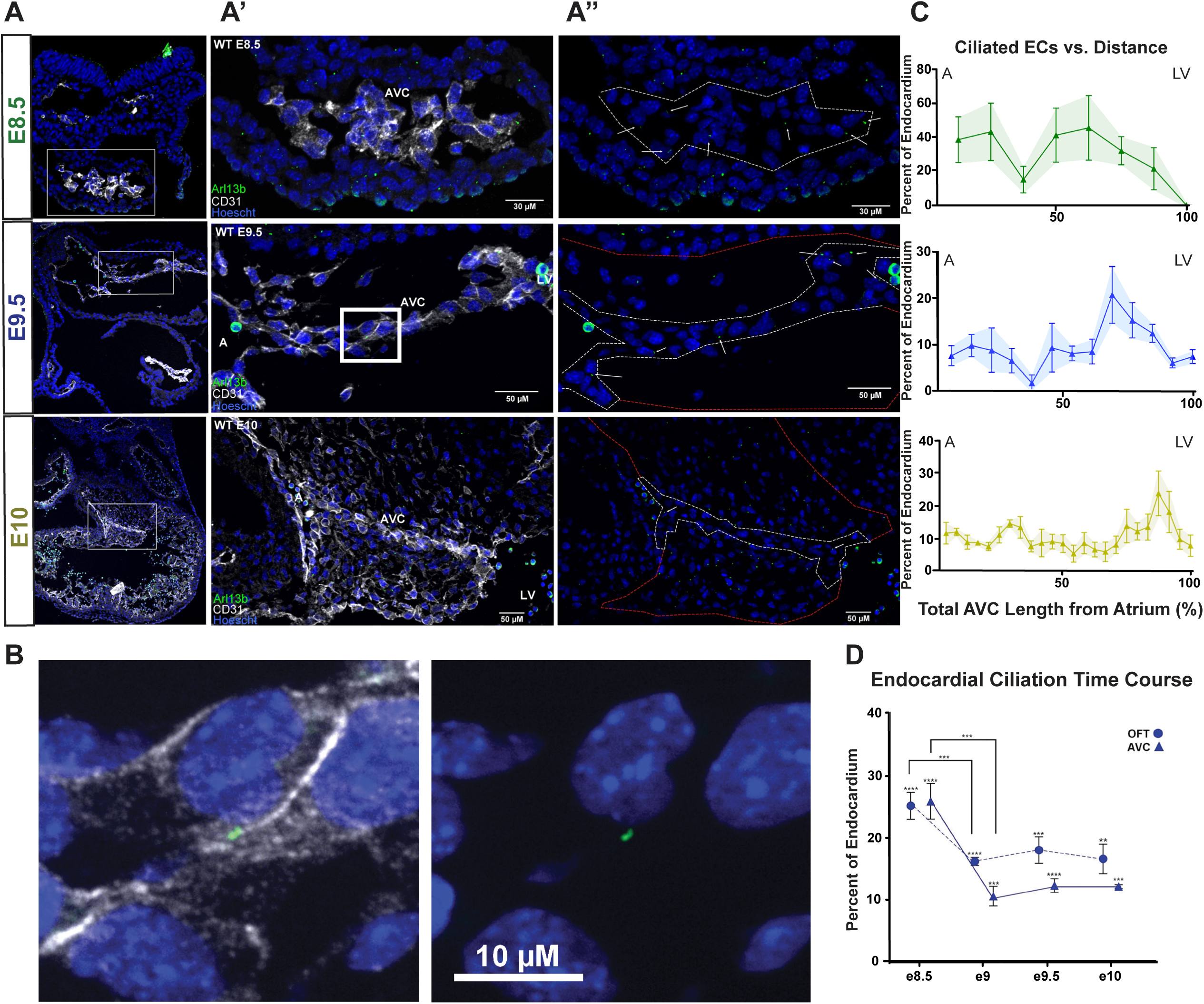
Endocardial ciliation of the ECCs is spatiotemporally dynamic during cushion EndoMT. **A)** Immunofluorescence on mouse embryo sections for cilia (ARL13B, green) on endocardial cells (CD31, white) of the AVC over cushion development (e8.5, e9.5, e10). Nuclei are shown in blue (Hoescht). Close ups **A’)** with and **A”)** without CD31; white arrows indicate cilia; luminal endocardial cells outlined with white and whole AVC (including cushion mesenchyme) outlined in red. **B)** Closeup of white box in **A’** showing a single cilium on an e9.5 endocardial cell. **C)** Ciliated endocardial cells as percentage of all endocardial cells versus distance in the AVC at e8.5 (n=3), e9.5 (n=4), and e10 (n=3). Distance runs left to right, Atrium to Left Ventricle. **D)** Endocardial ciliation over time as percentage of all endocardial cells in either the AVC or OFT (e8.5 (AVC n=10, OFT n=10), e9 (AVC n=7, OFT n=7), e9.5 (AVC n=6, OFT n=8), e10 (AVC n=4, OFT n=4)). Statistics: ** (p ≤ 0.01), *** (p ≤ 0.001), **** (p ≤ 0.0001). Data are represented as mean ± SEM. Abbreviations: A-Atrium, AVC-Atrioventricular Canal, EC-Endocardial Cell, LV-Left Ventricle, OFT-Outflow Tract, WT-Wildtype.

Endocardial ciliation across the developing AVC was dynamic between e8.5 and e10. Ciliation is increasingly absent at the AVC center and retained at the ventricular, and to a lesser degree atrial, junction **(Fig. 1C)**. The areas of low ciliation correlate with areas of high WSS^24^. This ciliation pattern becomes more regionally restricted over the course of AVC development, and by e9.5 also emerges in the OFT cushions **(Fig. S1D)**. Cells that have undergone EndoMT and migrated into the cardiac jelly away from the lumen regain ciliation **(Fig. S1E, S1F)**. These results identify two populations of luminal endocardial cells during cushion formation: ciliated cells at the lower WSS edges of the cushions, and nonciliated cells in the centers.

### The mechanosensitive transcription factors KLF2 and KLF4 are involved in human valve development and are dynamically expressed in mouse endocardium during cushion EndoMT

Beyond a mechanism by which endocardium can sense fluid flow, the mechanosensitive endocardial programs that ultimately drive cushion EndoMT are still poorly understood. To discover mechanosensitive genetic regulators involved in valve development, we assessed human variants of mechanosensitive factors in 11,555 patients with a broad range of CHD, identifying significant enrichment for dominant loss-of-function and damaging missense variants affecting Kruppel-Like Factors 2 and 4 (KLF2, KLF4) in patients possessing aortic and mitral valve defects associated with HLHS (Sierant et al, in preparation, **Table S1**).

KLF2 and KLF4 are mechanosensitive zinc-finger containing transcriptions factors that have overlapping functions in vascular development^25^, but upregulate in response to mechanical forces differently^26^. Our analyses showed different expression patterns of *Klf2* and *Klf4* mRNA during cushion development. Both *Klf2* and *Klf4* mRNA are expressed in the endocardial luminal lining of the AVC and OFT **(Fig. 2A, 2A”, 2B, 2B”, S2A, S2A”, S2B, S2B”)**. While *Klf4* was restricted to the ECCs, *Klf2* was more widely expressed, including in the intersomitic vessels, cardinal vein, and the ventricle, dorsal aorta, and atrium **(Fig. 2A’, 2B’, S2A’, S2B’, 2C)**, consistent with KLF2’s well defined role in vascular identity^27^. Total *Klf2* and *Klf4* expression increased following the onset of contraction **(Fig. 2D)**. Reanalysis of scRNAseq data^28^ demonstrates that in contrast to *Klf2*, *Klf4 (*which is required earlier in development as a pluripotency factor^29^), only becomes predominantly expressed in the endocardium between e8.25 and e9.25 **(Fig. 2E)**. Both *Klf2* and *Klf4* expression was lower in the OFT compared to the AVC until e9.5 **(Fig. 2D);** KLF4 protein positive cells in the OFT increases to that of the AVC by e10 **(Fig. S2C)**. This timing difference in expression between the two ECCs is consistent with the known developmental delay between the OFT and AVC^30^.

**Figure 2.**
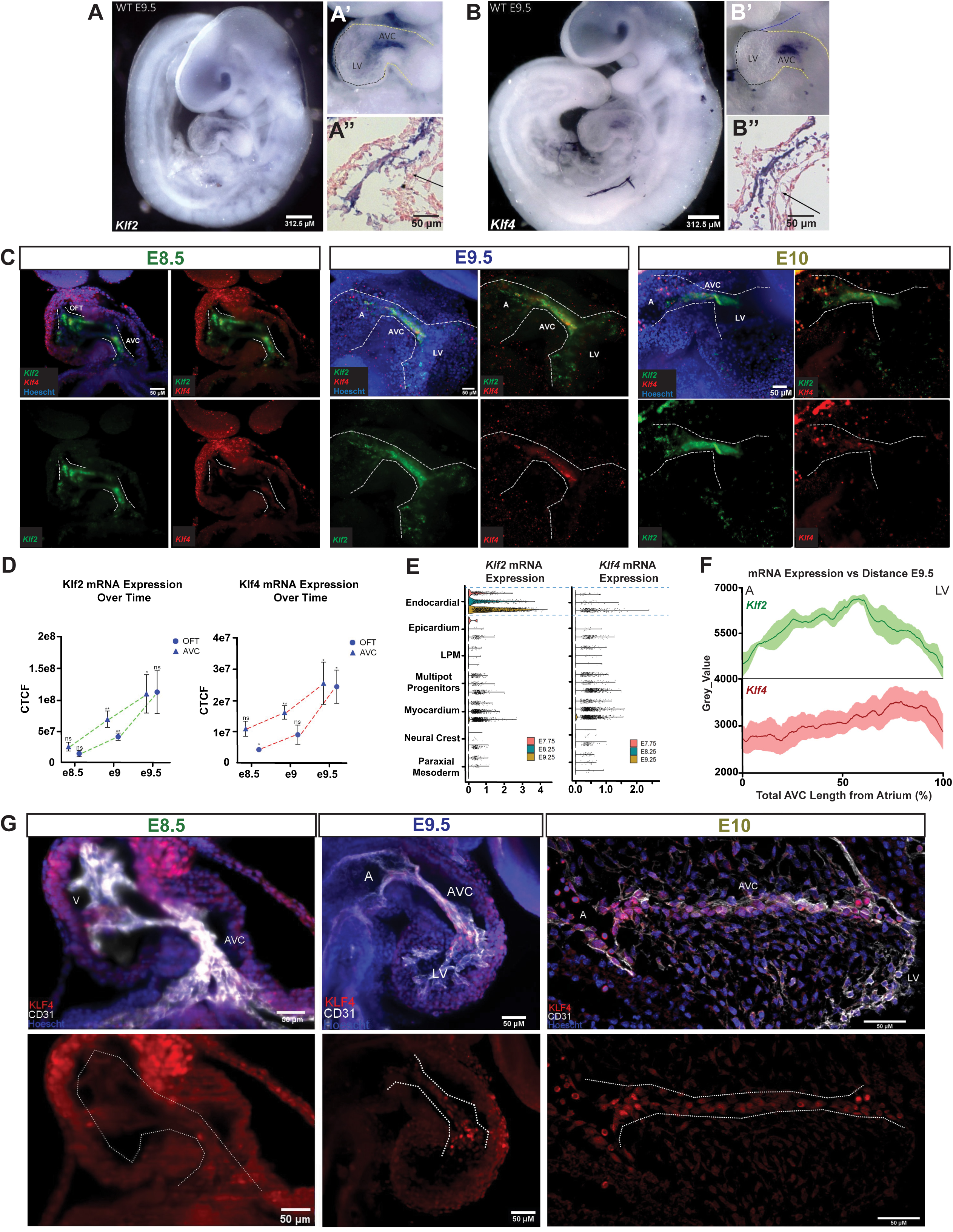
The mechanosensitive transcription factors KLF2 and KLF4 are dynamically expressed in mouse endocardium during cushion EndoMT. Whole mount ISH of e9.5 mouse embryos for A) *Klf2* mRNA and B) *Klf4* mRNA with closeup **(A’, B’)** and cryosection **(A”, B”)** of the AVC (black arrow). **C)** HCR-FISH on whole mount mouse embryos at e8.5, e9.5, and e10 showing *Klf2* mRNA (green)*, Klf4* mRNA (red), and nuclei (Hoescht, blue). AVC and OFT are outlined in white. **D)** Corrected total cell fluorescence (CTCF) analysis of HCR-FISH over time for *Klf2* and *Klf4* in the AVC and OFT (*Klf2:* e8.5 (AVC n=3, OFT, n=3), e9 (AVC n=5, OFT, n=4), e9.5 (AVC n=4, OFT, n=3) (*Klf4:* e8.5 (AVC n=3, OFT, n=3), e9 (AVC n=5, OFT, n=4), e9.5 (AVC n=4, OFT, n=3). **E)** Single-Cell RNAseq^28^ of mouse hearts showing total *Klf2* mRNA and *Klf4* mRNA expression in cardiac cell types at e7.75, e8.25, and e9.25. **F)** mRNA expression as measured by Grey_Value over distance for *Klf2* (n=4) and *Klf4* (n=5) in the e9.5 AVC. Distance runs left to right, Atrium to Left Ventricle. **G)** Immunofluorescence on mouse embryos for KLF4 protein (red) in endocardial cells (CD31, white) of the AVC over time (e8.5, e9.5, e10). Nuclei are shown in blue (Hoescht). KLF4 protein alone is shown with endocardium outlined in white. *Statistics*: ns (p > 0.05), * (p ≤ 0.05), ** (p ≤ 0.01), *** (p ≤ 0.001), **** (p ≤ 0.0001). Data are represented as mean ± SEM. Abbreviations: A-Atrium, AVC-Atrioventricular Canal, EC-Endocardial Cell, LPM-Lateral Plate Mesoderm, LV-Left Ventricle, OFT-Outflow Tract, WT-Wildtype.

*Klf2* expression across the AVC endocardium resembled a bell curve mirroring known WSS levels^24,31^ **(Fig. 2F)**. In contrast, *Klf4* exhibited more mRNA expression and KLF4 protein positive cells on the ends of the AVC, especially on the ventricular side **(Fig. 2F, 2G)**. This high KLF4 expressing cluster of cells is first noticeable in small amounts at e8.5, but takes up ∼20% of the total AVC length starting at e9.5. A second peak of KLF4 protein positive cells can also be seen on the atrial end by e10. A similar ventricular pattern can be observed in the OFT, and like the ciliation pattern it is not as robust as in the AVC **(Fig. S2D, S2E)**. This data highlights two distinct populations of KLF4-expressing cells: KLF4-low cells with low KLF4 expression in the center of the AVC and KLF4-high cells with high KLF4 expression levels on the ventricular and atrial ends.

### KLF4 expression at the AVC correlates with endocardial ciliation and depends on blood flow

KLF4 is a mechanosensitive transcription factor that has been shown to respond to shear-stress induced activation of TRP channels/PIEZO1 and Notch1 in vascular endothelium^32,33^. The mechanical forces in the developing ECCs are complex, and include circumferential stress caused by contraction against the ECC junctions, laminar flow in the center of the ECC and disturbed flow at the ECC junctions (reconstruction of previously published data^7,24,31,34,35^shown in **Fig. 3A**). KLF4 is not induced by circumferential stress^36^ or disturbed flow^37,38^, so we propose an endocardial mechanosensor capable of discriminating and responding to a range of WSS as the driving difference between KLF4-low and KLF4-high regions. Since primary cilia are present on the endocardium in a WSS-specific manner, we explored the possibility of intracardiac primary cilia filling this role by comparing the locations of KLF4-high, KLF4-low, ciliated and nonciliated cells in the AVC. Ciliated cells overlap with KLF4-high regions, and nonciliated with KLF4-low regions **(Fig. 3B, 3C)**, and this correlation increases over cushion development **(Fig. 3D)**. There is also a positive correlation between the amount of KLF4 protein in an endocardial cell and being ciliated **(Fig. 3E)**. Taken together, these results integrate ciliation and KLF4 expression into two cell populations: KLF4-high, ciliated and KLF4-low, nonciliated.

**Figure 3.**
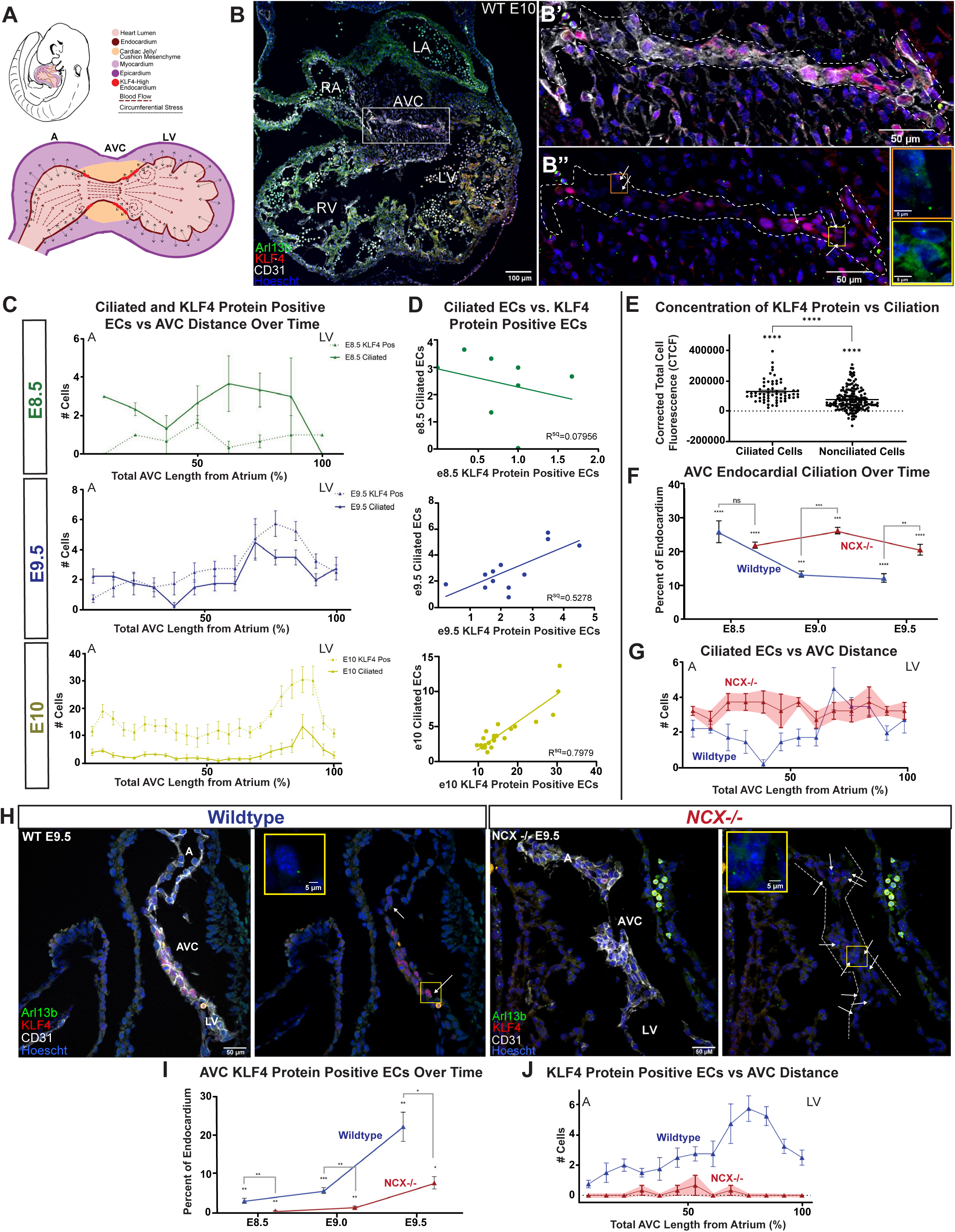
KLF4 expression at the AVC correlates with endocardial ciliation and depends on blood flow. **A)** A model of mechanical forces in an e9.5 mouse heart. Red arrows indicate blood flow/shear stress, grey arrows indicate contractile stress. **B)** Immunofluorescence on e10 mouse heart sections for KLF4 protein (red) and cilia (ARL13B, green) on endocardial cells (CD31, white). Nuclei are shown in blue (Hoescht). Closeup of boxed region in B) of the AVC **B’)** with and **B”)** without CD31; white arrows indicate cilia; endocardium is outlined in white. Closeups of endocardial cilia are shown in orange and yellow boxes. **C)** KLF4 protein positive endocardial cells (dotted line) and ciliated endocardial cells (solid line) versus distance in the AVC at e8.5 (n=3), e9.5 (n=4), and e10 (n=3). **D)** XY plots for correlation of average plot points from C). **E)** Violin plot for KLF4 protein immunofluorescent signal intensity (CTCF) in ciliated and nonciliated AVC endocardial cells. **F)** Endocardial ciliation in the AVC over time as percentage of all endocardial cells in wildtype mice (blue) and *Ncx1^−/−^* mice (red) (e8.5 (WT n=8, *Ncx1^−/−^* n=8), e9.0 (WT n=4, *Ncx1^−/−^* n=4), e9.5 (WT n=7, *Ncx1^−/−^* n=7)). **G)** Ciliated endocardial cells versus distance in the AVC of *Ncx1^−/−^* mice at e9.5 (n=4). For comparison, WT line is provided in blue. **H)** Immunofluorescence on e9.5 wildtype and *Ncx1^−/−^* mouse heart sections for KLF4 protein (red) and cilia (ARL13B, green) on endocardial cells (CD31, white). Nuclei are shown in blue (Hoescht). Version without CD31 is provided; white arrows indicate cilia; endocardium is outlined in white. Closeups of endocardial cilia are shown in yellow boxes. **I)** KLF4 protein positive endocardial cells in the AVC over time as percentage of all endocardial cells in wildtype mice (blue) and *Ncx1^−/−^* mice (red) (e8.5 (WT n=10, *Ncx1^−/−^* n=10), e9.0 (WT n=6, *Ncx1^−/−^* n=6), e9.5 (WT n=4, *Ncx1^−/−^* n=4)). **J)** KLF4 protein positive endocardial cells versus distance in the AVC of *Ncx1^−/−^* mice at e9.5 (n=4). For comparison, WT line is provided in blue. For **C)**, **G)**, and **J)**, distance runs left to right, Atrium to Left Ventricle. *Statistics*: ns (p > 0.05), * (p ≤ 0.05), ** (p ≤ 0.01), *** (p ≤ 0.001), **** (p ≤ 0.0001). Data are represented as mean ± SEM. Abbreviations: A-Atrium, AVC-Atrioventricular Canal, EC-Endocardial Cell, LA-Left Atrium, LV-Left Ventricle, OFT-Outflow Tract, RA-Right Atrium, RV-Right Ventricle, WT-Wildtype.

HUVECs reduce KLF4 following a high flow-dependent loss of primary cilia^22^. To test if our regional loss of cilia and KLF4 expression in the AVC was due to high flow, we examined ECC ciliation in *Ncx1^−/−^* mutant mouse embryos lacking cardiac contractility through loss of the Na^+^/Ca^2+^ exchanger protein^39^. *Ncx1^−/−^* mutant embryos are viable until e9.75, and at e9.5 do not exhibit an externally visible cardiac phenotype beyond the lack of a heartbeat. Unlike wild-type embryos, endocardial cilia were retained and distributed ubiquitously across the AVC of *Ncx1^−/−^* mutant embryos from e8.5-e9.5 **(Fig. 3F-H)**, and the amount of KLF4 protein positive cells decreased **(Fig. 3H, 3I)**. The distribution of the limited KLF4 positive cells did not replicate the ventricular bias of the wildtype, but instead as small peaks in the center of the AVC **(Fig. 3J)**. The OFT had identical results **(Fig. S3A, S3B)**. These data demonstrate that blood flow is necessary for regional ciliation and KLF4 expression.

### KLF4 expression negatively correlates with cushion EndoMT progression

EndoMT is required for ECC development, and KLF4 has been shown to inhibit or activate EndoMT ^22,40–43^. To decouple KLF4’s role in EndoMT, we examined the expression patterns of *Vimentin* (*Vim*, mesenchymal marker) and *Kinase Insert Domain Receptor* (*Kdr,* endothelial marker) to test whether KLF4-low cells in the center of the AVC were more or less likely to undergo EndoMT. Double positive cells (considered to be undergoing EndoMT) were abundant in the center of the AVC at E9.5 **(Fig. 4A)**. *Vim* expression decreases in the terminal ∼25% of the AVC, mirroring the KLF4-high cell cluster localization **(Fig. 4B)**. Cushion cellularization was also first noticeable in the center of the AVC and could be detected at low levels by E9.0 and increased until E10 **(Fig. 4C)**.

**Figure 4.**
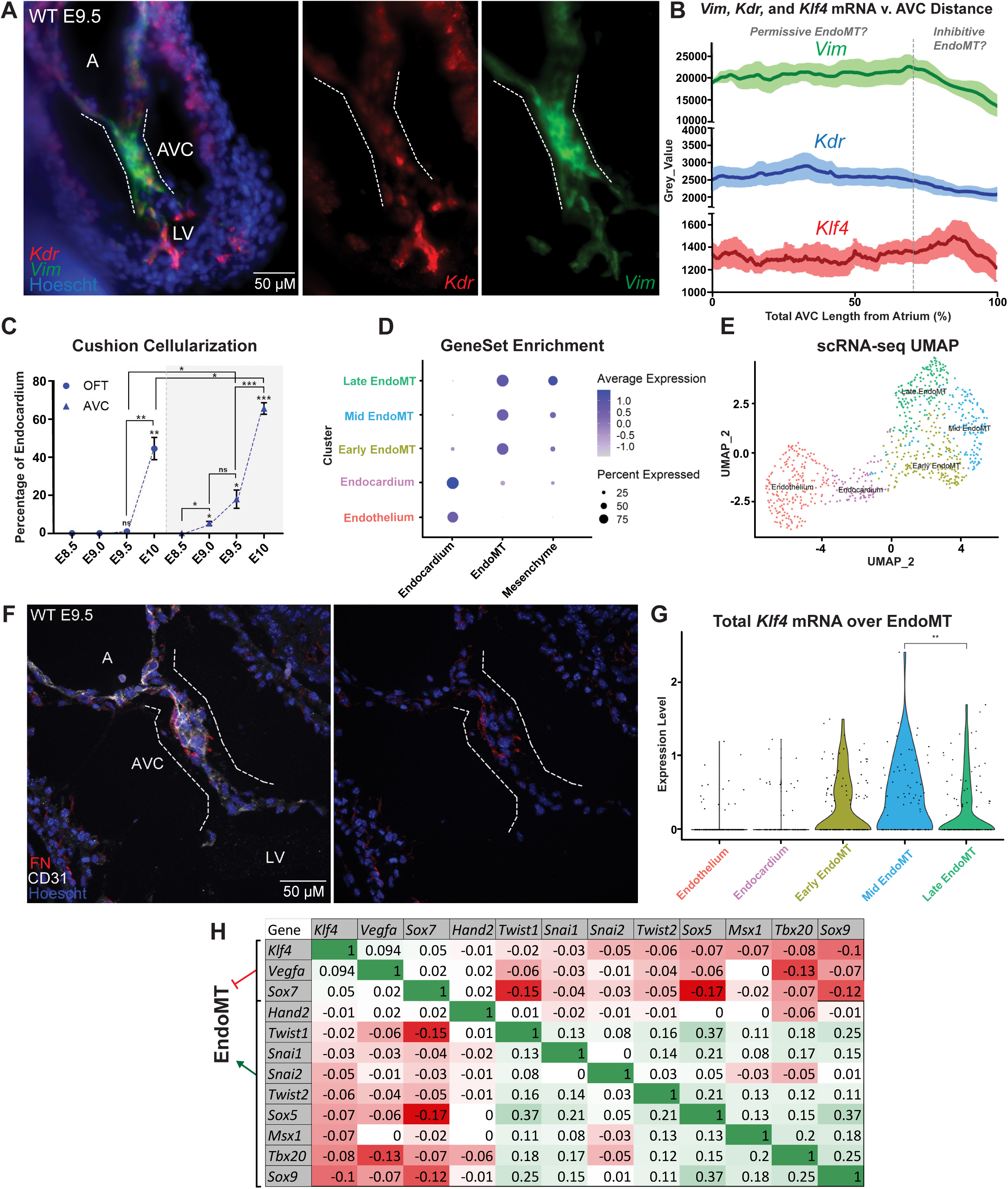
KLF4 expression negatively correlates with cushion EndoMT progression. **A)** HCR-FISH on whole mount mouse embryos at e9.5 showing *Kdr* mRNA (red)*, Vim* mRNA (green), and nuclei (Hoescht, blue). The AVC is outlined in white. **B)** mRNA expression as measured by Grey_Value over distance for *Kdr* (blue, n=4), *Vim* (green, n=4) and *Klf4* (red, n=6) in the e9.5 AVC. Distance runs left to right, Atrium to Left Ventricle. **C)** Cushion cellularization as a percentage of total CD31 positive endocardial cells in the AVC and OFT over time (e8.5 (AVC n=3, OFT, n=3), e9 (AVC n=4, OFT, n=4), e9.5 (AVC n=4, OFT, n=4), e10 (AVC n=4, OFT n=5)). **D)** SingleCell RNAseq^28^ GeneSet Enrichment for EndoMT progression. **E)** UMAP plot of SingleCell RNAseq^28^ EndoMT clusters. **F)** Immunofluorescence on e9.5 wildtype mouse heart sections for FN protein (red) in endocardial cells (CD31, white). Nuclei are shown in blue (Hoescht). Version without CD31 is provided; the AVC is outlined in white. **G)** Total *Klf4* mRNA expression over EndoMT pseudotime from SingleCell RNAseq^28^. **H)** Correlation matrix between *Klf4* and EndoMT regulators; mRNA expression from SingleNuc RNAseq. *Statistics*: ns (p > 0.05), * (p ≤ 0.05), ** (p ≤ 0.01), *** (p ≤ 0.001). Data are represented as mean ± SEM. Abbreviations: A-Atrium, AVC-Atrioventricular Canal, EC-Endocardial Cell, LV-Left Ventricle, OFT-Outflow Tract, WT-Wildtype.

To explore how endocardial *Klf4* expression correlates with EndoMT progression at the transcriptional level, we defined distinct pseudo-stages of EndoMT through measuring the ratio of Endocardial and Mesenchymal GeneSet enrichments on a previously published scRNA-seq dataset^28^ **(Fig. 4D, 4E)**. These data provide an atlas of gene expression networks associated with EndoMT progression **(Table S2A-D)**. This includes markers of each pseudo-stage, such as quiescent EC (*Tmem100, Emcn,* and *Gja4)*, early-EndoMT (*Wnt4, Fn1, Cd9,* and *Fabp5)*, mid-EndoMT (*Trpm3, Edn1, Bmper,* and *Col23a1),* and late-EndoMT (*Pdgfra, Vcan, Snai1,* and *Twist1/2)*, reminiscent of previously published EndoMT pseudotime analysis^44^. Immunofluorescent staining of Fibronectin (FN), identified in our dataset as an early EndoMT marker (consistent with Mjaatvedt et al. (1987)^45^), was found exclusively at the center of the AVC where cellularization first occurs **(Fig. 4F)**, suggesting that EndoMT begins in the center of the AVC.

Total *Klf4* mRNA peaks in mid-EndoMT and significantly decreases by late-EndoMT **(Fig. 4G)**. Since KLF4 inhibits EndoMT factors *Snai1* and *Snai2 (Slug)*^32,46,47^, we investigated nascent *Snai1* and *Snai2* expression in cells undergoing EndoMT by performing sn-RNAseq on micro-dissected e9.5 wildtype mouse hearts. Analysis of this snRNA-seq confirmed a negative correlation between *Klf4* and *Snai1*, *Snai2,* and multiple other essential EndoMT genes, such as *Twist1, Twist2*, *Hand2, Tbx20, Msx1*, and *Sox9*, which predominately peak in expression at late-EndoMT **(Fig. 4H**). EndoMT antagonists and endothelial genes, such as *Vegfa* and *Sox17*, were positively correlated to *Klf4.* Together these data suggest that the transition from mid- to late-EndoMT is linked to downregulation of *Klf4* expression.

### Failure of ciliogenesis results in abnormal KLF4 expression and failure to cellularize the ECCs

Given the strong correlation between endocardial KLF4 expression and ciliation in ECCs, we tested whether cilia are necessary for normal ECC KLF4 expression. We utilized two independent constitutive cilia knockout mouse models, *Kif3a^−/−^* (Kinesin Family Member 3A, responsible for anterograde intraflagellar transport) and *Ift20^−/−^* (Intraflagellar Transport Protein 20, responsible for transport of ciliary components from the golgi to the cilium)^48,49^, both of which lead to complete loss of cilia via distinct mechanisms **(Fig. 5A, S4A, S4B, S5A)**. Both *Kif3a^−/−^* and *Ift20^−/−^* mutant embryos are viable until e9.75 and have several defects, such as randomized left-right asymmetry, pericardial edema, and loss of neural tube closure.

**Figure 5.**
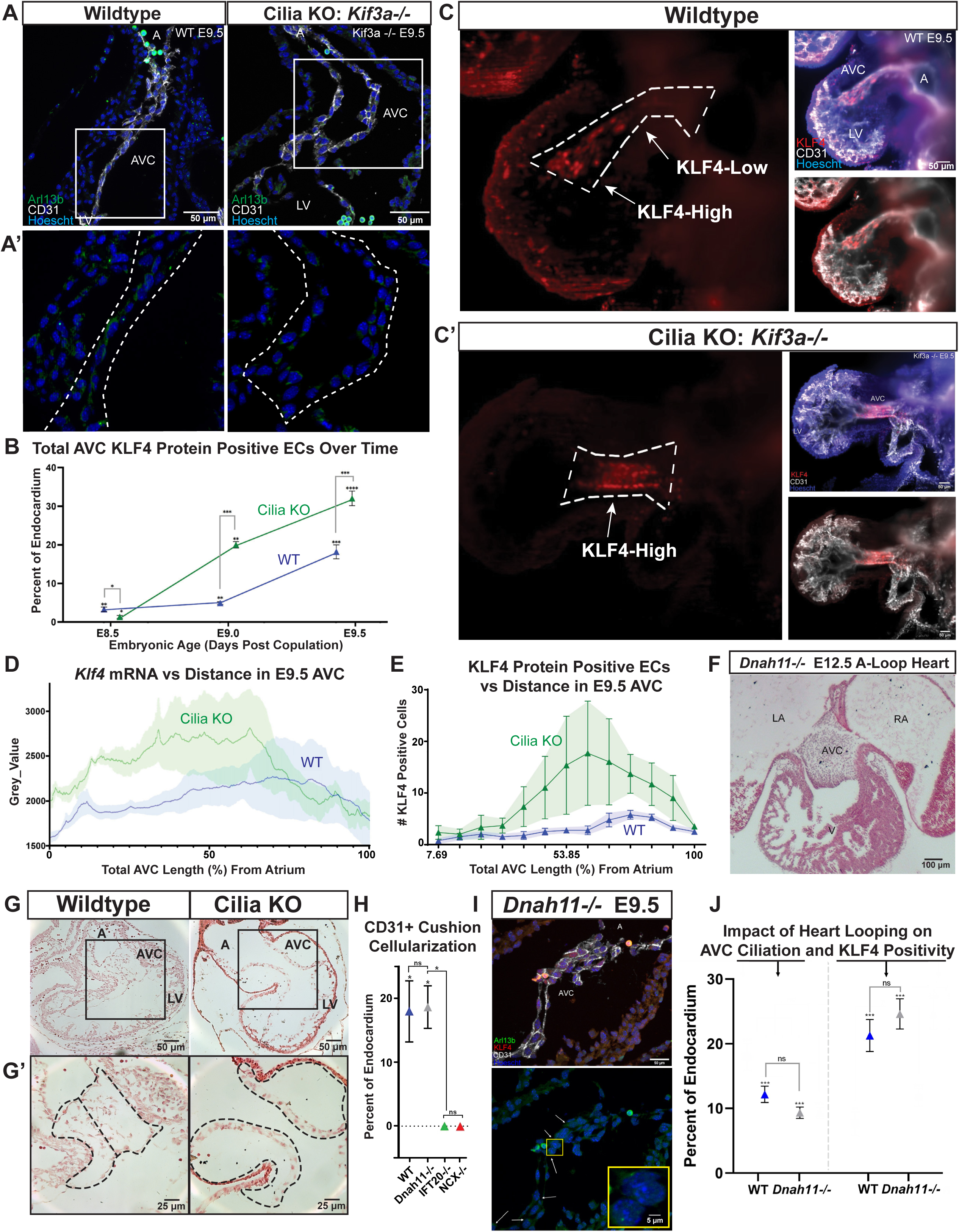
Failure of ciliogenesis results in abnormal KLF4 expression independent of direction of heart looping. **A)** Immunofluorescence on e9.5 wildtype and cilia KO (*Kif3a^−/−^)* mouse heart sections for cilia (ARL13B, green) on endocardial cells (CD31, white). Nuclei are shown in blue (Hoescht). A’) Closeup of boxed AVC region without CD31; the AVC is outlined in white. **B)** KLF4 protein positive endocardial cells in the AVC over time as percentage of all endocardial cells in wildtype mice (blue) and cilia KO mice (green) (e8.5 (WT n=6, cilia KO n=7), e9.0 (WT n=3, cilia KO n=4), e9.5 (WT n=6, cilia KO n=6)). Immunofluorescence on e9.5 **C)** wildtype and **C’)** cilia KO (*Kif3a^−/−^)* whole mount hearts for KLF4 (red) in endocardial cells (CD31, white). Nuclei are shown in blue (Hoescht); the AVC is outlined in white. **D)** *Klf4* mRNA expression as measured by Grey_Value over distance in the AVC of e9.5 wildtype mice (blue, n=3) and cilia KO mice (green, n=4). **E)** KLF4 protein positive endocardial cells versus distance in e9.5 AVCs of wildtype mice (blue, n=4) and cilia KO mice (green, n=3). **F)** H&E stain of an e12.5 *Dnah11^−/−^* A-looped mouse heart. **G)** Nuclear Fast Red stain of e9.5 wildtype and cilia KO mouse hearts with **G’)** closeups of boxed regions; cushions outlined in black. **H)** Cushion cellularization as a percentage of total endocardial cells in the AVC of wildtype (blue, n=4), *Dnah11^−/−^* (grey, n=4), cilia KO (green, n=5) and *Ncx1^−/−^* (red, n=4) hearts. **I)** Immunofluorescence on e9.5 *Dnah11^−/−^* mouse hearts for KLF4 protein (red) and cilia (ARL13B, green) on AVC endocardial cells (CD31, white). Nuclei are shown in blue (Hoescht). Cilia and nuclei alone are given; white arrows indicate cilia. Closeups of endocardial cilia are shown in yellow box. **J)** Ciliated endocardial cells and KLF4 protein positive endocardial cells in the AVC as percentage of all endocardial cells in e9.5 wildtype (blue) and *Dnah11^−/−^* (grey) mice (cilia: WT n=6, *Dnah11^−/−^* n=6; KLF4: WT n=6, *Dnah11^−/−^* n=6). For **D)** and **E)**, distance runs left to right, Atrium to Left Ventricle. *Statistics*: ns (p > 0.05), * (p ≤ 0.05), ** (p ≤ 0.01), *** (p ≤ 0.001), **** (p ≤ 0.0001). Data are represented as mean ± SEM. Abbreviations: A-Atrium, AVC-Atrioventricular Canal, EC-Endocardial Cell, LV-Left Ventricle, OFT-Outflow Tract, WT-Wildtype.

KLF4 expression was indistinguishable comparing *Ift20^−/−^* and *Kif3a^−/−^* mouse hearts, so they will be shown aggregated and collectively referred to as cilia KO **(Fig. S4C, S5B)**. At e8.5, cilia KO hearts appear phenotypically normal apart from randomization of the direction of cardiac looping but have fewer KLF4 positive cells than wildtype littermates **(Fig. 5B, S5C),** consistent with previous observations that *Klf4 mRNA* expression was reduced in cilia KD HUVECs^22,50^. Conversely, cilia KO mutants develop significantly higher *Klf4* mRNA and number of KLF4 positive cells at e9.0. The KLF4-low cell population at the center of the AVC observed in wildtype littermates is lost in cilia KO embryos, leaving only KLF4-high cells **(Fig. 5C, S4D, S4E)**. The KLF4-high cells are symmetrically distributed across the center of the AVC of e9.5 cilia KO hearts. This contrasts with the ventricular-biased distribution of KLF4-high AVC cells observed in wildtype littermates **(Fig. 5D, 5E).** Cilia KO OFTs do not differ in KLF4 expression to the same degree as AVCs but have a similar expression shift towards the OFT center **(Fig. S5D, S5E)**. This suggests that cilia are required to direct spatial gene expression of the developing ECCs, resulting in an inability to downregulate KLF4 in the center of the AVC or OFT in cilia KO hearts. By creating a double *Ift20/Ncx1* KO line, we were able to confirm that the overexpression of KLF4 in cilia KO is dependent on contractility and subsequent laminar flow. *Ift20/Ncx1* KO are not significantly different than single *Ncx1^−/−^* hearts in terms of KLF4 expression **(Fig. S5F, S5G)**, suggesting that blood flow-dependent deciliation lies upstream of KLF4.

To determine the impact of constitutive cilia loss on EndoMT, we examined cushion cellularization in wildtype and cilia KO heart nuclear-counterstained sections and observed a striking lack of interstitial cells in cilia KO cushions. **(Fig. 5G)**. To confirm interstitial cells were of endocardial origin we measured the proportion of cells that had infiltrated the cushion while also retaining CD31 expression **(Fig. 5H)**. Cilia KO mutants show 0% cellularization of the AVC by e9.5 (identical to *Ncx1^−/−^* embryos), compared to an average 17% cellularization in wildtype. Cilia KOs expressed fibronectin (FN), an early EndoMT marker, at comparable amounts to wildtype in the AVC endocardium **(Fig. S5H, S5I)**. Combined with the absence of mesenchymal cells in the cilia KO AVC, this indicates that cilia KO embryos can initiate, but not complete, EndoMT in the AVC by e9.5.

### Abnormal ECCs in ciliogenesis mutant embryos are not due to abnormal direction of heart looping

Cilia KOs have random heart looping direction due to a requirement for cilia motility and sensing at the left-right organizer (LRO) preceding AVC development. To determine whether KLF4 overexpression in cilia KO hearts was due to hemodynamic and structural changes secondary to variable heart looping, instead of a direct effect of AVC intracardiac cilia sensing, we evaluated *Dnah11^−/−^* (Dynein Axonemal Heavy Chain 11) mouse embryos. Constitutive knockout of *Dnah11* results in immotile LRO cilia and identical heart looping phenotypes as observed in ciliary KO mice without affecting intracardiac primary cilia^51^. *Dnah11^−/−^* embryos are viable and show normal cellularization of the AVC endocardial cushion, even in A-loop hearts, suggesting EndoMT and cushion formation are unaffected by variable heart looping **(Fig. 5F)**. Endocardial ciliation in *Dnah11^−/−^* embryos appears identical to that of wildtype siblings, both in total amount and regionalization across the AVC **(Fig. 5I, 5J, S5J)**. Importantly, *Dnah11^−/−^* embryos have no change in KLF4 expression when compared to wildtype embryos **(Fig. 5J, S5J)**. Together, these data demonstrate that KLF4 misexpression in ciliary KO embryos is not due to upstream LR defects but is instead caused by loss of intracardiac cilia sensing during cardiogenesis. This suggests that there is a previously unknown dependence on endocardial ciliation for proper cushion development independent of the development of cardiac left-right asymmetry.

### Defects in ciliogenesis block EndoMT

We then sought to identify the transcriptional differences that underlie the observed failure to progress through EndoMT in cilia KO embryo hearts. We performed single-nucleus RNAseq on micro-dissected e9.5 hearts from *Ift20^−/−^* and sibling (heterozygous and wildtype) embryos **(Fig. 6A)**. We obtained 30,823 single nuclei (29,313 after QC) with 27,998 genes over 2 replicates (Nuclei counts: *Ift20^+/+^* 9,882; *Ift20^+/−^* 14,825; *Ift20^−/−^* 6,116). Established marker genes were used to identify cluster identities, and the proportion of major cell types were not significantly different between genotypes **(Fig. 6B, 6D, S6A, S6B, S6D, S6E)**. Since we had observed increased KLF4 protein in cilia KO endocardium, we analyzed whether this is reflected in expression levels of known KLF4 targets in the endocardial cluster. These comprise genes that are activated by KLF4 including *Lama3* (Laminin subunit)^52^, *Malat1* (Metastasis Associated Lung Adenocarcinoma Transcript 1)^53^, *Vegfa*, and *Vegfc* (Vascular Endothelial Growth Factors)^54–56^, and genes known to be repressed by KLF4 including *Col3a1* (Collagen)^57^, *Pdgfrα* (Platelet-derived growth factor receptor)^58^, *Wnt4*^59^, and *Cdkn1c* (*p57,* cyclin dependent kinase inhibitor)^55^. In *Ift20^−/−^* endocardium (high KLF4 protein), we observed increased levels of KLF4-activated genes and decreased levels of KLF4-repressed genes, consistent with increased KLF4 expression in the absence of cilia that we previously observed through HCR-WISH and IF analysis **(Fig. 6C)**.

**Figure 6.**
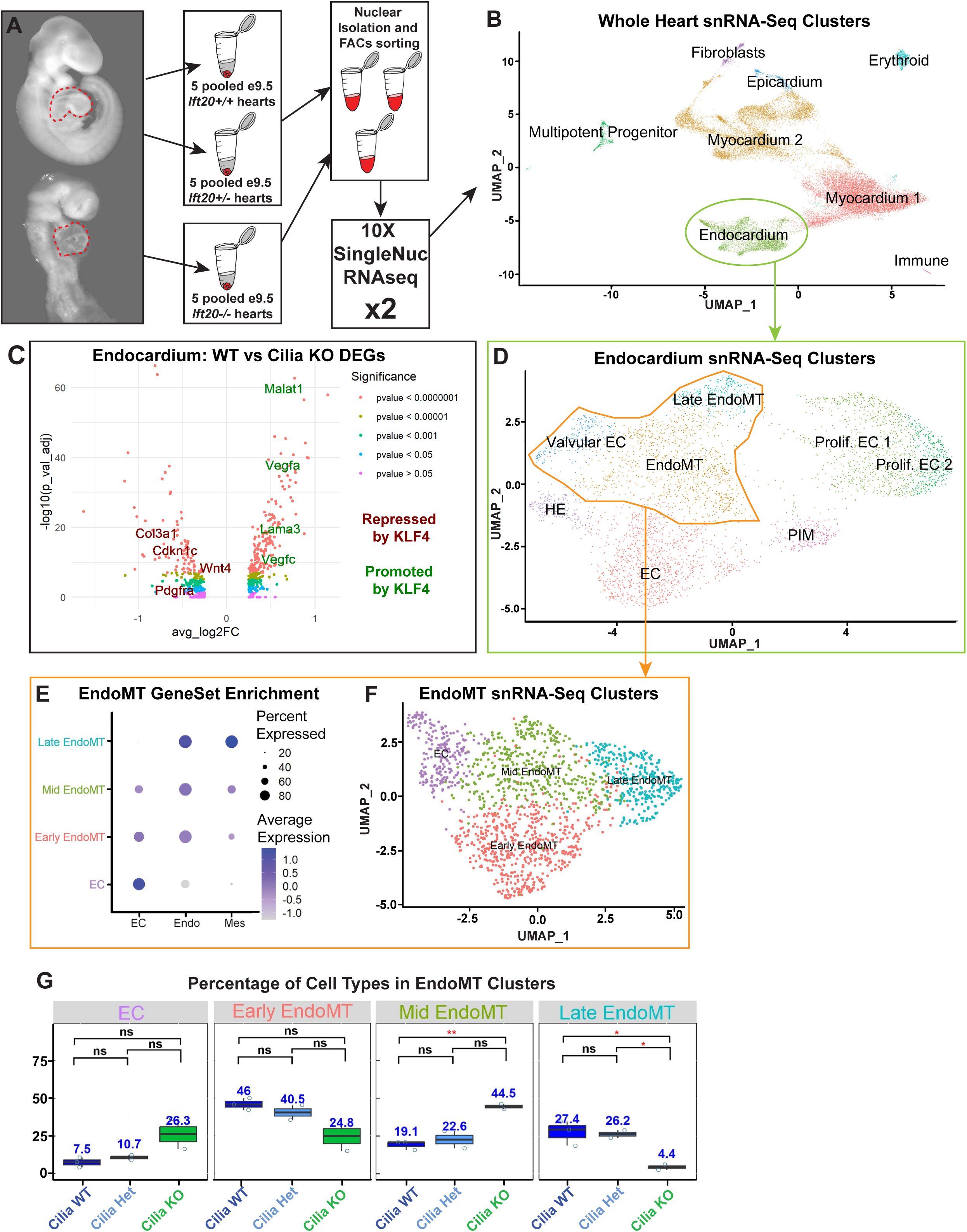
Defects in ciliogenesis block EndoMT. **A)** SingleNuc RNAseq schema. **B)** UMAP plot of e9.5 hearts. **C)** Volcano plot of DEG between wildtype and cilia KO (*Ift20^−/−^*); KLF4 targets are highlighted. **D)** UMAP plot of endocardial cluster from B). **E)** GeneSet Enrichment for EndoMT progression for EndoMT clusters from D). **F)** UMAP plot of EndoMT clusters. **G)** Percent enrichment of EndoMT cell types for wildtype, cilia het (*Ift20^+/−^*) and cilia KO (*Ift20^−/−^*). Statistically significant p-values are in red. *Statistics*: ns (p > 0.05), * (p ≤ 0.05). Data are represented as mean ± SEM. Abbreviations: A-Atrium, AVC-Atrioventricular Canal, EC-Endocardial Cell, HE-Hematoendothelium, LV-Left Ventricle, PIM-Pro-inflammatory, WT-Wildtype.

Utilizing our wildtype EndoMT pseudo-stage markers and GeneSet Enrichment **(Fig. 6E, 6F, S6C)**, we identified the extent to which cilia KO embryos can progress through EndoMT. In *Ift20^−/−^* embryos, the proportion of cells in mid-EndoMT is 2.13x higher than in wildtype and heterozygous littermates, and the proportion of cells in late-EndoMT is 6.1x lower **(Fig. 6G)**. This indicates that cilia KO endocardium has difficulty progressing to the final pseudo-stage of EndoMT, consistent with our observation that cilia KO embryos express FN at the AVC, but fail to cellularize the cushions. This correlates with the timing of *Klf4* downregulation in wildtype hearts, and suggests that *Klf4* downregulation is essential for EndoMT completion. We next wanted to confirm that the observed decreased proportion of late-EndoMT cells in cilia KO mutants was specific to EndoMT progression rather than being caused by non-specific changes in cell number, such as increased apoptosis or reduced proliferation. Pro-apoptotic markers (*Trp53, Casp3, Casp7*) and proportion of G2M cells were not significantly different in the *Ift20^−/−^* EndoMT cluster **(Fig. S6F, S6G)**. This indicates that loss of cilia inhibits EndoMT progression by affecting cell-fate trajectories.

### Known EndoMT molecular pathways are dysregulated in Cilia KO hearts

To understand why cilia KO mutant endocardium fails to progress to late-EndoMT, we identified differentially expressed genes (DEGs) in the EndoMT clusters of *Ift20^−/−^* hearts compared to both wildtype and heterozygous littermates. Upregulated DEGs (avg_LOG2FC of 0.4 or higher, adj_pval < 0.05) in *Ift20^−/−^* hearts were overwhelmingly expressed in quiescent endocardium and mid-EndoMT and included vasculature development genes such as *Tmem100* (Transmembrane protein)*, Slc2a1* (Glucose transporter), and *Ldb2I* (LIM domain binding protein). Downregulated DEGs (avg_LOG2FC of -0.4 or lower, adj_pval < 0.05) in *Ift20^−/−^* hearts were expressed in late-EndoMT and included migration regulatory genes such as *Robo2* (Roundabout Guidance receptor), *Epha7* (Ephrin receptor), and *Erbb4* (Erb-B2 Receptor Tyrosine Kinase) **(Fig. 7A)**. Additionally, *Ift20^−/−^* hearts had decreased expression of mesenchymal (*Sox5, Vcan, Col3a1* and *Ncam1)* and EndoMT markers (*Snai1, Sox9, Twist1, Msx1, Wnt4, Tbx20*), and increased expression of endothelial (*Emcn, Vwf, Cdh5* and *Kdr*) and EndoMT antagonist markers (*Vegfa*) **(Fig. 7B) (Table S3)**. A subset of the DEGs were validated *in vivo* using HCR-FISH **(Fig. 7C, Fig. S7A)**. Gene Set Enrichment Analysis (GSEA) of the top DEGs showed an overwhelming downregulation of Gene Ontology-Biological Process terms related to mesenchymal differentiation, cushion/valve development, and septation formation (30/166) in *Ift20^−/−^* hearts, while terms related to endothelial/vascular identity/integrity (33/140) were upregulated **(Fig. 7D)**. Further, the Smad2/3/4 TGFβ pathway required for EndoMT was perturbed in cilia KOs, as Smad2/3/4 transcriptional targets were down in the *Ift20^−/−^* EndoMT cluster, but not in other cell types such as the epicardium **(Fig. S7B)**. This shows an inability of *Ift20^−/−^* endocardium to adapt a mesenchymal identity during EndoMT.

**Figure 7.**
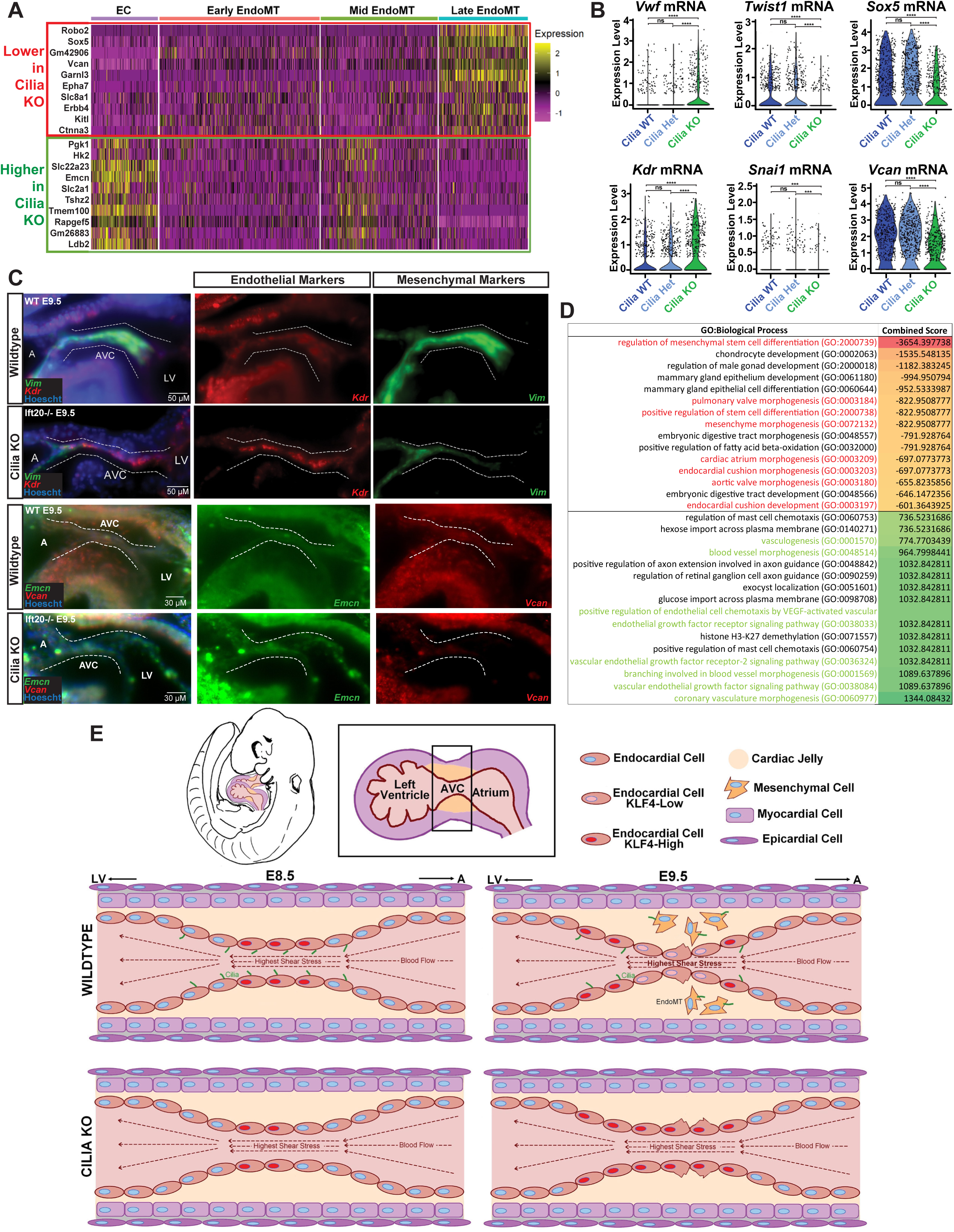
Known EndoMT molecular pathways are dysregulated in Cilia KO hearts. **A)** Top DEGs between cilia KO (*Ift20−/−)* and wildtype hearts over EndoMT clusters. **B)** Endocardial, EndoMT and mesenchymal gene expression in wildtype, cilia het (*Ift20^+/−^*) and cilia KO (*Ift20^−/−^*). **C)** Whole mount HCR-FISH for *Kdr* (red), *Vim* (green), *Emcn* (green) and *Vcan* (red) in wildtype and cilia KO (*Ift20^−/−^*) e9.5 AVCs. Nuclei are shown in blue (Hoescht); the AVC is outlined in white. **D)** Gene Ontology-Biological Processes lower and higher in cilia KO (*Ift20^−/−^*). Cushion relevant terms are highlighted. **E)** <u>Summary graphic:</u> wildtype hearts selectively lose cilia in areas of highest shear stress, allowing KLF4 reduction and subsequent EndoMT/cushion cellularization. Without cilia present, KLF4 is unable to be regionally turned off and EndoMT/cushion cellularization cannot progress. *Statistics*: ns (p > 0.05), *** (p ≤ 0.001), **** (p ≤ 0.0001). Abbreviations: A-Atrium, AVC-Atrioventricular Canal, EC-Endocardial Cell, LV-Left Ventricle, WT-Wildtype.

To determine transcriptional changes driving abnormal transition between mid- and late-EndoMT in *Ift20^−/−^* hearts we identified genes that *Ift20^−/−^* hearts were unable to turn on or off between each stage of EndoMT as compared to wildtype hearts **(Table S4)**. GSEA of these gene networks showed that *Ift20^−/−^* hearts had an excess of cell-cell/adherens junctions (#1 GO:CC, Adj. p-value: 2.589e^-8^) and deficiencies in collagen synthesis (#1 GO:CC, Adj. p-value: 4.88e^-6^) between mid- and late-EndoMT, which suggests *Ift20^−/−^* endocardial cells on the AVC lumen lack both the ability to separate from each other and to physically migrate into the cushion, providing a possible explanation for the complete lack of cellularization in cilia KO cushions. Since these pathways are predominately controlled by well-established networks inhibited by KLF4^32^, we hypothesized that these transcriptional changes may be due to abnormally sustained KLF4 expression into late-EndoMT in *Ift20^−/−^* hearts. Accordingly, we identified KLF4 as a significant TRRUST 2019 transcription factor (the current largest database of literature-curated transcription factor-target interactions^60^) (Adj. p-value: 0.016) for DEGs higher in the *Ift20^−/−^* EndoMT cluster (Adj. p-value: 0.17 for DEGs lower in *Ift20^−/−^*). Additionally, motif enrichment analysis^61^ found that DEGs in the *Ift20^−/−^* EndoMT cluster disproportionally possessed KLF4 binding motifs (enrichment ratio of 5.27 for higher DEGs (p=value: 2E-09), 1.82 for lower DEGs (p-value: 0.003) compared to shuffled control sequences). This is consistent with a model in which the EndoMT defects observed in *Ift20^−/−^* hearts are driven by sustained endocardial KLF4 expression.

## Discussion

Our data is consistent with a model whereby intracardiac primary cilia are used transiently to sense blood flow and spatially control EndoMT in early cardiac cushions through regulation of endocardial KLF4. We found that e8.5-10 mouse embryo endocardial primary cilia extending into the cardiac lumen become spatially regionalized to the junctions between the ventricle and the AVC and OFT cushions as the cushions develop in a blood-flow dependent manner. Variants affecting KLF4 contribute to human valve defects associated with HLHS, and KLF4 expression spatially correlates with endocardial ciliation during cushion development. Mouse embryos constitutively lacking cilia express significantly higher amounts of KLF4, with a loss of KLF4 spatial patterning in the AVC and an inability to cellularize their cushions. These phenotypes were independent of the LR defects associated with loss of cilia. Comparison of snRNA-seq datasets of micro-dissected wildtype and cilia knockout mouse hearts demonstrated that cilia KO endocardium was unable to progress from mid- to late-EndoMT due to mis-regulated KLF4 targets, corresponding to when *Klf4* becomes significantly reduced in wildtype EndoMT.

### Cilia as intracardiac mechanosensors

Cardiac contraction and blood flow initiate at the same time as major events in cardiogenesis: looping, chamber formation and valve formation. Much like a self-assembling machine, cardiac function is required for cardiac development, and resulting hemodynamic forces provide a variety of mechanical signals that need to be sensed and integrated for cardiogenesis to progress^8^. Forces resulting from contraction (circumferential stress), blood flow (shear stress) or a combination of both have been proposed as potential drivers of valve development^11,31,62–64^, however, the cellular structure(s) that function as mechanosensors remain elusive.

Work utilizing particle image velocity analysis, ultrasound scanning, and *in silico* models point to blood flow-dependent wall shear stress (laminar and oscillatory) as the predominant force in the AVC and OFT^7,24,34^. Genes known to be shear-stress specific (or inhibited by cyclic strain^65^), such as *klf2a*, become increasingly restricted to the AVC following the onset of blood flow in zebrafish hearts^31,66^. Consistent with this, we show *Klf2* and *Klf4* mRNA (also specific to shear stress^67^) specifically expressed in mouse ECCs. Therefore, endocardial mechanosensors capable of responding to shear forces, such as Notch, TRP/Piezo1 channels, or primary cilia, are candidate mechanosensors regulating *Klf2/4* expression and cushion development.

At the organelle level, primary cilia are excellent candidate shear stress mechanosensors due to their architecture, the vast hub of signaling networks that localize to them, and their ability to be endogenously removed. At the start of contraction, mouse embryonic hearts experience shear stress averaging <2-5 dyn/cm^2 68^, with increments as low as ∼1.2 dyn/cm^2^ between the center and edge of the AVC as development progresses^35^. This is smaller than the shear sensitivity of other known shear-sensitive endothelial mechanosensors, such as NOTCH1 (>10 dyn/cm^2 69^) or ion channels/Piezo1 (5-10 dyn/cm^2 70–72^). In contrast, primary cilia can detect shear forces as low as 0.2 dyn/cm^2 73^, and elicit a calcium signal in response to 0.5 dyn/cm^2 74^.

Beyond their ability to sense extremely low forces, intracardiac primary cilia are also biased toward areas of lower shear stress, adding an additional layer of control in determining genetic outcomes in accordance with stress levels^15,75^. Primary cilia disassemble at ∼15 dyn/cm^2 21^, and the center of the AVC during late cushion formation can experience an average of ∼30 dyn/cm^2^ while the edges experience an average of ∼10 dyn/cm^2 31,34^, though these results have not been validated in *in vivo* mouse hearts. In line with this, we found endocardial primary cilia were maintained on the edges of the AVC and OFT and were reduced in the center, showing correlation between shear stress and ciliation. This pattern was absent in mouse mutants lacking contractility, demonstrating a need for blood flow to establish endocardial ciliary regionalization.

For cilia to act as cardiac mechanosensors, they need to not only sense but translate force into gene regulation. Previously, primary cilia were found to sensitize endocardial cells for mechanosensitive gene expression^76^. Further, *ift88* KD (cilia KD) causes endocardial misexpression of mechanosensitive gene targets in zebrafish AVCs^23^. We now establish a correlation between endocardial ciliation and mechanosensitive gene expression and show that a constitutive loss of primary cilia perturbs these mechanosensitive targets, supporting a function for primary cilia as mediators of shear forces that are transient by design, directing mechanosensitive gene expression through selective, regional decilliation.

### Intracardiac Primary Cilia guide cushion development independently of LR Asymmetry

Since cilia mechanosensation at the LRO is essential to establishing LR asymmetry and direction of the heart loop, it has been unclear whether the wide range of cardiac defects observed in mice and humans with abnormal cilia are due to a primary requirement for intracardiac cilia or a secondary manifestation of abnormal blood flow due to defective heart looping. Although altered fluid flow within the heart through manipulation of heart looping has been previously shown to affect subsequent valve formation in zebrafish^77^, work by Burnicka-Turek et al. (2016)^78^ found a high prevalence of variants in ciliary signaling genes in patients with atrioventricular septal defects (AVSD) irrespective of LR patterning defects. To distinguish between a requirement for ciliary motility/LR patterning and ciliary sensation in ECC development, we utilized the *Dnah11^−/−^* mouse mutant which lacks ciliary motility but has intact ciliogenesis and mechanosensation. We found that *Dnah11^−/−^* mouse hearts, even with the same randomized heart looping as full cilia KO, were indistinguishable from wildtype with respect to the number or location of KLF4 positive cells, endocardial cilia, or cushion cellularization. This data is consistent with a model whereby the increase and loss of regionalization of KLF4 observed in cilia KO is due to a specific role of intracardiac primary cilia during cushion development versus upstream functions in LR asymmetry.

### Intracardiac cilia regulate endocardial KLF4 expression

Though various endothelial mechanosensors for KLF4, such as TRP/Piezo1 channels, Notch1 and integrins, have been described^32,79^, we propose that primary cilia are KLF4 mechanosensors during ECC formation. We observed a relationship between primary cilia and KLF4 expression during ECC formation and predicted that constitutive loss of cilia through knockout of *Ift20* or *Kif3a* would also cause KLF4 loss. However, KLF4 only decreased in cilia KO during early cardiogenesis, when there is blood flow, but shear stress is still relatively low. As contractility develops and shear stress increases, we found that paradoxically cilia KO hearts accumulate KLF4 and have a loss of the organized spatial patterning of KLF4 expression along the developing AVC. These conflicting results provide a potential mechanism by which cilia can interpret a gradient of mechanical forces and translate them into developmentally relevant signaling: first in sensing low shear stress and turning KLF4 on, and second in turning KLF4 off in response to deciliation mediated by high shear stress.

We show that at early timepoints constitutive cilia loss inhibits *Klf4* upregulation, while at later timepoints (when shear stress is increased) active loss of cilia correlates to reduced KLF4. Previous work by Goetz et al. (2014)^80^ demonstrated primary cilia amplify mechanical forces in low flow settings, suggesting a ciliary role in triggering gene expression through cytoskeletal deformation prior to shear forces strong enough to trigger deformation alone. This aligns with what we know of shear forces on early endocardium (<2-5 dyn/cm^2^) and cytoskeletal deformation resulting in intracellular calcium signals (10 dyn/cm^2^)^81^. Work by Zheng et al. (2022)^33^ implicated intracellular calcium signaling in the CaMKII-mediated Mekk3 activation upstream of *Klf4* expression. In this model, primary cilia are needed for *Klf4* induction in low flow to activate CaMKII, but not in high flow.

Unexpectedly, KLF4 was increased at later timepoints in cilia KOs. TGFβ signaling is known to inhibit KLF4 *in vivo* and *in vitro* ^82–84^. TGFBR1/2 localize to the ciliary axoneme, and constitutive loss of cilia results in decreased Smad2/3/4 signaling^85–87^, which we confirmed in the EndoMT cluster of cilia KO hearts. Therefore, a mechanism by which constitutive loss of cilia leads to lower TGFβ-dependent inhibition of KLF4 in cilia KO seems plausible. It is also interesting to consider the mechanism(s) leading to shear-stress induced deciliation, such as HDAC-mediated resorption or whole-cilium shedding through p60 katanin activity^88^. Additionally, there is a well-defined role for calcium in both initiating deciliation and ultimately deciding on the method of deciliation^44,89–91^. However, the question remains what distinguishes endogenous loss of cilia due to shear stress, such as in wildtype hearts, from never possessing a cilium at all in a cilia KO in the resulting KLF4 expression? One possible explanation is in work by Rozycki et al. (2014)^92^ which found a necessity for primary cilia in epithelial-myofibroblast transition through TGFβ signaling that was accelerated upon ciliary loss. Similarly, selective cilia loss in high shear areas regulates BMP signaling during vascular regression^93^. In both examples, primary cilia are needed initially to activate transition, and then their loss is needed to complete it. Our results suggest that cilia presence and subsequent loss are required to regionally control KLF4 expression through TGFβ. Mutants that never possessed cilia are unable to activate TGFβ at all, resulting in overexpression and mislocalization of KLF4.

### KLF4 inhibits EndoMT progression during cushion formation

Whether KLF4 activates or inhibits cushion EndoMT is still up for debate^40,41^. Given these inconsistencies, it is important to not only understand how KLF4 is regulated during cushion formation, but also how it impacts EndoMT progression. To explore the functional significance of KLF4 regionalization on cushion formation, we investigated the spatial localization of EndoMT, where we found both genetic and morphological EndoMT markers first appear in the center of the AVC where KLF4 expression was the lowest. This suggested to us that KLF4 could be playing an inhibitory role in cushion EndoMT. Cementing this idea, we examined the correlation between *Klf4* expression and multiple essential EndoMT genes, including *Snai1,* in each case finding a negative correlation. This is consistent with previous work that found a KLF4-dependent inhibition of mesenchymal transition through suppression of *Snai1*^47^, suggesting a requirement for *Klf4* downregulation in EndoMT progression.

This interpretation is consistent with accumulation of KLF4 in the cilia KO leading to an EndoMT defect. We confirmed this KLF4 abundance transcriptionally in cilia KO by seeing upregulated KLF4 targets. Functionally, KLF4 not only inhibits multiple essential EndoMT genes, but it can promote antagonistic EndoMT factors such as VEGF signaling^56,94^. In line with this, we saw a large increase in VEGF pathway members *Vegfc* and *Vegfa* in cilia KO hearts, as well as multiple GO:BP terms related to VEGF signaling. Collectively, these results point to abnormal KLF4 overexpression stalling EndoMT progression in cilia KO hearts.

### “Churning” model for early valve development

Taken together, our data puts forward a model by which primary cilia spatially control cushion growth through selective regulation of mechanosensitive factors. In this model, cilia dependent KLF4 expression is used to maintain endocardial quiescence, as has been seen previously^41^, thus providing an endocardial pool for replenishing differentiated luminal cells. Ciliated endocardium becomes *Klf4* positive following the onset of blood flow, and then selectively deciliates in the high shear stress center of the AVC leading to reduced KLF4 **(Fig. 7E)**. No longer being inhibited by KLF4, cells in the center can undergo EndoMT, but must be replaced by neighboring luminal endocardial cells as they migrate away from the lumen into the cardiac jelly. These neighboring cells will then be subjected to the higher shear stress in the center of the AVC, causing their own deciliation and initiating the process over again. Two important mechanisms come from this schema, which we call the “churning” model of ECC formation. First, it provides a method by which a mass quantity of endocardial cells can migrate to and cellularize the cushion in a relatively brief period, and is supported by previous findings of ventricular ECs contributing to the AVC cushion in zebrafish^66,95^. Second, it provides a mechanism by which the spatial identity of the endocardial cushion can be maintained through a positive mechanosensitive feedback loop during development: high shear stress leads to regional deciliation, downregulating KLF4 and promoting EndoMT, thus increasing the thickness of the AVC. This will then only act to make the lumen smaller and increase shear stress, as the same amount of blood must now flow through a smaller orifice. In this way, deciliation is used as a tool for spatial identity of the cushion as it grows **(Fig. 7E)**.

### Limitations

We identify primary cilia as a link connecting blood flow and transcriptional regulation of cardiogenesis. Paradoxically, constitutive loss of cilia seems to inhibit the wildtype functionality of selective deciliation. Therefore, the impact of forced continued ciliation would be important to explore in future studies, though the means to do this may impact cell proliferation and homeostasis. While we uncovered a necessary function for primary cilia in KLF4 regulation during cushion formation, the exact mechanism by which cilia translate blood flow into KLF4 expression still needs to be addressed, especially since many pathways, such as TGFβ, have feedback loops with KLF4^96,97^. Finally, the temporal dynamics of our model, such as our proposed churning mechanism, are difficult to achieve in murine embryos. Further studies using other models could further elucidate these aspects.

## Acknowledgements

We thank the Yale Center for Genome Analysis for DNA sequencing, the Yale Imaging Facility on Science Hill for lightsheet microscopy and the Yale Center for Cellular and Molecular Imaging for confocal microscopy and analysis workstations. We thank Jeffrey Drozd and Thomas Nuzzo for mouse support. We thank Daniel DeLaughter for snRNA-seq alignment. We also thank Dr. Zhaoxia Sun for sharing transgenic zebrafish with us, and Drs David Breslow, Zhaoxia Sun, Shiaulou Yuan, and Mark Pownall for critical review of the manuscript. This work was supported by the NHLBI Ruth L. Kirschstein National Research Service Award (NRSA-F31) to K.B. (F31HL158091) and NHLBI UO1HL098162-12 and R35HL145249 to M.B.

## Author Contributions

Conceptualization: K.B. and M.B.; Formal Analysis, Data Curation, and Visualization: K.B.; Investigation: K.B., F.L., J.G., J.S., M.B.; Writing – original draft: K.B. and M.B.; Supervision, Project administration, Funding acquisition: M.B.

## STAR Methods

### Resource Availability

#### Lead Contact

Information and resource requests should be directed to the lead contact, Martina Brueckner.

#### Material Availability

This study did not generate new unique reagents.

#### Data and Code Availability

- Single-nuclei RNA-seq data reported in this paper have been deposited in GEO: GSE252341 and are publicly available. The single cell RNA-seq data from De Soysa et al. (2019)^28^ is available in GEO under accession number GSE126128 and through the UCSC Cell Browser at https://mouse-cardiac.cells.ucsc.edu. Accession numbers for both datasets are listed in the key resources table.
- This study did not generate original code. See section ‘Single Nucleus RNAseq’ for published packages used in this paper.
- Any additional information required to reanalyze the data reported in this paper is available upon request from the lead contact.

### Experimental Model Details

*<u>Mice:</u> Dnah11^−/−^*(*lrd^GFPΔneo/GFPΔneo^*)^51^ embryos were obtained from crossing a homozygous female and male. *Ift20^−/−^* ^98^, *Kif3a^−/−^* ^49^, and *Ncx1^−/−^* ^39^ embryos were obtained from crossing a heterozygous female and male. We generated *Ift20^+/−^/Ncx1^+/−^* mice which were viable and presented no phenotype, and crossed a female and male to obtain *Ift20^−/−^/Ncx1^−/−^* embryos. All the lines used in this study are maintained on the C57BL/6 background and fed breeder diets. Timed matings were conducted for each strain such that noon on the day of vaginal plug detection was considered e0.5. Plugged females were weighed prior to the day of dissection to validate pregnancy. Embryonic staging was determined by somite count as described^99^ (6-8 somites for e8, 11-14 somites for e8.5, 15-19 somites for e9, 20-25 somites for e9.5, and 26-29 somites for e10). Both sexes were pooled for each embryonic experiment, and randomized littermates were used as controls. This research complies with all ethical regulations; mouse experiments were performed in a manner approved by the Yale University Institutional Animal Care and Use Committee.

*<u>Zebrafish</u>*: Tg(*Sco:eGFP;Kdrl:mCherry;cmlc2:eGFP*)^100,101^ zebrafish were obtained from crossing homozygous transgenic females and males. Standard protocols were used for maintaining zebrafish colonies^102^. Embryos were obtained through natural spawning. All zebrafish experiments were conducted according to Yale Animal Resources Center and Institutional Animal Care and Use Committee guidelines.

### Histology

Embryos were dissected in cold 1x Phosphate Buffered Solution (PBS). A sample of the allantois was taken for genotyping. Embryonic somites were then counted and embryos were fixed in 4% paraformaldehyde in 1xPBS overnight (16 hours) at 4°C. Embryos were then prepared for cryosectioning as described^103^. Briefly, embryos were saturated with 30% sucrose and mounted in 10x10x5mm cryomolds filled with O.C.T. medium (Tissue-Tek) and frozen in chilled 100% ethanol. Following freezing, blocks were sliced into 12 µm sections with a Leica CM3050 S Research Cryostat and mounted on Superfrost Plus slides (Thermo Fisher). Defrosted slides were washed with 1x PBS before being counterstained with Nuclear Fast Red Solution (Ricca) for 5 minutes. They were then rinsed with water and dehydrated prior to mounting with CytoSeal (Thermo Fisher) on a coverslip. Slides were then imaged on a Zeiss Axiovert microscope, using an Axiocam 1077 driven by Axiovision software.

### Whole mount and cryosection *in situ* hybridization

*<u>Colorimetric:</u>* Digoxigenin-labeled antisense probes were prepared using SP6 and T7 RNA polymerase (NEB) off linearized expression plasmids. pCX-Klf2 and pCX-Klf4 were gifts from Barak Cohen (Addgene plasmid #s 66655 and 66656, respectively). For linearization and transcription, pCX-Klf2 used EcoRV/SP6 and pCX-Klf4 used HindIII/T7. Embryos were stained as described^104^. Briefly, embryos were dehydrated in a methanol series and rehydrated, permeabilzed, and incubated overnight in hybridization buffer with probe (∼ 1 ug/ml) at 68°C. The next day, embryos went through a series of washes into 1x MABTw, blocked for 2 hours in 2% Blocking Reagent (Roche) in MABTw with 20% NGS, and incubated overnight in anti-DIG antibody (1:2000, Roche) at 4°C. The next day, embryos were washed in MABTw, followed by NTMT (pH 9.5), and then developed in BM-Purple solution (Roche) with 2mM levamisole and 1% Tween. Once the reaction was complete, development was stopped with PBSTw. Whole embryos were mounted in 1xPBS and imaged on a Nikon SMZ 745T dissection scope equipped with an Excelis HDS HD camera and monitor system. Embryos were then prepared for cryosectioning and Nuclear Fast Red (Ricca) counterstaining as described^103^. Slides were imaged on a Zeiss Axiovert microscope, using an Axiocam 1077 driven by Axiovision software.

*<u>Hybridization Chain Reaction (HCR)-FISH:</u>* HCR oligo probes were synthesized for *Klf2, Klf4, Vim, Kdr, Vcan,* and *Emcn* by Molecular Instruments^105^. The standard company protocol given for whole-mount e9.5 mouse embryos was performed, with the following change: ProK digestion (step 12) for 10 minutes. Following the last 1xSSCTw wash, the embryos were incubated for 20 minutes with 1:2,000 Hoescht 333342 before another 1xSSCTw wash. Embryos were stored protected from light in 1xSSCTw at 4°C until being imaged on a Zeiss Z.1 Lightsheet. Imaged embryos were recollected and frozen in O.C.T. medium (Tissue-Tek) for later cryosectioning as described^103^. Defrosted slides were rinsed with 1xPBS and mounted with ProLong Gold antifade reagent (Invitrogen, P36935) on coverslips. Sections were examined with a Zeiss Axiovert microscope equipped with Apotome imaging.

### Immunofluorescence

*<u>Whole mount:</u>* Embryos were dissected in cold 1x Phosphate Buffered Solution (PBS). A sample of the allantois was taken for genotyping. Embryonic somites were then counted and embryos were fixed in 4% paraformaldehyde in 1xPBS overnight (16 hours) at 4°C. The next day, embryos were briefly washed with 1xPBS and then put through an ethanol dehydration and rehydration series (25%, 50%, 75%, 100% at -20°C for 30 minutes, 75% back at room temperature, 50%, 25%, 0%). Embryos were then further permeabilized with 2 washes of 1xPBS with 1% Triton-X (1xPBSTx), followed by blocking for >2 hours (3% BSA in 1xPBSTx). Embryos were then incubated in block with primary antibody overnight at 4°C. The next day, embryos were washed 4 x 15 minutes with 1xPBSTx before incubating with secondary antibody in block for 2 hours. Embryos were then rinsed 3 x 15 minutes in 1xPBS and incubated for 20 minutes with 1:2,000 Hoescht 33342 (Thermo Fisher) before being rinsed again. Whole embryos were kept in 1xPBS at 4°C protected from light until being imaged on a Zeiss Z.1 Lightsheet and Zeiss 880 Confocal Microscope.

*<u>Cryosections:</u>* Embryos were embedded in O.C.T. medium (Tissue-Tek) and sectioned as described^103^. Defrosted slides were then put through the same protocol as whole mount immunofluorescsence. After the last 1xPBS rinse, slides were mounted with ProLong Gold antifade reagent (Invitrogen, P36935) on coverslips. Sections were examined with a Zeiss Axiovert microscope equipped with Apotome imaging.

*Antibodies:* Mouse anti-Arl13B (1:200 Neuromab), Rabbit anti-Arl13b (1:200 ProteinTech), Goat anti-KLF4 polyclonal (1:100 R&D Systems), Rat anti-CD31 (1:200 BD Biosciences), Rabbit anti-FN (1:200 Sigma Aldrich), Alexa 488 anti-mouse (1:500, Invitrogen), Alexa 488 anti-rabbit (1:500, Invitrogen), Alexa 594 anti-goat (1:500, Invitrogen), Alexa 647 anti-rat (1:500, Invitrogen), Hoescht 33342 (1:2000, Thermo Fisher), Sheep anti-DIG-AP (1:2000, Roche).

### Lightsheet Microscopy

*<u>Fixed:</u>* Embryos were embedded in Zeiss glass capillaries suited for their size (blue for e9.5-e10 mice, black for e8-e9 mice and zebrafish) in 1% purified low-melt agarose (Thermo Fisher). The chamber was filled with filtered 1xPBS prior to imaging on a Zeiss Z.1 Lightsheet.

*<u>Live:</u>* 1x 1-phenyl 2-thiourea (PTU, Sigma) was added to zebrafish media at 24 hours post fertilization to block pigment formation. Zebrafish embryos were then screened for fluorescent signal using a Leica fluorescence stereomicroscope. Prior to embedding in glass capillaries, tricaine (Sigma) was added to embryo media to immobilize them at 168 ug/ml. The chamber was filled with embryo media containing tricaine and allowed to heat to 26°C prior to imaging. The embryos were then embedded in glass capillaries and imaged as described^106^. Briefly, 200 heartbeats were recorded every 3 uM of a z-stack encompassing the entire heart at the maximum frame rate. Embryos were imaged every 12 hours between 36 and 96 hpf.

### Imaging Analysis

Cell and cilia quantification was done using Cellpose^107^, Imaris Software (RRID:SCR_007370) and Fiji^108^. 4D analysis of live zebrafish utilized Fiji to create hyperstacks for one cardiac cycle and FluoRender^109^ for rendering and figure/video creation. HCR quantification was done using Fiji’s “Measure” and “Plot Profile” functions. To compare replicates with various percentage lengths, a Matlab script was used to autofill points between experimentally provided points.

### Single Nucleus RNAseq

*<u>Single Nucleus Collection, Sequencing and Alignment:</u>* e9.5 embryos were dissected in cold 1x Phosphate Buffered Solution (PBS) with diethyl pyrocarbonate (DEPC). A sample of the allantois was taken for genotyping. Whole hearts were removed, allowed to self-perfuse and flash frozen in liquid nitrogen. These were kept frozen in liquid nitrogen until processing. Pools of 5 embryonic hearts were used per genotype, with two replicates for *Ift20−/−* and *Ift20+/−* genotypes, and three replicates for *Ift20+/+*. Samples were prepped as described^110,111^. Briefly, RNA quality was checked using the Agilent TapeStation with an RNA integrity number (RIN) cutoff of ≥ 5. The sample was then homogenized in buffer (250 mM sucrose, 25 mM KCL, 5 mM MgCl_2_, 10 mM Tris-HCL, 1 mM dithiothreitol (DTT), 1x protease inhibitor) with a TissueLyser II (Qiagen) and filtered through a 40-uM strainer (Corning). The nuclei were then centrifuged at 1200g for 5 minutes at 4°C and resuspended in storage buffer (1x PBS, 4% BSA, 0.2 U ul-1 Protector RnaseIn). Nuclei were FACS sorted using NucBlue Live ReadyProbes Reagent (Thermo Ficher) and assessed for morphology, size, and concentration with a Countess (Thermo Fisher). Intact nuclei were fractionated with the 10x Genomics Chromium iX according to manufacturer’s protocol. 10x Genomics Chromium Single Cell Reagent Kit (v3.1) was used to generate gene expression libraries. Libraries were sequenced with a NovaSeq 6000 (Illumina). Reads were aligned to the mm10 mouse genome using Cell Ranger (v3.0.1).

*<u>Single Nucleus Seurat Object Creation:</u>* Doublet prediction was assigned with Scrublet v0.2.3^112^ in Python v3.8.8^113^ with an expected_doublet_rate of 0.06, min_counts of 2, min_cells of 3, min_gene_variability_pctl of 85 and n_prin_comps of 30. Data files were read into R v4.1.2^114^ with Seurat v4.3.0^115–118^ and subset based on Scrublet Score (WT1-3: 0.2, 0.2, 0.15, Het1-2: 0.3, 0.3, Mut1-2: 0.3, 0.2). These 7 Seurat objects were then merged, log normalized (1e4) and examined for variable features (“vst”, 2500 features). The resulting Seurat object was then subset based on number of counts (>300, <15000), number of features (>300, <5000), and percent mitochondrial (< 0.02) before being regressed for mitochondrial and ribosomal genes. Integration of cells was run with Harmony v0.1.1^119^ in R. The resulting object contained 29,313 single nuclei with 27,998 features over 2 replicates (Nuclei counts: *Ift20^+/+^* 9,882; *Ift20^+/−^* 14,825; *Ift20^−/−^* 6,116).

*<u>Single Nucleus Analysis:</u>* Cell cycle analysis was done as described (Tirosh et al, 2015). The AddModuleScore function in Seurat was used for Geneset Enrichment. Gene Ontology Analysis for Biological Processes (BP), Cellular Compartments (CC) and TRRUST 2019^60^ enrichment was done with Enrichr^120,121^. Statistics for boxplots were calculated with rstatix v0.7.2.999 using a t-test with Bonferroni correction. Statistics for violin plots were calculated with ggpubr v0.6.0 using a student’s t-test. Motif enrichment analysis was performed with MEME Suite v5.5.5 Simple Enrichment Analysis^61^. P values are represented as follows: ns (p > 0.05), * (p ≤ 0.05), ** (p ≤ 0.01), *** (p ≤ 0.001), **** (p ≤ 0.0001).

*<u>Single Cell Analysis from DeSoysa et al. (2019):</u>* The expression matrix and meta files were downloaded for the “Early Mouse Cardiogenesis” and “Endocardial/Endothelial” subcluster from UCSC Cell Browser^28,122^. The AddModuleScore function in Seurat was used for Geneset Enrichment. Statistics for violin plots were calculated with ggpubr v0.6.0 using a student’s t-test. P values are represented as follows: ns (p > 0.05), * (p ≤ 0.05), ** (p ≤ 0.01), *** (p ≤ 0.001), **** (p ≤ 0.0001).

### Statistics

All statistical analysis was performed in GraphPad Prism version 9.2.0.332 for Windows (San Diego, CA USA, www.graphpad.com). Statistical details, such as number of animals (n), can be found in figure legends. Graphs were created using the mean and SEM for error bars. Simple linear regression was used to measure R^sq^ values of XY graphs for best fit. A two-tailed, one sample t-test was performed to measure reproducibility between biological replicates, and a two-tailed unpaired t-test with Welch’s correction was performed to measure variance between experimental groups. Statistics for snRNA-seq boxplots were calculated with rstatix v0.7.2.999 using a t-test with Bonferroni correction. Statistics for snRNA-seq and scRNA-seq violin plots were calculated with ggpubr v0.6.0 using a student’s t-test. P values are represented as follows: ns (p > 0.05), * (p ≤ 0.05), ** (p ≤ 0.01), *** (p ≤ 0.001), **** (p ≤ 0.0001).

---

**Figure S1.**
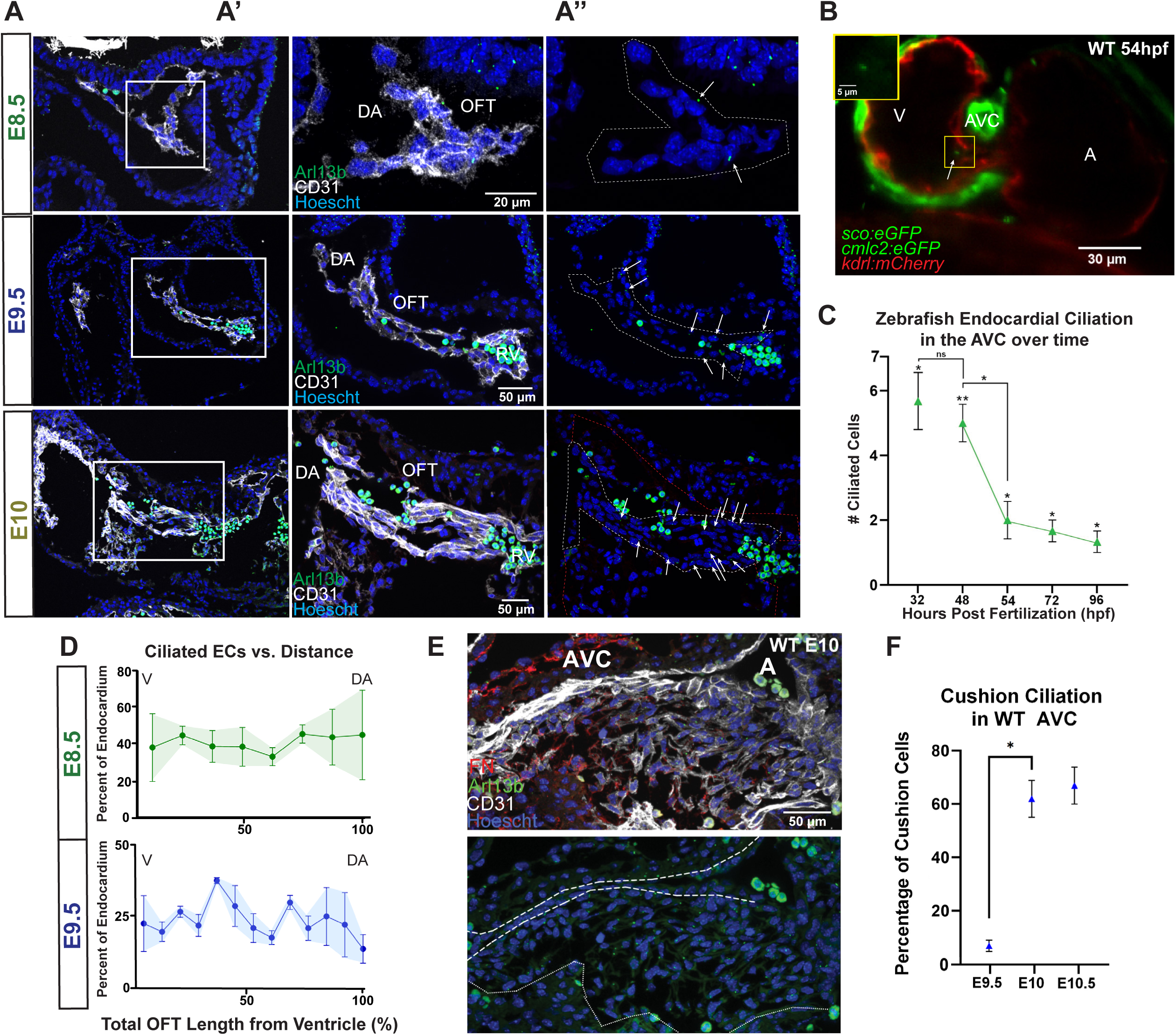
Endocardial ciliation of the ECCs is spatiotemporally dynamic during cushion EndoMT and is conserved in zebrafish. **A)** Immunofluorescence on mouse embryo sections for cilia (ARL13B, green) on endocardial cells (CD31, white) of the OFT over cushion development (e8.5, e9.5, e10). Nuclei are shown in blue (Hoescht). Close ups **A’)** with and **A”)** without CD31; white arrows indicate cilia; luminal endocardial cells outlined with white and whole OFT (including cushion mesenchyme) outlined in red. **B)** Still of live 54hpf tg(Arl13b:eGPF;Kdrl (Flk-1):mCherry;Cmlc2:eGFP) zebrafish heart showing cilia and myocardium in green and endocardium in red. Closeup of cilia provided in yellow box. **C)** Zebrafish endocardial ciliation over time in the atrioventricular canal (n=3). **D)** Ciliated endocardial cells as percentage of all endocardial cells versus distance in the mouse OFT at e8.5 (n=2) and e9.5 (n=3). Distance runs left to right, Ventricle to Dorsal Aorta. **E)** Immunofluorescence on e10 mouse embryo sections for cilia (ARL13B, green), endocardial cells (CD31, white) and mesenchyme (FN, red) of the AVC. Nuclei are shown in blue (Hoescht). Nuclei and cilia are shown alone with the luminal endocardial cells outlined with dashed white and whole AVC (including cushion mesenchyme) outlined in dotted white. **F)** Number of ciliated cushion cells as a percentage of total cushion cells in the AVC over time (e9.5 (n=3), e10 (n=3), e10.5 (n=3)). Statistics: ns (p > 0.05), * (p ≤ 0.05), ** (p ≤ 0.01). Data are represented as mean ± SEM. Abbreviations: A-Atrium, AVC-Atrioventricular Canal, EC-Endocardial Cell, V-Ventricle, OFT-Outflow Tract, WT-Wildtype.

**Figure S2.**
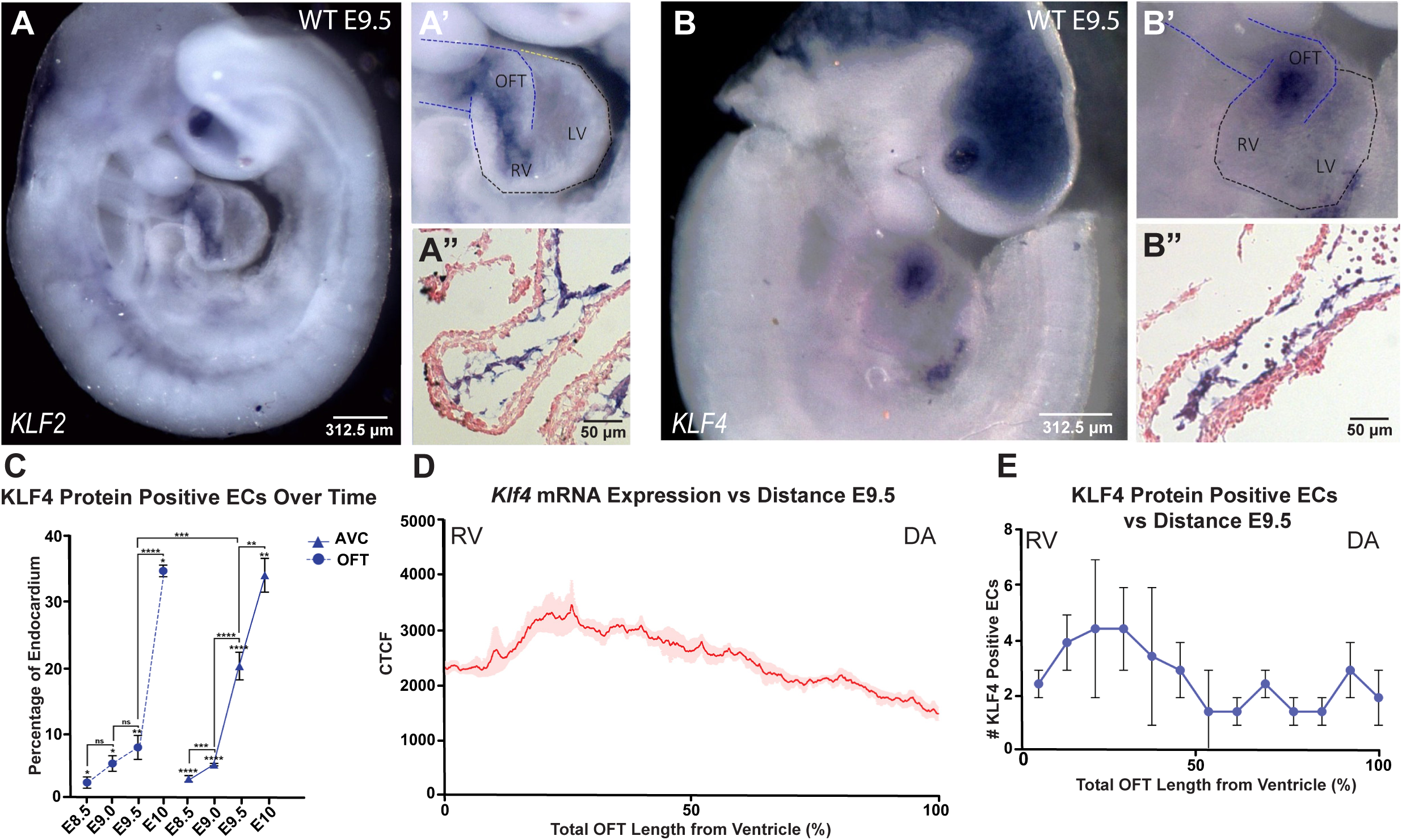
The mechanosensitive transcription factors KLF2 and KLF4 are dynamically expressed in mouse endocardium during cushion EndoMT. Whole mount ISH of e9.5 mouse embryos for A) *Klf2* mRNA and B) *Klf4* mRNA with closeup **(A’, B’)** and cryosection **(A”, B”)** of the OFT. **C)** KLF4 protein positive endocardial cells over time as percentage of all endocardial cells in the OFT (e8.5 (n=11), e9 (n=3), e9.5 (n=6), e10 (n=3)) and AVC (e8.5 (n=14), e9 (n=6), e9.5 (n=8), e10 (n=3)). **D)** mRNA expression as measured by Grey_Value over distance for *Klf4* (n=3) in the e9.5 OFT. **E)** KLF4 protein positive endocardial cells versus distance in the e9.5 OFT (n=3). For D) and E), distance runs left to right, Atrium to Left Ventricle. *Statistics*: ns (p > 0.05), * (p ≤ 0.05), ** (p ≤ 0.01), **** (p ≤ 0.0001). Data are represented as mean ± SEM. Abbreviations: DA-Dorsal Aorta, EC-Endocardial Cell, LV-Left Ventricle, OFT-Outflow Tract, RV-Right Ventricle, WT-Wildtype.

**Figure S3.**
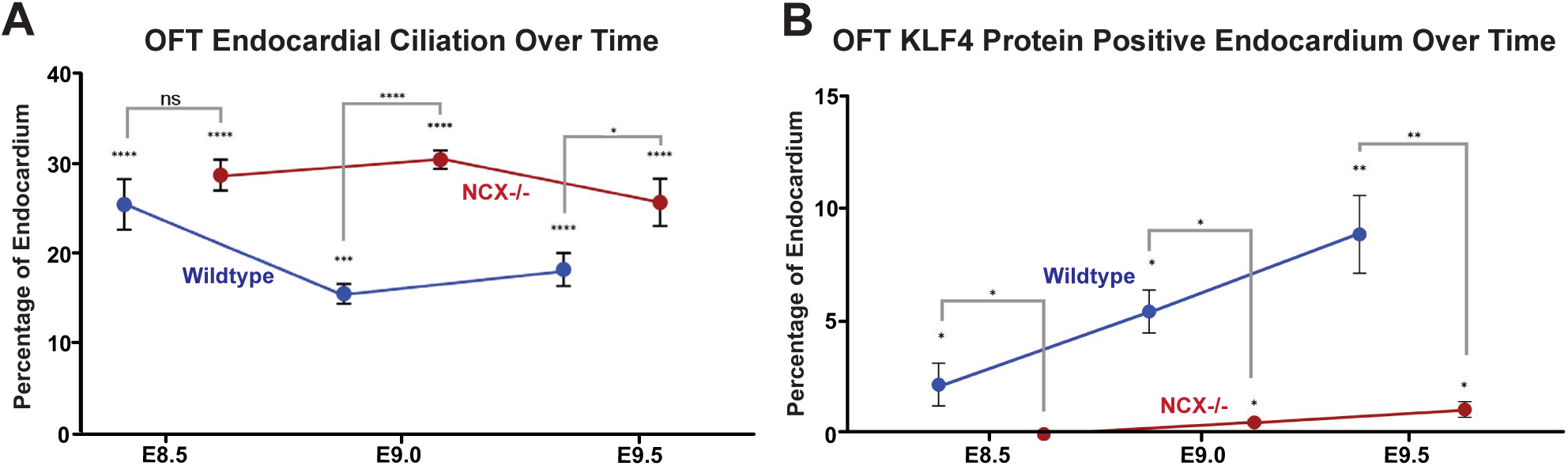
KLF4 expression at the OFT correlates with endocardial ciliation in a blood flow dependent manner. **A)** Endocardial ciliation in the OFT over time as percentage of all endocardial cells in wildtype mice (blue) and *Ncx1^−/−^* mice (red) (e8.5 (WT n=7, *Ncx1^−/−^* n=7), e9.0 (WT n=4, *Ncx1^−/−^* n=4), e9.5 (WT n=7, *Ncx1^−/−^* n=7)). **B)** KLF4 protein positive endocardial cells in the OFT over time as percentage of all endocardial cells in wildtype mice (blue) and *Ncx1^−/−^* mice (red) (e8.5 (WT n=10, *Ncx1^−/−^* n=10), e9.0 (WT n=4, *Ncx1^−/−^* n=4), e9.5 (WT n=5, *Ncx1^−/−^* n=5)). *Statistics*: ns (p > 0.05), * (p ≤ 0.05), ** (p ≤ 0.01), *** (p ≤ 0.001), **** (p ≤ 0.0001). Data are represented as mean ± SEM. Abbreviations: OFT-Outflow Tract, WT-Wildtype.

**Figure S4.**
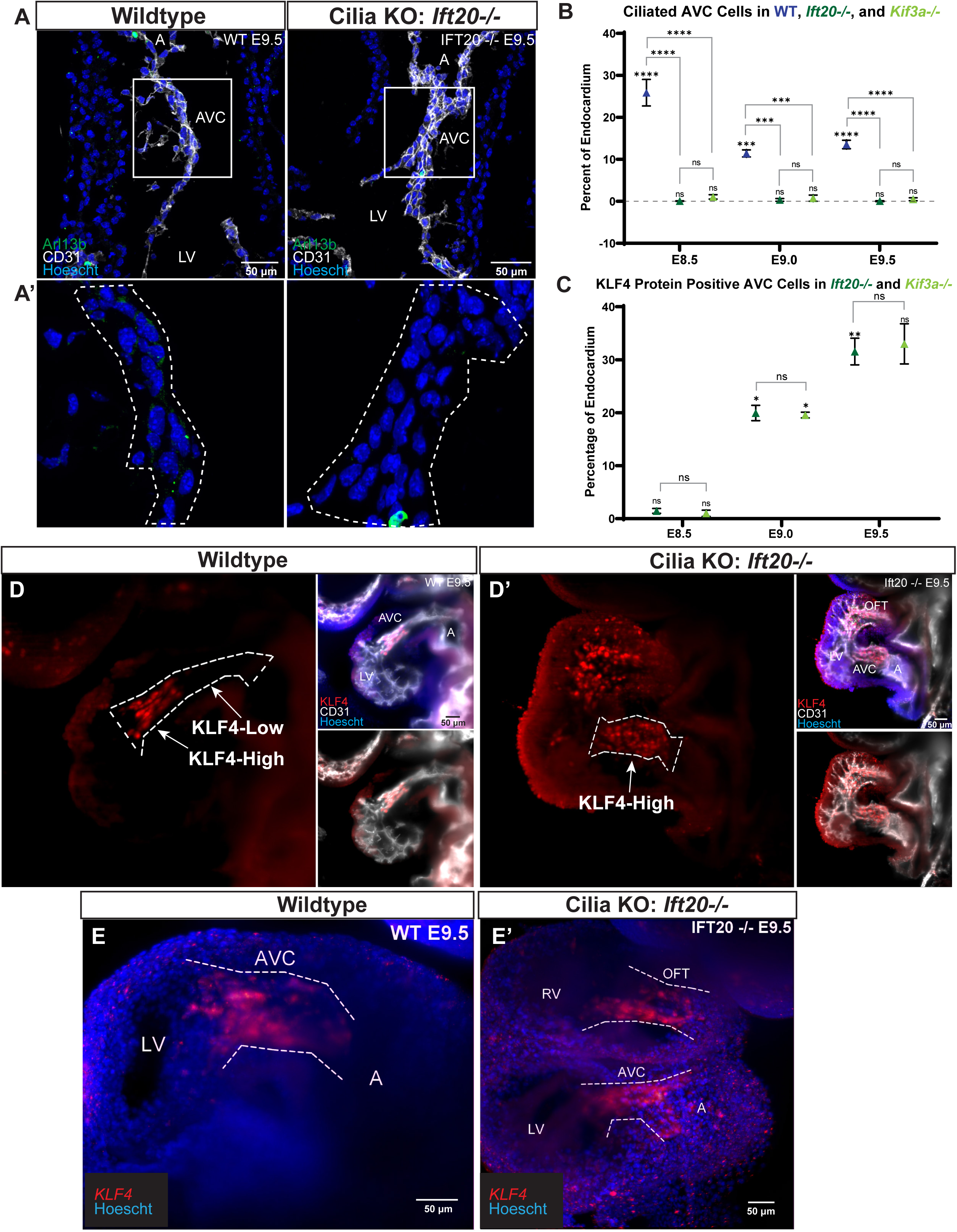
Failure of ciliogenesis results in abnormal KLF4 expression. **A)** Immunofluorescence on e9.5 wildtype and cilia KO (*Ift20^−/−^)* mouse heart sections for cilia (ARL13B, green) on endocardial cells (CD31, white). Nuclei are shown in blue (Hoescht). **A’)** Closeup of boxed AVC region without CD31 signal; the AVC is outlined in white. **B)** Ciliated endocardial cells in the AVC over time as percentage of all endocardial cells in wildtype (blue), *Ift20^−/−^* (dark green) and *Kif3a^−/−^* (light green) (e8.5 (WT n=8, *Ift20^−/−^* n=6, *Kif3a^−/−^* n=4), e9.0 (WT n=4, *Ift20^−/−^* n=3, *Kif3a^−/−^* n=3), e9.5 (WT n=6, *Ift20^−/−^* n=3, *Kif3a^−/−^* n=4)). **C)** KLF4 protein positive endocardial cells in the AVC over time as percentage of all endocardial cells in *Ift20^−/−^* (dark green) and *Kif3a^−/−^* (light green) (e8.5 (*Ift20^−/−^* n=4, *Kif3a^−/−^* n=3), e9.0 (*Ift20^−/−^* n=2, *Kif3a^−/−^* n=2), e9.5 (*Ift20^−/−^* n=4, *Kif3a^−/−^* n=2)). Immunofluorescence on e9.5 **D)** wildtype and **E)** cilia KO (*Ift20^−/−^)* whole mount hearts for KLF4 (red) in endocardial cells (CD31, white). Nuclei are shown in blue (Hoescht); the AVC is outlined in white. HCR-FISH on whole mount **F)** wildtype and **G)** Cilia KO mouse embryos at e9.5 showing *Klf2* mRNA (green)*, Klf4* mRNA (red), and nuclei (Hoescht, blue). Endocardial regions are outlined in white. *Statistics*: ns (p > 0.05), * (p ≤ 0.05), ** (p ≤ 0.01), *** (p ≤ 0.001), **** (p ≤ 0.0001). Data are represented as mean ± SEM. Abbreviations: A-Atrium, AVC-Atrioventricular Canal, LV-Left Ventricle, OFT-Outflow Tract, RV-Right Ventricle, WT-Wildtype.

**Figure S5.**
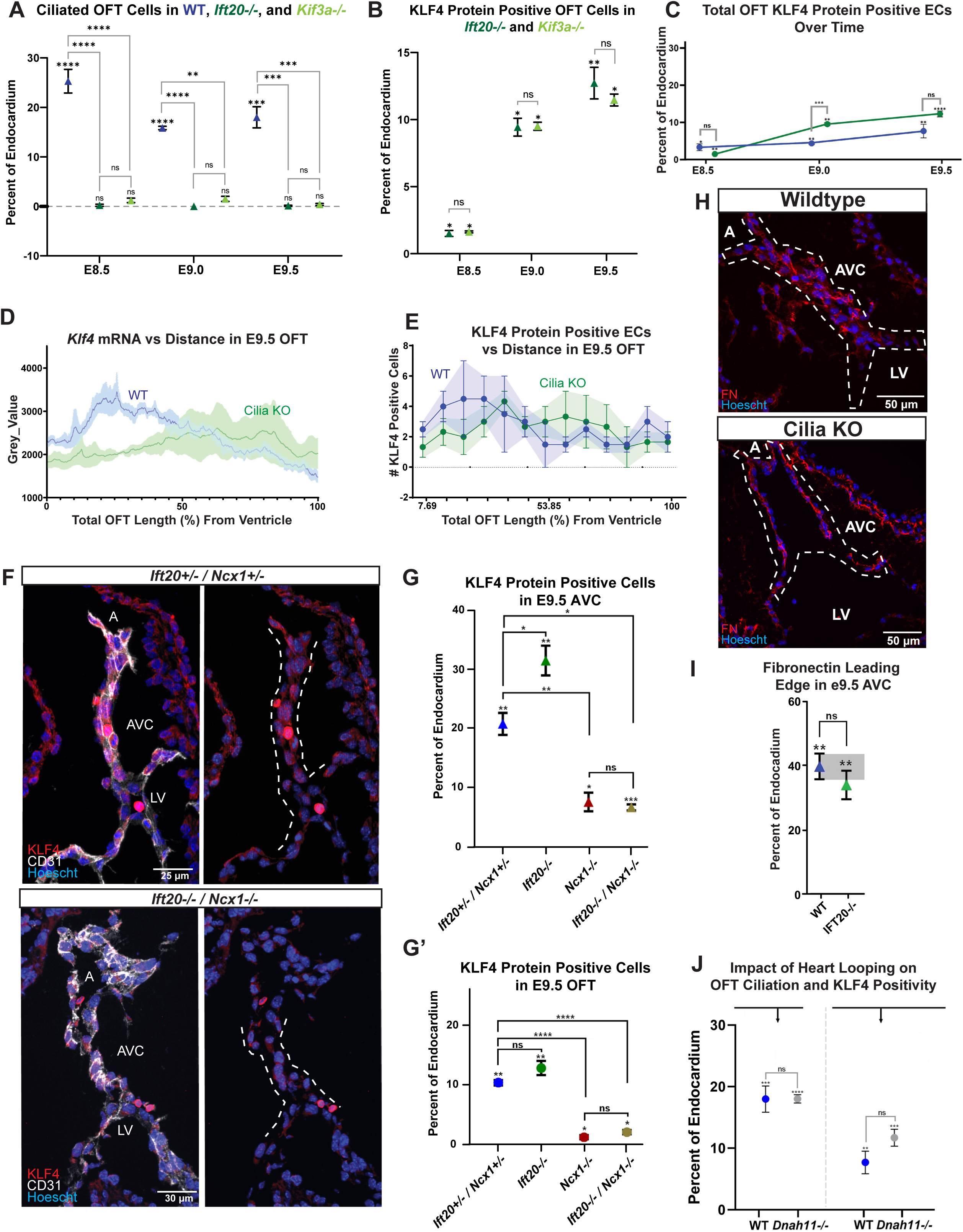
Failure of ciliogenesis results in abnormal KLF4 expression in a flow-dependent manner. **A)** Ciliated endocardial cells in the OFT over time as percentage of all endocardial cells in wildtype (blue), *Ift20^−/−^* (dark green) and *Kif3a^−/−^* (light green) (e8.5 (WT n=8, *Ift20^−/−^* n=4, *Kif3a^−/−^* n=4), e9.0 (WT n=4, *Ift20^−/−^* n=2, *Kif3a^−/−^* n=2), e9.5 (WT n=6, *Ift20^−/−^* n=4, *Kif3a^−/−^* n=3)). **B)** KLF4 protein positive endocardial cells in the OFT over time as percentage of all endocardial cells in *Ift20^−/−^* (dark green) and *Kif3a^−/−^* (light green) (e8.5 (*Ift20^−/−^* n=3, *Kif3a^−/−^* n=2), e9.0 (*Ift20^−/−^* n=2, *Kif3a^−/−^* n=2), e9.5 (*Ift20^−/−^* n=4, *Kif3a^−/−^* n=2)). **C)** KLF4 protein positive endocardial cells in the OFT over time as percentage of all endocardial cells in wildtype (blue) and cilia KO mice (green) (e8.5 (WT n=4, cilia KO n=4), e9.0 (WT n=3, cilia KO n=3), e9.5 (WT n=6, cilia KO n=6)). **D)** *Klf4* mRNA expression as measured by Grey_Value over distance in the OFT of e9.5 cilia KO mice (green, n=3); wildtype is given for comparison in blue (n=3). **E)** Number of KLF4 protein positive endocardial cells versus distance in the OFT of e9.5 cilia KO mice (green, n=3); wildtype is given for comparison in blue (n=3). **F)** Immunofluorescence on e9.5 sections of *Ift2^+/−^/ Ncx1^+/−^* and *Ift20^−/−^/ Ncx1^−/−^* hearts for KLF4 (red) in endocardial cells (CD31, white). Nuclei are shown in blue (Hoescht); the AVC is outlined in white. KLF4 protein positive endocardial cells in the e9.5 **G)** AVC and **G’)** OFT as percentage of all endocardial cells in *Ift2^+/−^/ Ncx1^+/−^* (blue, AVC n=3, OFT n=3) and *Ift20^−/−^/ Ncx1^−/−^* (brown, AVC n=4, OFT n=4). *Ift20^−/−^* (green, AVC n=4, OFT n=4) and *Ncx1^−/−^* (red, AVC n=4, OFT n=5) are given for comparison. **H)** Immunofluorescence on e9.5 wildtype and cilia KO AVC sections for Fibronectin (red). Nuclei are shown in blue (Hoescht); AVC is outlined in white. **I)** Endocardial cells with a leading edge of fibronectin as percentage of all endocardial cells in e9.5 AVCs of wildtype (blue, n=4) and cilia KO (green, n=5) mice. **J)** Ciliated endocardial cells and KLF4 protein positive endocardial cells in the OFT as percentage of all endocardial cells in e9.5 wildtype (blue) and *Dnah11^−/−^* (grey) mice (cilia: WT n=6, *Dnah11^−/−^* n=6; KLF4: WT n=6, *Dnah11^−/−^* n=6). For D) and E), distance runs left to right, Right Ventricle to Dorsal Aorta. *Statistics*: ns (p > 0.05), * (p ≤ 0.05), ** (p ≤ 0.01), *** (p ≤ 0.001), **** (p ≤ 0.0001). Data are represented as mean ± SEM. Abbreviations: A-Atrium, EC-Endocardial Cell, OFT-Outflow Tract, RV-Right Ventricle, WT-Wildtype.

**Figure S6.**
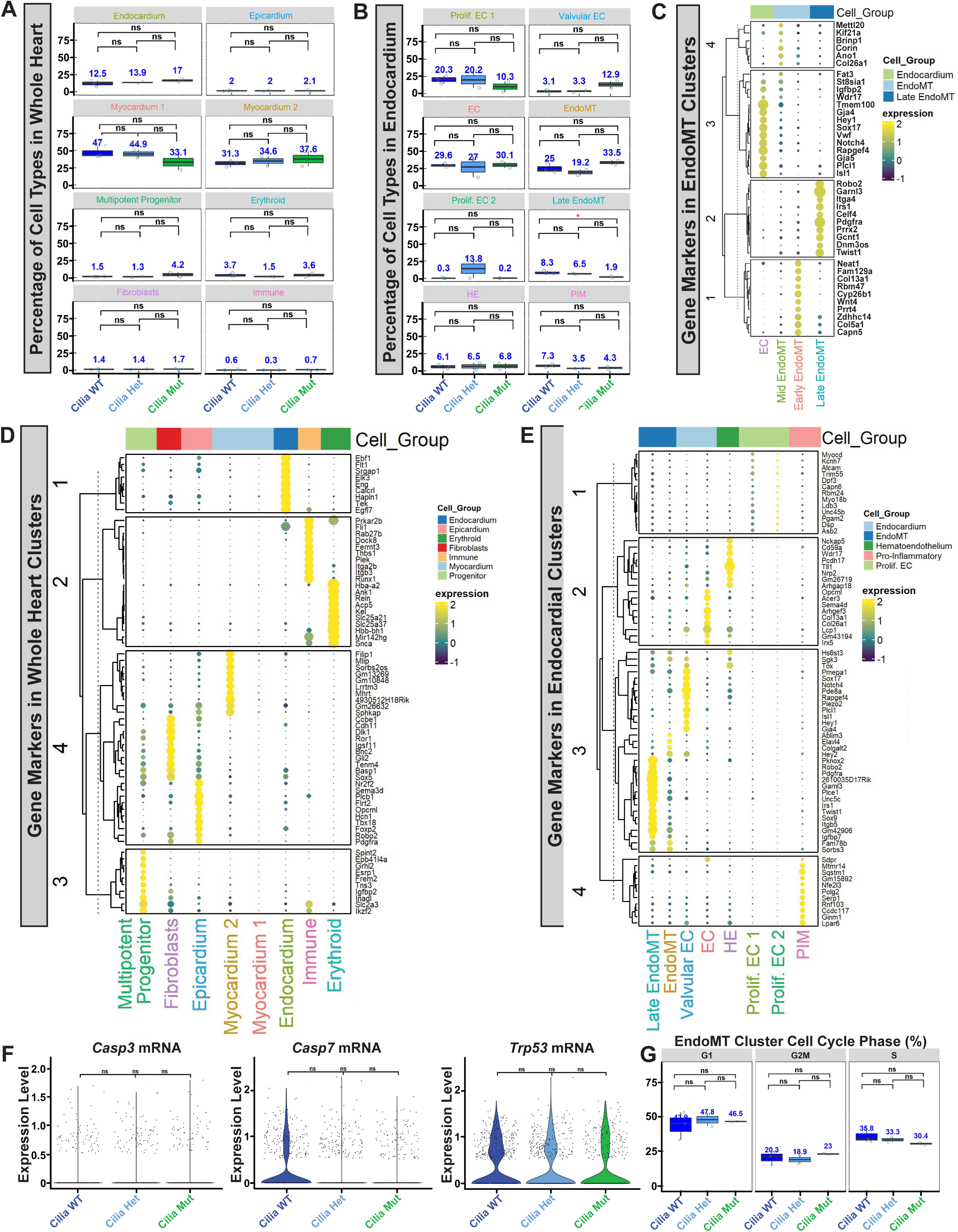
Defects in ciliogenesis block EndoMT. Graphs of the percentage of cell types in the **A)** whole heart and **B)** endocardium from snRNA-seq data for each genotype (Replicates: Cilia WT n=3, Cilia Het n=2, Cilia Mut n=2). Statistically significant numbers are given in red. Dotplots for top DEGs for each cell type in the **C)** EndoMT cluster, **D)** whole heart and **F)** endocardium from snRNA-seq data. **G)** Violin plots for proliferation gene expression in each genotype. **H)** Percentage of cells in each cell cycle phase for each genotype in the snRNA-seq EndoMT cluster (Replicates: Cilia WT n=3, Cilia Het n=2, Cilia Mut n=2). *Statistics*: ns (p > 0.05), ** (p ≤ 0.01). Data are represented as mean ± SEM. Abbreviations: A-Atrium, DEG-Differentially Expressed Gene, EC-Endocardial Cell, HE-Hematoendothelium, LV-Left Ventricle, PIM-Pro-inflammatory, WT-Wildtype.

**Figure S7.**
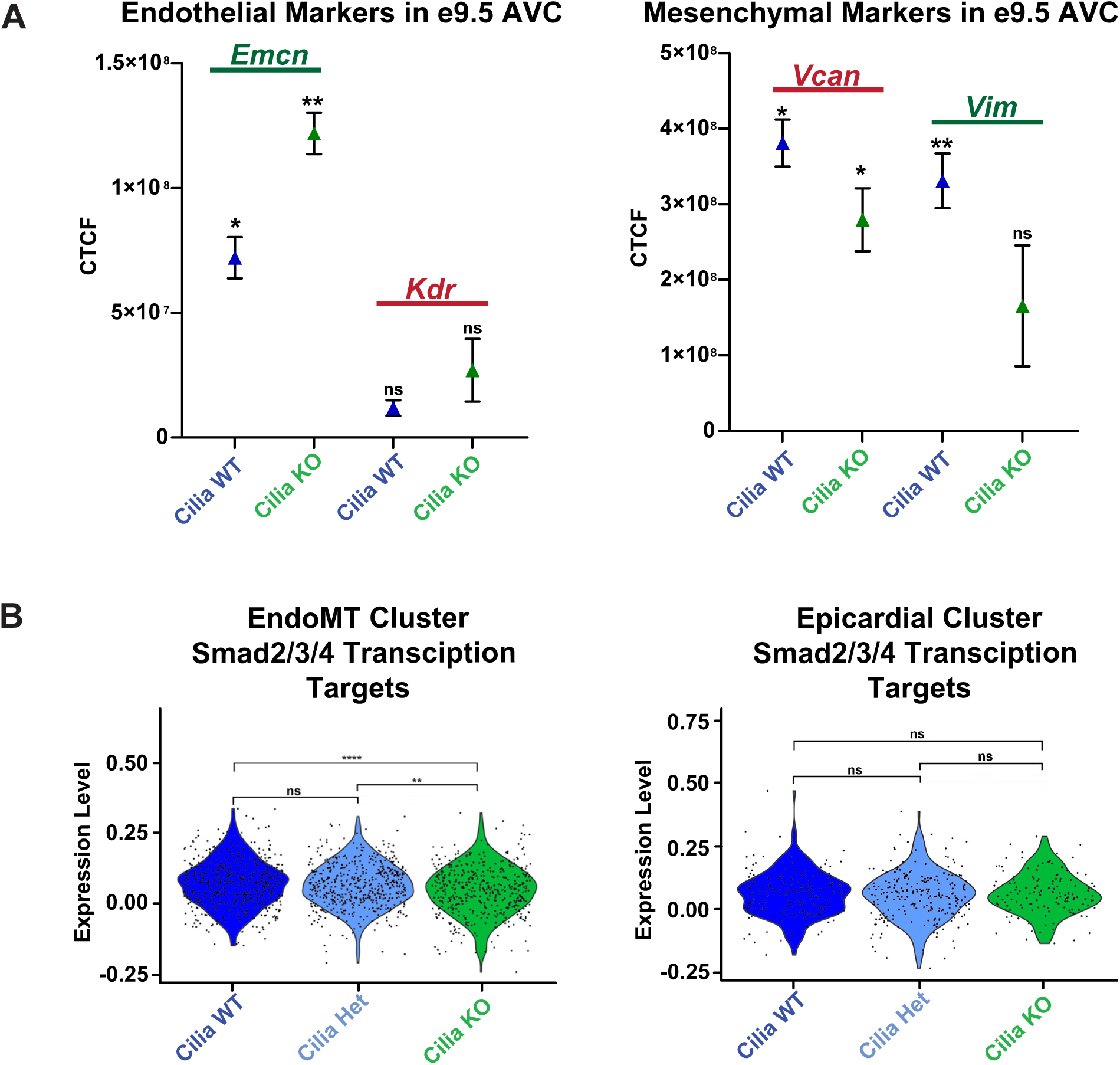
Known EndoMT molecular pathways are dysregulated in Cilia KO hearts. **A)** Corrected total cell fluorescence (CTCF) analysis of HCR-FISH for *Emcn, Kdr, Vcan, and Vim* in the e9.5 AVC (WT n=3, Cilia KO n=3). **B)** Violin plots for GeneSet Enrichment of Smad2/3/4 transcriptional targets (GSEA-MSigDB:MM14876) in each genotype for the EndoMT cluster or the Epicardial Cluster. *Statistics*: ns (p > 0.05), *** (p ≤ 0.001), **** (p ≤ 0.0001). Abbreviations: WT-Wildtype.

## References

1. Geng, X., Cha, B., Mahamud, M.R., and Srinivasan, R.S. (2017). Intraluminal valves: development, function and disease. Dis Model Mech 10, 1273–1287. 10.1242/dmm.030825.

2. Ahuja, N., Ostwald, P., Bark, D., and Garrity, D. (2020). Biomechanical Cues Direct Valvulogenesis. J Cardiovasc Dev Dis 7. 10.3390/jcdd7020018.

3. Collaborators, G.B.D.C.H.D. (2020). Global, regional, and national burden of congenital heart disease, 1990-2017: a systematic analysis for the Global Burden of Disease Study 2017. Lancet Child Adolesc Health 4, 185–200. 10.1016/S2352-4642(19)30402-X.

4. Umapathi, K.K., and Agasthi, P. (2023). Atrioventricular Canal Defects. In StatPearls.

5. Hay, D.A., and Low, F.N. (1972). The fusion of dorsal and ventral endocardial cushions in the embryonic chick heart: a study in fine structure. Am J Anat 133, 1–23. 10.1002/aja.1001330102.

6. Miao, Y., Tian, L., Martin, M., Paige, S.L., Galdos, F.X., Li, J., Klein, A., Zhang, H., Ma, N., Wei, Y., et al. (2020). Intrinsic Endocardial Defects Contribute to Hypoplastic Left Heart Syndrome. Cell Stem Cell 27, 574–589 e578. 10.1016/j.stem.2020.07.015.

7. Ho, S., Chan, W.X., and Yap, C.H. (2021). Fluid mechanics of the left atrial ligation chick embryonic model of hypoplastic left heart syndrome. Biomech Model Mechanobiol 20, 1337–1351. 10.1007/s10237-021-01447-3.

8. Andres-Delgado, L., and Mercader, N. (2016). Interplay between cardiac function and heart development. Biochim Biophys Acta 1863, 1707–1716. 10.1016/j.bbamcr.2016.03.004.

9. Courchaine, K., Rykiel, G., and Rugonyi, S. (2018). Influence of blood flow on cardiac development. Prog Biophys Mol Biol 137, 95–110. 10.1016/j.pbiomolbio.2018.05.005.

10. Midgett, M., Thornburg, K., and Rugonyi, S. (2017). Blood flow patterns underlie developmental heart defects. Am J Physiol Heart Circ Physiol 312, H632–H642. 10.1152/ajpheart.00641.2016.

11. Bartman, T., Walsh, E.C., Wen, K.K., McKane, M., Ren, J., Alexander, J., Rubenstein, P.A., and Stainier, D.Y. (2004). Early myocardial function affects endocardial cushion development in zebrafish. PLoS Biol 2, E129. 10.1371/journal.pbio.0020129.

12. Huang, C., Sheikh, F., Hollander, M., Cai, C., Becker, D., Chu, P.H., Evans, S., and Chen, J. (2003). Embryonic atrial function is essential for mouse embryogenesis, cardiac morphogenesis and angiogenesis. Development 130, 6111–6119. 10.1242/dev.00831.

13. Fukui, H., Chow, R.W., Xie, J., Foo, Y.Y., Yap, C.H., Minc, N., Mochizuki, N., and Vermot, J. (2021). Bioelectric signaling and the control of cardiac cell identity in response to mechanical forces. Science 374, 351–354. 10.1126/science.abc6229.

14. Samsa, L.A., Givens, C., Tzima, E., Stainier, D.Y., Qian, L., and Liu, J. (2015). Cardiac contraction activates endocardial Notch signaling to modulate chamber maturation in zebrafish. Development 142, 4080–4091. 10.1242/dev.125724.

15. Van der Heiden, K., Groenendijk, B.C., Hierck, B.P., Hogers, B., Koerten, H.K., Mommaas, A.M., Gittenberger-de Groot, A.C., and Poelmann, R.E. (2006). Monocilia on chicken embryonic endocardium in low shear stress areas. Dev Dyn 235, 19–28. 10.1002/dvdy.20557.

16. Slough, J., Cooney, L., and Brueckner, M. (2008). Monocilia in the embryonic mouse heart suggest a direct role for cilia in cardiac morphogenesis. Dev Dyn 237, 2304–2314. 10.1002/dvdy.21669.

17. Djenoune, L., Berg, K., Brueckner, M., and Yuan, S. (2022). A change of heart: new roles for cilia in cardiac development and disease. Nat Rev Cardiol 19, 211–227. 10.1038/s41569-021-00635-z.

18. Fulmer, D., Toomer, K., Guo, L., Moore, K., Glover, J., Moore, R., Stairley, R., Lobo, G., Zuo, X., Dang, Y., et al. (2019). Defects in the Exocyst-Cilia Machinery Cause Bicuspid Aortic Valve Disease and Aortic Stenosis. Circulation 140, 1331–1341. 10.1161/CIRCULATIONAHA.119.038376.

19. Toomer, K.A., Fulmer, D., Guo, L., Drohan, A., Peterson, N., Swanson, P., Brooks, B., Mukherjee, R., Body, S., Lipschutz, J.H., et al. (2017). A role for primary cilia in aortic valve development and disease. Dev Dyn 246, 625–634. 10.1002/dvdy.24524.

20. Toomer, K.A., Yu, M., Fulmer, D., Guo, L., Moore, K.S., Moore, R., Drayton, K.D., Glover, J., Peterson, N., Ramos-Ortiz, S., et al. (2019). Primary cilia defects causing mitral valve prolapse. Sci Transl Med 11. 10.1126/scitranslmed.aax0290.

21. Iomini, C., Tejada, K., Mo, W., Vaananen, H., and Piperno, G. (2004). Primary cilia of human endothelial cells disassemble under laminar shear stress. J Cell Biol 164, 811–817. 10.1083/jcb.200312133.

22. Egorova, A.D., Khedoe, P.P., Goumans, M.J., Yoder, B.K., Nauli, S.M., ten Dijke, P., Poelmann, R.E., and Hierck, B.P. (2011). Lack of primary cilia primes shear-induced endothelial-to-mesenchymal transition. Circ Res 108, 1093–1101. 10.1161/CIRCRESAHA.110.231860.

23. Li, X., Lu, Q., Peng, Y., Geng, F., Shao, X., Zhou, H., Cao, Y., and Zhang, R. (2020). Primary cilia mediate Klf2-dependant Notch activation in regenerating heart. Protein Cell 11, 433–445. 10.1007/s13238-020-00695-w.

24. Boselli, F., and Vermot, J. (2016). Live imaging and modeling for shear stress quantification in the embryonic zebrafish heart. Methods 94, 129–134. 10.1016/j.ymeth.2015.09.017.

25. Chiplunkar, A.R., Curtis, B.C., Eades, G.L., Kane, M.S., Fox, S.J., Haar, J.L., and Lloyd, J.A. (2013). The Kruppel-like factor 2 and Kruppel-like factor 4 genes interact to maintain endothelial integrity in mouse embryonic vasculogenesis. BMC Dev Biol 13, 40. 10.1186/1471-213X-13-40.

26. Zhang, Y., Li, X., Gao, S., Liao, Y., Luo, Y., Liu, M., Bian, Y., Xiong, H., Yue, Y., and He, A. (2023). Genetic reporter for live tracing fluid flow forces during cell fate segregation in mouse blastocyst development. Cell Stem Cell 30, 1110–1123 e1119. 10.1016/j.stem.2023.07.003.

27. Novodvorsky, P., and Chico, T.J. (2014). The role of the transcription factor KLF2 in vascular development and disease. Prog Mol Biol Transl Sci 124, 155–188. 10.1016/B978-0-12-386930-2.00007-0.

28. de Soysa, T.Y., Ranade, S.S., Okawa, S., Ravichandran, S., Huang, Y., Salunga, H.T., Schricker, A., Del Sol, A., Gifford, C.A., and Srivastava, D. (2019). Single-cell analysis of cardiogenesis reveals basis for organ-level developmental defects. Nature 572, 120–124. 10.1038/s41586-019-1414-x.

29. Bourillot, P.Y., and Savatier, P. (2010). Kruppel-like transcription factors and control of pluripotency. BMC Biol 8, 125. 10.1186/1741-7007-8-125.

30. Todorovic, V., Finnegan, E., Freyer, L., Zilberberg, L., Ota, M., and Rifkin, D.B. (2011). Long form of latent TGF-beta binding protein 1 (Ltbp1L) regulates cardiac valve development. Dev Dyn 240, 176–187. 10.1002/dvdy.22521.

31. Heckel, E., Boselli, F., Roth, S., Krudewig, A., Belting, H.G., Charvin, G., and Vermot, J. (2015). Oscillatory Flow Modulates Mechanosensitive klf2a Expression through trpv4 and trpp2 during Heart Valve Development. Curr Biol 25, 1354–1361. 10.1016/j.cub.2015.03.038.

32. Ghaleb, A.M., and Yang, V.W. (2017). Kruppel-like factor 4 (KLF4): What we currently know. Gene 611, 27–37. 10.1016/j.gene.2017.02.025.

33. Zheng, Q., Zou, Y., Teng, P., Chen, Z., Wu, Y., Dai, X., Li, X., Hu, Z., Wu, S., Xu, Y., et al. (2022). Mechanosensitive Channel PIEZO1 Senses Shear Force to Induce KLF2/4 Expression via CaMKII/MEKK3/ERK5 Axis in Endothelial Cells. Cells 11. 10.3390/cells11142191.

34. Peterson, L.M., Jenkins, M.W., Gu, S., Barwick, L., Watanabe, M., and Rollins, A.M. (2012). 4D shear stress maps of the developing heart using Doppler optical coherence tomography. Biomed Opt Express 3, 3022–3032. 10.1364/BOE.3.003022.

35. Lee, J., Vedula, V., Baek, K.I., Chen, J., Hsu, J.J., Ding, Y., Chang, C.C., Kang, H., Small, A., Fei, P., et al. (2018). Spatial and temporal variations in hemodynamic forces initiate cardiac trabeculation. JCI Insight 3. 10.1172/jci.insight.96672.

36. Alejandre Alcazar, M.A., Kaschwich, M., Ertsey, R., Preuss, S., Milla, C., Mujahid, S., Masumi, J., Khan, S., Mokres, L.M., Tian, L., et al. (2018). Elafin Treatment Rescues EGFR-Klf4 Signaling and Lung Cell Survival in Ventilated Newborn Mice. Am J Respir Cell Mol Biol 59, 623–634. 10.1165/rcmb.2017-0332OC.

37. Andueza, A., Kumar, S., Kim, J., Kang, D.W., Mumme, H.L., Perez, J.I., Villa-Roel, N., and Jo, H. (2020). Endothelial Reprogramming by Disturbed Flow Revealed by Single-Cell RNA and Chromatin Accessibility Study. Cell Rep 33, 108491. 10.1016/j.celrep.2020.108491.

38. Jiang, Y.Z., Jimenez, J.M., Ou, K., McCormick, M.E., Zhang, L.D., and Davies, P.F. (2014). Hemodynamic disturbed flow induces differential DNA methylation of endothelial Kruppel-Like Factor 4 promoter in vitro and in vivo. Circ Res 115, 32–43. 10.1161/CIRCRESAHA.115.303883.

39. Koushik, S.V., Wang, J., Rogers, R., Moskophidis, D., Lambert, N.A., Creazzo, T.L., and Conway, S.J. (2001). Targeted inactivation of the sodium-calcium exchanger (Ncx1) results in the lack of a heartbeat and abnormal myofibrillar organization. FASEB J 15, 1209–1211. 10.1096/fj.00-0696fje.

40. Cuttano, R., Rudini, N., Bravi, L., Corada, M., Giampietro, C., Papa, E., Morini, M.F., Maddaluno, L., Baeyens, N., Adams, R.H., et al. (2016). KLF4 is a key determinant in the development and progression of cerebral cavernous malformations. EMBO Mol Med 8, 6–24. 10.15252/emmm.201505433.

41. Mastej, V., Axen, C., Wary, A., Minshall, R.D., and Wary, K.K. (2022). A requirement for Kruppel Like Factor-4 in the maintenance of endothelial cell quiescence. Front Cell Dev Biol 10, 1003028. 10.3389/fcell.2022.1003028.

42. Wang, X., Li, X., Huang, C., Li, L., Qu, H., Yu, X., Ni, H., and Cui, Q. (2016). Kruppel-like factor 4 (KLF-4) inhibits the epithelial-to-mesenchymal transition and proliferation of human endometrial carcinoma cells. Gynecol Endocrinol 32, 772–776. 10.3109/09513590.2016.1163673.

43. Zhang, Y., Li, C., Huang, Y., Zhao, S., Xu, Y., Chen, Y., Jiang, F., Tao, L., and Shen, X. (2020). EOFAZ inhibits endothelial-to-mesenchymal transition through downregulation of KLF4. Int J Mol Med 46, 300–310. 10.3892/ijmm.2020.4572.

44. Lotto, J., Cullum, R., Drissler, S., Arostegui, M., Garside, V.C., Fuglerud, B.M., Clement-Ranney, M., Thakur, A., Underhill, T.M., and Hoodless, P.A. (2023). Cell diversity and plasticity during atrioventricular heart valve EMTs. Nat Commun 14, 5567. 10.1038/s41467-023-41279-6.

45. Mjaatvedt, C.H., Lepera, R.C., and Markwald, R.R. (1987). Myocardial specificity for initiating endothelial-mesenchymal cell transition in embryonic chick heart correlates with a particulate distribution of fibronectin. Dev Biol 119, 59–67. 10.1016/0012-1606(87)90206-5.

46. Liu, S., Yang, H., Chen, Y., He, B., and Chen, Q. (2016). Kruppel-Like Factor 4 Enhances Sensitivity of Cisplatin to Lung Cancer Cells and Inhibits Regulating Epithelial- to-Mesenchymal Transition. Oncol Res 24, 81–87. 10.3727/096504016X14597766487717.

47. Subbalakshmi, A.R., Sahoo, S., McMullen, I., Saxena, A.N., Venugopal, S.K., Somarelli, J.A., and Jolly, M.K. (2021). KLF4 Induces Mesenchymal-Epithelial Transition (MET) by Suppressing Multiple EMT-Inducing Transcription Factors. Cancers (Basel) 13. 10.3390/cancers13205135.

48. Follit, J.A., Tuft, R.A., Fogarty, K.E., and Pazour, G.J. (2006). The intraflagellar transport protein IFT20 is associated with the Golgi complex and is required for cilia assembly. Mol Biol Cell 17, 3781–3792. 10.1091/mbc.e06-02-0133.

49. Marszalek, J.R., Ruiz-Lozano, P., Roberts, E., Chien, K.R., and Goldstein, L.S. (1999). Situs inversus and embryonic ciliary morphogenesis defects in mouse mutants lacking the KIF3A subunit of kinesin-II. Proc Natl Acad Sci U S A 96, 5043–5048. 10.1073/pnas.96.9.5043.

50. Ten Dijke, P., Egorova, A.D., Goumans, M.J., Poelmann, R.E., and Hierck, B.P. (2012). TGF-beta signaling in endothelial-to-mesenchymal transition: the role of shear stress and primary cilia. Sci Signal 5, pt2. 10.1126/scisignal.2002722.

51. McGrath, J., Somlo, S., Makova, S., Tian, X., and Brueckner, M. (2003). Two populations of node monocilia initiate left-right asymmetry in the mouse. Cell 114, 61–73. 10.1016/s0092-8674(03)00511-7.

52. Wei, D., Kanai, M., Huang, S., and Xie, K. (2006). Emerging role of KLF4 in human gastrointestinal cancer. Carcinogenesis 27, 23–31. 10.1093/carcin/bgi243.

53. Yang, H., Xi, X., Zhao, B., Su, Z., and Wang, Z. (2018). KLF4 protects brain microvascular endothelial cells from ischemic stroke induced apoptosis by transcriptionally activating MALAT1. Biochem Biophys Res Commun 495, 2376–2382. 10.1016/j.bbrc.2017.11.205.

54. Cao, C., Zhou, Y., Zhang, Y., Ma, Y., Du, S., Fan, L., Niu, R., Zhang, Y., and He, M. (2022). GCN5 participates in KLF4-VEGFA feedback to promote endometrial angiogenesis. iScience 25, 104509. 10.1016/j.isci.2022.104509.

55. Choi, D., Park, E., Jung, E., Seong, Y.J., Hong, M., Lee, S., Burford, J., Gyarmati, G., Peti-Peterdi, J., Srikanth, S., et al. (2017). ORAI1 Activates Proliferation of Lymphatic Endothelial Cells in Response to Laminar Flow Through Kruppel-Like Factors 2 and 4. Circ Res 120, 1426–1439. 10.1161/CIRCRESAHA.116.309548.

56. Wang, Y., Yang, C., Gu, Q., Sims, M., Gu, W., Pfeffer, L.M., and Yue, J. (2015). KLF4 Promotes Angiogenesis by Activating VEGF Signaling in Human Retinal Microvascular Endothelial Cells. PLoS One 10, e0130341. 10.1371/journal.pone.0130341.

57. Li, H., Li, C., Zheng, T., Wang, Y., Wang, J., Fan, X., Zheng, X., Tian, G., Yuan, Z., and Chen, T. (2023). Cardiac Fibroblast Activation Induced by Oxygen-Glucose Deprivation Depends on the HIF-1alpha/miR-212-5p/KLF4 Pathway. J Cardiovasc Transl Res 16, 778–792. 10.1007/s12265-023-10360-2.

58. Aksoy, I., Giudice, V., Delahaye, E., Wianny, F., Aubry, M., Mure, M., Chen, J., Jauch, R., Bogu, G.K., Nolden, T., et al. (2014). Klf4 and Klf5 differentially inhibit mesoderm and endoderm differentiation in embryonic stem cells. Nat Commun 5, 3719. 10.1038/ncomms4719.

59. Malaab, M., Renaud, L., Takamura, N., Zimmerman, K.D., da Silveira, W.A., Ramos, P.S., Haddad, S., Peters-Golden, M., Penke, L.R., Wolf, B., et al. (2022). Antifibrotic factor KLF4 is repressed by the miR-10/TFAP2A/TBX5 axis in dermal fibroblasts: insights from twins discordant for systemic sclerosis. Ann Rheum Dis 81, 268–277. 10.1136/annrheumdis-2021-221050.

60. Han, H., Cho, J.W., Lee, S., Yun, A., Kim, H., Bae, D., Yang, S., Kim, C.Y., Lee, M., Kim, E., et al. (2018). TRRUST v2: an expanded reference database of human and mouse transcriptional regulatory interactions. Nucleic Acids Res 46, D380–D386. 10.1093/nar/gkx1013.

61. Bailey, T.L., Johnson, J., Grant, C.E., and Noble, W.S. (2015). The MEME Suite. Nucleic Acids Res 43, W39–49. 10.1093/nar/gkv416.

62. Granados-Riveron, J.T., and Brook, J.D. (2012). The impact of mechanical forces in heart morphogenesis. Circ Cardiovasc Genet 5, 132–142. 10.1161/CIRCGENETICS.111.961086.

63. Midgett, M., Lopez, C.S., David, L., Maloyan, A., and Rugonyi, S. (2017). Increased Hemodynamic Load in Early Embryonic Stages Alters Endocardial to Mesenchymal Transition. Front Physiol 8, 56. 10.3389/fphys.2017.00056.

64. Steed, E., Boselli, F., and Vermot, J. (2016). Hemodynamics driven cardiac valve morphogenesis. Biochim Biophys Acta 1863, 1760–1766. 10.1016/j.bbamcr.2015.11.014.

65. Huang, R.T., Wu, D., Meliton, A., Oh, M.J., Krause, M., Lloyd, J.A., Nigdelioglu, R., Hamanaka, R.B., Jain, M.K., Birukova, A., et al. (2017). Experimental Lung Injury Reduces Kruppel-like Factor 2 to Increase Endothelial Permeability via Regulation of RAPGEF3-Rac1 Signaling. Am J Respir Crit Care Med 195, 639–651. 10.1164/rccm.201604-0668OC.

66. Steed, E., Faggianelli, N., Roth, S., Ramspacher, C., Concordet, J.P., and Vermot, J. (2016). klf2a couples mechanotransduction and zebrafish valve morphogenesis through fibronectin synthesis. Nat Commun 7, 11646. 10.1038/ncomms11646.

67. Nakajima, H., and Mochizuki, N. (2017). Flow pattern-dependent endothelial cell responses through transcriptional regulation. Cell Cycle 16, 1893–1901. 10.1080/15384101.2017.1364324.

68. Jones, E.A., Baron, M.H., Fraser, S.E., and Dickinson, M.E. (2004). Measuring hemodynamic changes during mammalian development. Am J Physiol Heart Circ Physiol 287, H1561–1569. 10.1152/ajpheart.00081.2004.

69. Mack, J.J., Mosqueiro, T.S., Archer, B.J., Jones, W.M., Sunshine, H., Faas, G.C., Briot, A., Aragon, R.L., Su, T., Romay, M.C., et al. (2017). NOTCH1 is a mechanosensor in adult arteries. Nat Commun 8, 1620. 10.1038/s41467-017-01741-8.

70. Kefaloyianni, E., and Coetzee, W.A. (2011). Transcriptional remodeling of ion channel subunits by flow adaptation in human coronary artery endothelial cells. J Vasc Res 48, 357–367. 10.1159/000323475.

71. Lai, A., Chen, Y.C., Cox, C.D., Jaworowski, A., Peter, K., and Baratchi, S. (2021). Analyzing the shear-induced sensitization of mechanosensitive ion channel Piezo-1 in human aortic endothelial cells. J Cell Physiol 236, 2976–2987. 10.1002/jcp.30056.

72. Yarishkin, O., Phuong, T.T.T., Baumann, J.M., De Ieso, M.L., Vazquez-Chona, F., Rudzitis, C.N., Sundberg, C., Lakk, M., Stamer, W.D., and Krizaj, D. (2021). Piezo1 channels mediate trabecular meshwork mechanotransduction and promote aqueous fluid outflow. J Physiol 599, 571–592. 10.1113/JP281011.

73. Gopalakrishnan, J., Feistel, K., Friedrich, B.M., Grapin-Botton, A., Jurisch-Yaksi, N., Mass, E., Mick, D.U., Muller, R.U., May-Simera, H., Schermer, B., et al. (2023). Emerging principles of primary cilia dynamics in controlling tissue organization and function. EMBO J, e113891. 10.15252/embj.2023113891.

74. Djenoune, L., Mahamdeh, M., Truong, T.V., Nguyen, C.T., Fraser, S.E., Brueckner, M., Howard, J., and Yuan, S. (2023). Cilia function as calcium-mediated mechanosensors that instruct left-right asymmetry. Science 379, 71–78. 10.1126/science.abq7317.

75. Fang, Y., Wu, D., and Birukov, K.G. (2019). Mechanosensing and Mechanoregulation of Endothelial Cell Functions. Compr Physiol 9, 873–904. 10.1002/cphy.c180020.

76. Hierck, B.P., Van der Heiden, K., Alkemade, F.E., Van de Pas, S., Van Thienen, J.V., Groenendijk, B.C., Bax, W.H., Van der Laarse, A., Deruiter, M.C., Horrevoets, A.J., and Poelmann, R.E. (2008). Primary cilia sensitize endothelial cells for fluid shear stress. Dev Dyn 237, 725–735. 10.1002/dvdy.21472.

77. Kalogirou, S., Malissovas, N., Moro, E., Argenton, F., Stainier, D.Y., and Beis, D. (2014). Intracardiac flow dynamics regulate atrioventricular valve morphogenesis. Cardiovasc Res 104, 49–60. 10.1093/cvr/cvu186.

78. Burnicka-Turek, O., Steimle, J.D., Huang, W., Felker, L., Kamp, A., Kweon, J., Peterson, M., Reeves, R.H., Maslen, C.L., Gruber, P.J., et al. (2016). Cilia gene mutations cause atrioventricular septal defects by multiple mechanisms. Hum Mol Genet 25, 3011–3028. 10.1093/hmg/ddw155.

79. Ma, B., Zhang, L., Zou, Y., He, R., Wu, Q., Han, C., and Zhang, B. (2019). Reciprocal regulation of integrin beta4 and KLF4 promotes gliomagenesis through maintaining cancer stem cell traits. J Exp Clin Cancer Res 38, 23. 10.1186/s13046-019-1034-1.

80. Goetz, J.G., Steed, E., Ferreira, R.R., Roth, S., Ramspacher, C., Boselli, F., Charvin, G., Liebling, M., Wyart, C., Schwab, Y., and Vermot, J. (2014). Endothelial cilia mediate low flow sensing during zebrafish vascular development. Cell Rep 6, 799–808. 10.1016/j.celrep.2014.01.032.

81. Rahimzadeh, J., Meng, F., Sachs, F., Wang, J., Verma, D., and Hua, S.Z. (2011). Real-time observation of flow-induced cytoskeletal stress in living cells. Am J Physiol Cell Physiol 301, C646–652. 10.1152/ajpcell.00099.2011.

82. Park, C.S., Shen, Y., Lewis, A., and Lacorazza, H.D. (2016). Role of the reprogramming factor KLF4 in blood formation. J Leukoc Biol 99, 673–685. 10.1189/jlb.1RU1215-539R.

83. Penke, L.R., Speth, J.M., Huang, S.K., Fortier, S.M., Baas, J., and Peters-Golden, M. (2022). KLF4 is a therapeutically tractable brake on fibroblast activation that promotes resolution of pulmonary fibrosis. JCI Insight 7. 10.1172/jci.insight.160688.

84. Tiwari, A., Swamynathan, S., Alexander, N., Gnalian, J., Tian, S., Kinchington, P.R., and Swamynathan, S.K. (2019). KLF4 Regulates Corneal Epithelial Cell Cycle Progression by Suppressing Canonical TGF-beta Signaling and Upregulating CDK Inhibitors P16 and P27. Invest Ophthalmol Vis Sci 60, 731–740. 10.1167/iovs.18-26423.

85. Labour, M.N., Riffault, M., Christensen, S.T., and Hoey, D.A. (2016). TGFbeta1 - induced recruitment of human bone mesenchymal stem cells is mediated by the primary cilium in a SMAD3-dependent manner. Sci Rep 6, 35542. 10.1038/srep35542.

86. Pala, R., Alomari, N., and Nauli, S.M. (2017). Primary Cilium-Dependent Signaling Mechanisms. Int J Mol Sci 18. 10.3390/ijms18112272.

87. Yanardag, S., and Pugacheva, E.N. (2021). Primary Cilium Is Involved in Stem Cell Differentiation and Renewal through the Regulation of Multiple Signaling Pathways. Cells 10. 10.3390/cells10061428.

88. Mirvis, M., Siemers, K.A., Nelson, W.J., and Stearns, T.P. (2019). Primary cilium loss in mammalian cells occurs predominantly by whole-cilium shedding. PLoS Biol 17, e3000381. 10.1371/journal.pbio.3000381.

89. Ishikawa, H., Thompson, J., Yates, J.R., 3rd, and Marshall, W.F. (2012). Proteomic analysis of mammalian primary cilia. Curr Biol 22, 414–419. 10.1016/j.cub.2012.01.031.

90. Overgaard, C.E., Sanzone, K.M., Spiczka, K.S., Sheff, D.R., Sandra, A., and Yeaman, C. (2009). Deciliation is associated with dramatic remodeling of epithelial cell junctions and surface domains. Mol Biol Cell 20, 102–113. 10.1091/mbc.e08-07-0741.

91. Plotnikova, O.V., Nikonova, A.S., Loskutov, Y.V., Kozyulina, P.Y., Pugacheva, E.N., and Golemis, E.A. (2012). Calmodulin activation of Aurora-A kinase (AURKA) is required during ciliary disassembly and in mitosis. Mol Biol Cell 23, 2658–2670. 10.1091/mbc.E11-12-1056.

92. Rozycki, M., Lodyga, M., Lam, J., Miranda, M.Z., Fatyol, K., Speight, P., and Kapus, A. (2014). The fate of the primary cilium during myofibroblast transition. Mol Biol Cell 25, 643–657. 10.1091/mbc.E13-07-0429.

93. Vion, A.C., Alt, S., Klaus-Bergmann, A., Szymborska, A., Zheng, T., Perovic, T., Hammoutene, A., Oliveira, M.B., Bartels-Klein, E., Hollfinger, I., et al. (2018). Primary cilia sensitize endothelial cells to BMP and prevent excessive vascular regression. J Cell Biol 217, 1651–1665. 10.1083/jcb.201706151.

94. Li, Y.Z., Wen, L., Wei, X., Wang, Q.R., Xu, L.W., Zhang, H.M., and Liu, W.C. (2016). Inhibition of miR-7 promotes angiogenesis in human umbilical vein endothelial cells by upregulating VEGF via KLF4. Oncol Rep 36, 1569–1575. 10.3892/or.2016.4912.

95. Pestel, J., Ramadass, R., Gauvrit, S., Helker, C., Herzog, W., and Stainier, D.Y. (2016). Real-time 3D visualization of cellular rearrangements during cardiac valve formation. Development 143, 2217–2227. 10.1242/dev.133272.

96. Fujimoto, S., Hayashi, R., Hara, S., Sasamoto, Y., Harrington, J., Tsujikawa, M., and Nishida, K. (2019). KLF4 prevents epithelial to mesenchymal transition in human corneal epithelial cells via endogenous TGF-beta2 suppression. Regen Ther 11, 249–257. 10.1016/j.reth.2019.08.003.

97. Liu, Y.N., Abou-Kheir, W., Yin, J.J., Fang, L., Hynes, P., Casey, O., Hu, D., Wan, Y., Seng, V., Sheppard-Tillman, H., et al. (2012). Critical and reciprocal regulation of KLF4 and SLUG in transforming growth factor beta-initiated prostate cancer epithelial-mesenchymal transition. Mol Cell Biol 32, 941–953. 10.1128/MCB.06306-11.

98. Jonassen, J.A., San Agustin, J., Follit, J.A., and Pazour, G.J. (2008). Deletion of IFT20 in the mouse kidney causes misorientation of the mitotic spindle and cystic kidney disease. J Cell Biol 183, 377–384. 10.1083/jcb.200808137.

99. Kaufman, M.H., and Kaufman, M.H. (1992). The atlas of mouse development (Academic Press).

100. Hogan, B.M., Bos, F.L., Bussmann, J., Witte, M., Chi, N.C., Duckers, H.J., and Schulte-Merker, S. (2009). Ccbe1 is required for embryonic lymphangiogenesis and venous sprouting. Nat Genet 41, 396–398. 10.1038/ng.321.

101. Yuan, S., Zhao, L., Brueckner, M., and Sun, Z. (2015). Intraciliary calcium oscillations initiate vertebrate left-right asymmetry. Curr Biol 25, 556–567. 10.1016/j.cub.2014.12.051.

102. Westerfield, M. (2000). The zebrafish book : a guide for the laboratory use of zebrafish (Danio rerio), Ed. 5. Edition (M. Westerfield).

103. Barish, S., Berg, K., Drozd, J., Berglund-Brown, I., Khizir, L., Wasson, L.K., Seidman, C.E., Seidman, J.G., Chen, S., and Brueckner, M. (2023). The H2Bub1-deposition complex is required for human and mouse cardiogenesis. Development 150. 10.1242/dev.201899.

104. Lowe, L.A., Supp, D.M., Sampath, K., Yokoyama, T., Wright, C.V., Potter, S.S., Overbeek, P., and Kuehn, M.R. (1996). Conserved left-right asymmetry of nodal expression and alterations in murine situs inversus. Nature 381, 158–161. 10.1038/381158a0.

105. Choi, H.M.T., Schwarzkopf, M., Fornace, M.E., Acharya, A., Artavanis, G., Stegmaier, J., Cunha, A., and Pierce, N.A. (2018). Third-generation in situ hybridization chain reaction: multiplexed, quantitative, sensitive, versatile, robust. Development 145. 10.1242/dev.165753.

106. Schlaeppi, A., Graves, A., Weber, M., and Huisken, J. (2021). Light Sheet Microscopy of Fast Cardiac Dynamics in Zebrafish Embryos. J Vis Exp. 10.3791/62741.

107. Stringer, C., Wang, T., Michaelos, M., and Pachitariu, M. (2021). Cellpose: a generalist algorithm for cellular segmentation. Nat Methods 18, 100–106. 10.1038/s41592-020-01018-x.

108. Schindelin, J., Arganda-Carreras, I., Frise, E., Kaynig, V., Longair, M., Pietzsch, T., Preibisch, S., Rueden, C., Saalfeld, S., Schmid, B., et al. (2012). Fiji: an open-source platform for biological-image analysis. Nat Methods 9, 676–682. 10.1038/nmeth.2019.

109. Wan, Y., Otsuna, H., Holman, H.A., Bagley, B., Ito, M., Lewis, A.K., Colasanto, M., Kardon, G., Ito, K., and Hansen, C. (2017). FluoRender: joint freehand segmentation and visualization for many-channel fluorescence data analysis. BMC Bioinformatics 18, 280. 10.1186/s12859-017-1694-9.

110. Nadelmann, E.R., Gorham, J.M., Reichart, D., Delaughter, D.M., Wakimoto, H., Lindberg, E.L., Litvinukova, M., Maatz, H., Curran, J.J., Ischiu Gutierrez, D., et al. (2021). Isolation of Nuclei from Mammalian Cells and Tissues for Single-Nucleus Molecular Profiling. Curr Protoc 1, e132. 10.1002/cpz1.132.

111. Reichart, D., Newby, G.A., Wakimoto, H., Lun, M., Gorham, J.M., Curran, J.J., Raguram, A., DeLaughter, D.M., Conner, D.A., Marsiglia, J.D.C., et al. (2023). Efficient in vivo genome editing prevents hypertrophic cardiomyopathy in mice. Nat Med 29, 412–421. 10.1038/s41591-022-02190-7.

112. Wolock, S.L., Lopez, R., and Klein, A.M. (2019). Scrublet: Computational Identification of Cell Doublets in Single-Cell Transcriptomic Data. Cell Syst 8, 281–291 e289. 10.1016/j.cels.2018.11.005.

113. Van Rossum, G., and Drake, F.L. (2011). The Python language reference manual : for Python version 3.2, Rev. and updated. Edition (Network Theory Ltd).

114. Team, R.D.C., and Team, R.D.C. (2011). The R reference manual (Network Theory Ltd).

115. Butler, A., Hoffman, P., Smibert, P., Papalexi, E., and Satija, R. (2018). Integrating single-cell transcriptomic data across different conditions, technologies, and species. Nat Biotechnol 36, 411–420. 10.1038/nbt.4096.

116. Hao, Y., Hao, S., Andersen-Nissen, E., Mauck, W.M., 3rd, Zheng, S., Butler, A., Lee, M.J., Wilk, A.J., Darby, C., Zager, M., et al. (2021). Integrated analysis of multimodal single-cell data. Cell 184, 3573–3587 e3529. 10.1016/j.cell.2021.04.048.

117. Satija, R., Farrell, J.A., Gennert, D., Schier, A.F., and Regev, A. (2015). Spatial reconstruction of single-cell gene expression data. Nat Biotechnol 33, 495–502. 10.1038/nbt.3192.

118. Stuart, T., Butler, A., Hoffman, P., Hafemeister, C., Papalexi, E., Mauck, W.M., 3rd, Hao, Y., Stoeckius, M., Smibert, P., and Satija, R. (2019). Comprehensive Integration of Single-Cell Data. Cell 177, 1888–1902 e1821. 10.1016/j.cell.2019.05.031.

119. Korsunsky, I., Millard, N., Fan, J., Slowikowski, K., Zhang, F., Wei, K., Baglaenko, Y., Brenner, M., Loh, P.R., and Raychaudhuri, S. (2019). Fast, sensitive and accurate integration of single-cell data with Harmony. Nat Methods 16, 1289–1296. 10.1038/s41592-019-0619-0.

120. Chen, E.Y., Tan, C.M., Kou, Y., Duan, Q., Wang, Z., Meirelles, G.V., Clark, N.R., and Ma’ayan, A. (2013). Enrichr: interactive and collaborative HTML5 gene list enrichment analysis tool. BMC Bioinformatics 14, 128. 10.1186/1471-2105-14-128.

121. Kuleshov, M.V., Jones, M.R., Rouillard, A.D., Fernandez, N.F., Duan, Q., Wang, Z., Koplev, S., Jenkins, S.L., Jagodnik, K.M., Lachmann, A., et al. (2016). Enrichr: a comprehensive gene set enrichment analysis web server 2016 update. Nucleic Acids Res 44, W90–97. 10.1093/nar/gkw377.

122. Speir, M.L., Bhaduri, A., Markov, N.S., Moreno, P., Nowakowski, T.J., Papatheodorou, I., Pollen, A.A., Raney, B.J., Seninge, L., Kent, W.J., and Haeussler, M. (2021). UCSC Cell Browser: visualize your single-cell data. Bioinformatics 37, 4578–4580. 10.1093/bioinformatics/btab503.

