## Supplemental Materials for "Endocardial primary cilia and blood flow are required for regulation of EndoMT during endocardial cushion development"

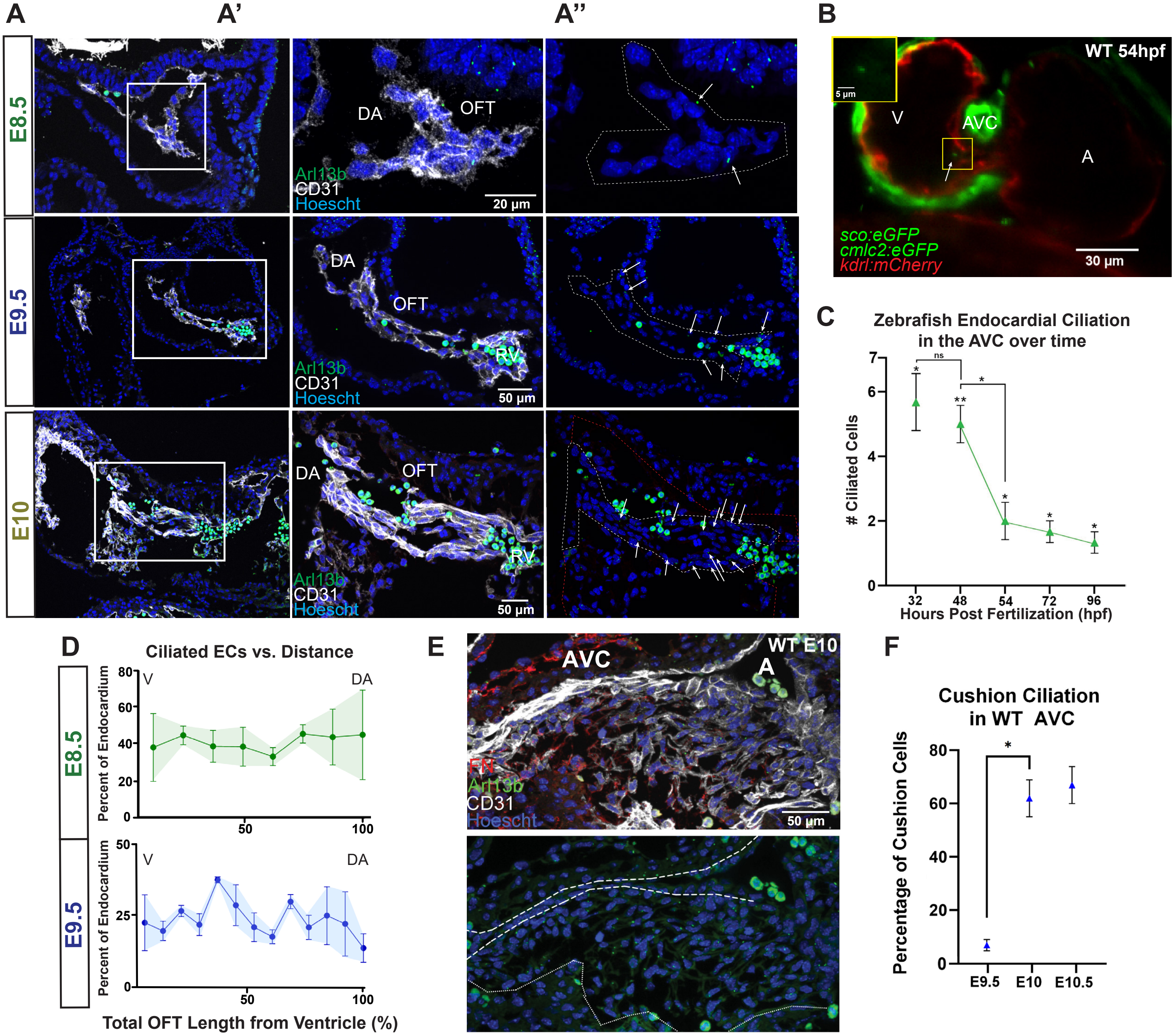

A
A'
A''
B
E8.5
E9.5
E10
DA
OFT
RV
Arl13b
CD31
Hoescht
20 µm
50 µm
WT 54hpf
V
AVC
A
5 µm
sco:eGFP
cmlc2:eGFP
kdrl:mCherry
30 µm
C
Zebrafish Endocardial Ciliation in the AVC over time
### Ciliated Cells
Hours Post Fertilization (hpf)
32
48
54
72
96
ns
D
Ciliated ECs vs. Distance
Percent of Endocardium
Total OFT Length from Ventricle (%)
E
WT E10
FN
F
Cushion Ciliation in WT AVC
Percentage of Cushion Cells
E9.5
E10
E10.5

**Figure S1. Endocardial ciliation of the ECCs is spatiotemporally dynamic during cushion EndoMT and is conserved in zebrafish**

**A)** Immunofluorescence on mouse embryo sections for cilia (ARL13B, green) on endocardial cells (CD31, white) of the OFT over cushion development (e8.5, e9.5, e10). Nuclei are shown in blue (Hoescht). Close ups **A')** with and **A'')** without CD31; white arrows indicate cilia; luminal endocardial cells outlined with white and whole OFT (including cushion mesenchyme) outlined in red. **B)** Still of live 54hpf tg(Arl13b:eGPF;Kdrl (Flk-1):mCherry;Cmlc2:eGFP) zebrafish heart showing cilia and myocardium in green and endocardium in red. Closeup of cilia provided in yellow box. **C)** Zebrafish endocardial ciliation over time in the atrioventricular canal (n=3). **D)** Ciliated endocardial cells as percentage of all endocardial cells versus distance in the mouse OFT at e8.5 (n=2) and e9.5 (n=3). Distance runs left to right, Ventricle to Dorsal Aorta. **E)** Immunofluorescence on e10 mouse embryo sections for cilia (ARL13B, green), endocardial cells (CD31, white) and mesenchyme (FN, red) of the AVC. Nuclei are shown in blue (Hoescht). Nuclei and cilia are shown alone with the luminal endocardial cells outlined with dashed white and whole AVC (including cushion mesenchyme) outlined in dotted white. **F)** Number of ciliated cushion cells as a percentage of total cushion cells in the AVC over time (e9.5 (n=3), e10 (n=3), e10.5 (n=3)). Statistics: ns ($p > 0.05$), * ($p \leq 0.05$), ** ($p \leq 0.01$). Data are represented as mean ± SEM. Abbreviations: A-Atrium, AVC-Atrioventricular Canal, EC-Endocardial Cell, V-Ventricle, OFT-Outflow Tract, WT-Wildtype.

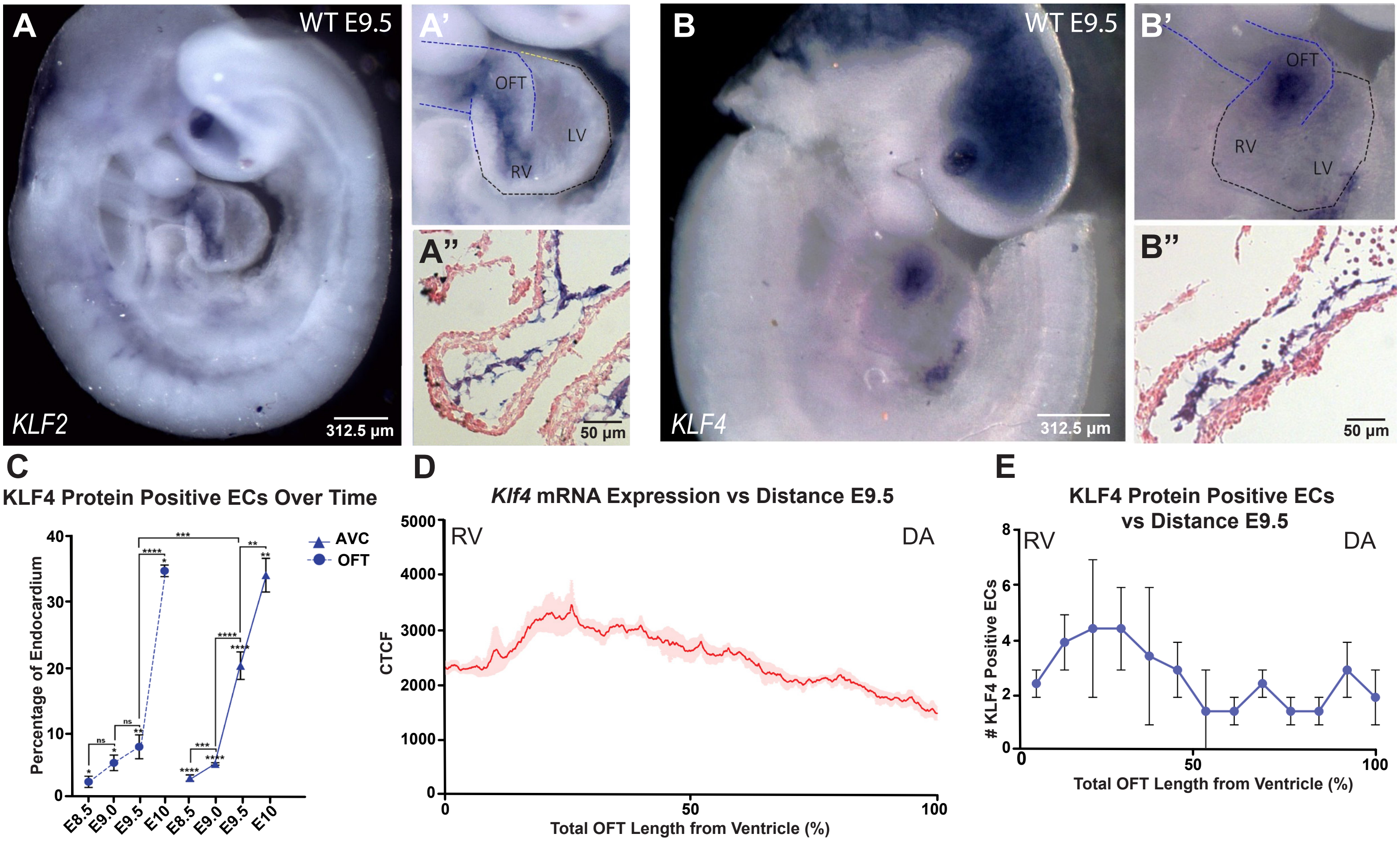

A
WT E9.5
KLF2
312.5 μm
A'
OFT
LV
RV
A"
50 μm
B
WT E9.5
KLF4
312.5 μm
B'
OFT
RV
LV
B"
50 μm
C
KLF4 Protein Positive ECs Over Time
Percentage of Endocardium
AVC
OFT
E8.5
E9.0
E9.5
E10
D
*Klf4* mRNA Expression vs Distance E9.5
CTCF
RV
DA
Total OFT Length from Ventricle (%)
E
KLF4 Protein Positive ECs vs Distance E9.5
### KLF4 Positive ECs
RV
DA
Total OFT Length from Ventricle (%)

**Figure S2. The mechanosensitive transcription factors KLF2 and KLF4 are dynamically expressed in mouse endocardium during cushion EndoMT**

Whole mount ISH of e9.5 mouse embryos for **A)** *Klf2* mRNA and **B)** *Klf4* mRNA with closeup **(A', B')** and cryosection **(A", B")** of the OFT. **C)** KLF4 protein positive endocardial cells over time as percentage of all endocardial cells in the OFT (e8.5 (n=11), e9 (n=3), e9.5 (n=6), e10 (n=3)) and AVC (e8.5 (n=14), e9 (n=6), e9.5 (n=8), e10 (n=3)). **D)** mRNA expression as measured by Grey_Value over distance for *Klf4* (n=3) in the e9.5 OFT. **E)** KLF4 protein positive endocardial cells versus distance in the e9.5 OFT (n=3). For D) and E), distance runs left to right, Atrium to Left Ventricle. *Statistics*: ns ($p > 0.05$), * ($p \leq 0.05$), ** ($p \leq 0.01$), **** ($p \leq 0.0001$). Data are represented as mean ± SEM. Abbreviations: DA-Dorsal Aorta, EC-Endocardial Cell, LV-Left Ventricle, OFT-Outflow Tract, RV-Right Ventricle, WT-Wildtype.

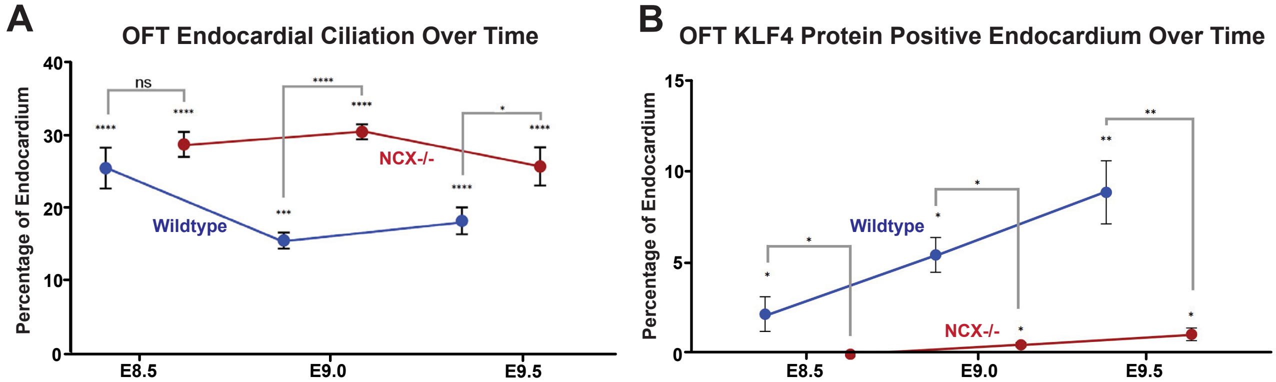
A
OFT Endocardial Ciliation Over Time
Percentage of Endocardium
40
30
20
10
0
E8.5
E9.0
E9.5
ns
****
***
*
Wildtype
NCX-/-
B
OFT KLF4 Protein Positive Endocardium Over Time
Percentage of Endocardium
15
10
5
0
E8.5
E9.0
E9.5
*
**
Wildtype
NCX-/-

**Figure S3. KLF4 expression at the OFT correlates with endocardial ciliation in a blood flow dependent manner**

**A)** Endocardial ciliation in the OFT over time as percentage of all endocardial cells in wildtype mice (blue) and *Ncx1*$^{-/-}$ mice (red) (e8.5 (WT n=7, *Ncx1*$^{-/-}$ n=7), e9.0 (WT n=4, *Ncx1*$^{-/-}$ n=4), e9.5 (WT n=7, *Ncx1*$^{-/-}$ n=7)). **B)** KLF4 protein positive endocardial cells in the OFT over time as percentage of all endocardial cells in wildtype mice (blue) and *Ncx1*$^{-/-}$ mice (red) (e8.5 (WT n=10, *Ncx1*$^{-/-}$ n=10), e9.0 (WT n=4, *Ncx1*$^{-/-}$ n=4), e9.5 (WT n=5, *Ncx1*$^{-/-}$ n=5)). *Statistics*: ns ($p > 0.05$), * ($p \leq 0.05$), ** ($p \leq 0.01$), *** ($p \leq 0.001$), **** ($p \leq 0.0001$). Data are represented as mean ± SEM. Abbreviations: OFT-Outflow Tract, WT-Wildtype.

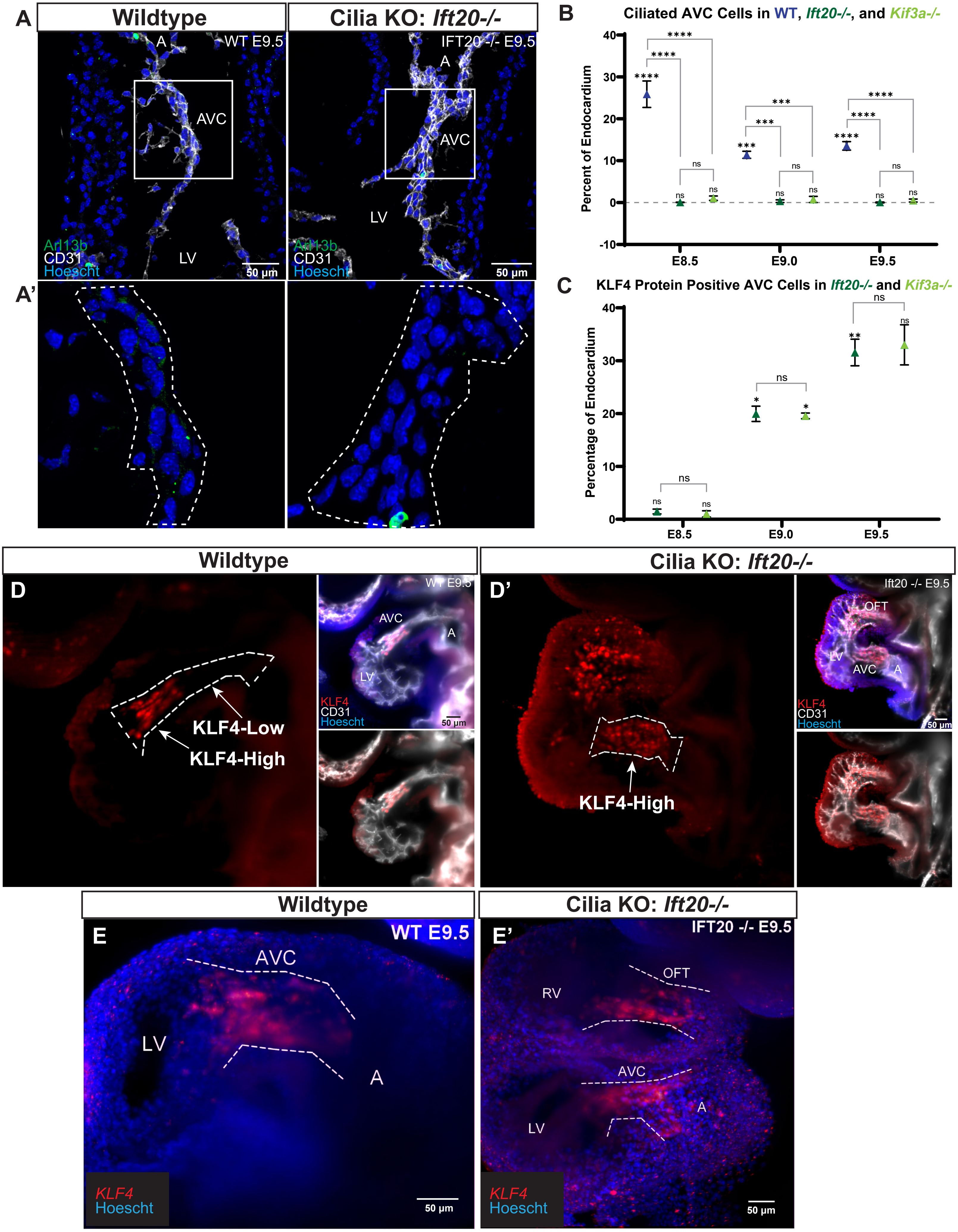

A
Wildtype
Cilia KO: Ift20-/-
WT E9.5
IFT20 -/- E9.5
A
AVC
LV
Arl13b
CD31
Hoescht
50 µm
A'
B
Ciliated AVC Cells in WT, Ift20-/-, and Kif3a-/-
Percent of Endocardium
E8.5
E9.0
E9.5
C
KLF4 Protein Positive AVC Cells in Ift20-/- and Kif3a-/-
Percentage of Endocardium
Wildtype
Cilia KO: Ift20-/-
D
D'
WT E9.5
Ift20 -/- E9.5
AVC
A
LV
OFT
KLF4-Low
KLF4-High
KLF4
CD31
Hoescht
50 µm
Wildtype
Cilia KO: Ift20-/-
E
E'
WT E9.5
IFT20 -/- E9.5
AVC
LV
A
RV
OFT
KLF4
Hoescht
50 µm

**Figure S4. Failure of ciliogenesis results in abnormal KLF4 expression**

**A)** Immunofluorescence on e9.5 wildtype and cilia KO (*Ift20*$^{-/-}$) mouse heart sections for cilia (ARL13B, green) on endocardial cells (CD31, white). Nuclei are shown in blue (Hoescht). **A')** Closeup of boxed AVC region without CD31 signal; the AVC is outlined in white. **B)** Ciliated endocardial cells in the AVC over time as percentage of all endocardial cells in wildtype (blue), *Ift20*$^{-/-}$ (dark green) and *Kif3a*$^{-/-}$ (light green) (e8.5 (WT n=8, *Ift20*$^{-/-}$ n=6, *Kif3a*$^{-/-}$ n=4), e9.0 (WT n=4, *Ift20*$^{-/-}$ n=3, *Kif3a*$^{-/-}$ n=3), e9.5 (WT n=6, *Ift20*$^{-/-}$ n=3, *Kif3a*$^{-/-}$ n=4)). **C)** KLF4 protein positive endocardial cells in the AVC over time as percentage of all endocardial cells in *Ift20*$^{-/-}$ (dark green) and *Kif3a*$^{-/-}$ (light green) (e8.5 (*Ift20*$^{-/-}$ n=4, *Kif3a*$^{-/-}$ n=3), e9.0 (*Ift20*$^{-/-}$ n=2, *Kif3a*$^{-/-}$ n=2), e9.5 (*Ift20*$^{-/-}$ n=4, *Kif3a*$^{-/-}$ n=2)). Immunofluorescence on e9.5 **D)** wildtype and **E)** cilia KO (*Ift20*$^{-/-}$) whole mount hearts for KLF4 (red) in endocardial cells (CD31, white). Nuclei are shown in blue (Hoescht); the AVC is outlined in white. HCR-FISH on whole mount **F)** wildtype and **G)** Cilia KO mouse embryos at e9.5 showing *Klf4* mRNA (red) and nuclei (Hoescht, blue). Endocardial regions are outlined in white. *Statistics*: ns ($p > 0.05$), * ($p \leq 0.05$), ** ($p \leq 0.01$), *** ($p \leq 0.001$), **** ($p \leq 0.0001$). Data are represented as mean ± SEM. Abbreviations: A-Atrium, AVC-Atrioventricular Canal, LV-Left Ventricle, OFT-Outflow Tract, RV-Right Ventricle, WT-Wildtype.

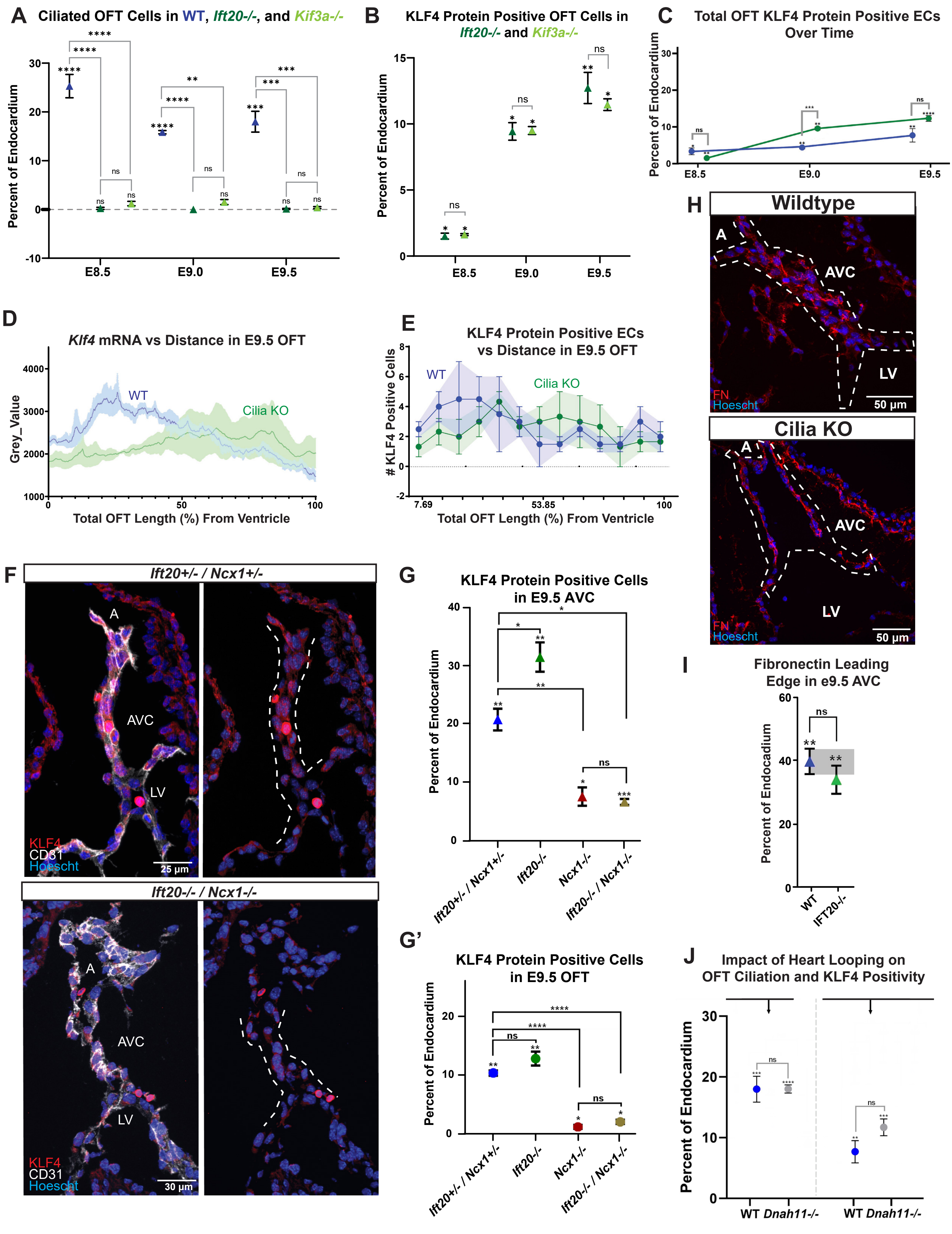

A
Ciliated OFT Cells in WT, Ift20-/-, and Kif3a-/-
Percent of Endocardium
E8.5
E9.0
E9.5
B
KLF4 Protein Positive OFT Cells in Ift20-/- and Kif3a-/-
Percent of Endocardium
C
Total OFT KLF4 Protein Positive ECs Over Time
Percent of Endocardium
D
Klf4 mRNA vs Distance in E9.5 OFT
Grey_Value
WT
Cilia KO
Total OFT Length (%) From Ventricle
E
KLF4 Protein Positive ECs vs Distance in E9.5 OFT
### KLF4 Positive Cells
WT
Cilia KO
Total OFT Length (%) From Ventricle
F
Ift20+/- / Ncx1+/-
A
AVC
LV
KLF4
CD31
Hoescht
25 µm
Ift20-/- / Ncx1-/-
A
AVC
LV
KLF4
CD31
Hoescht
30 µm
G
KLF4 Protein Positive Cells in E9.5 AVC
Percent of Endocardium
Ift20+/- / Ncx1+/-
Ift20-/-
Ncx1-/-
Ift20-/- / Ncx1-/-
G'
KLF4 Protein Positive Cells in E9.5 OFT
Percent of Endocardium
Ift20+/- / Ncx1+/-
Ift20-/-
Ncx1-/-
Ift20-/- / Ncx1-/-
H
Wildtype
A
AVC
LV
FN
Hoescht
50 µm
Cilia KO
A
AVC
LV
FN
Hoescht
50 µm
I
Fibronectin Leading Edge in e9.5 AVC
Percent of Endocadium
WT
IFT20-/-
J
Impact of Heart Looping on OFT Ciliation and KLF4 Positivity
Percent of Endocardium
WT Dnah11-/-
WT Dnah11-/-

**Figure S5. Failure of ciliogenesis results in abnormal KLF4 expression in a flow-dependent manner**

**A)** Ciliated endocardial cells in the OFT over time as percentage of all endocardial cells in wildtype (blue), *Ift20*$^{-/-}$ (dark green) and *Kif3a*$^{-/-}$ (light green) (e8.5 (WT n=8, *Ift20*$^{-/-}$ n=4, *Kif3a*$^{-/-}$ n=4), e9.0 (WT n=4, *Ift20*$^{-/-}$ n=2, *Kif3a*$^{-/-}$ n=2), e9.5 (WT n=6, *Ift20*$^{-/-}$ n=4, *Kif3a*$^{-/-}$ n=3)). **B)** KLF4 protein positive endocardial cells in the OFT over time as percentage of all endocardial cells in *Ift20*$^{-/-}$ (dark green) and *Kif3a*$^{-/-}$ (light green) (e8.5 (*Ift20*$^{-/-}$ n=3, *Kif3a*$^{-/-}$ n=2), e9.0 (*Ift20*$^{-/-}$ n=2, *Kif3a*$^{-/-}$ n=2), e9.5 (*Ift20*$^{-/-}$ n=4, *Kif3a*$^{-/-}$ n=2)). **C)** KLF4 protein positive endocardial cells in the OFT over time as percentage of all endocardial cells in wildtype (blue) and cilia KO mice (green) (e8.5 (WT n=4, cilia KO n=4), e9.0 (WT n=3, cilia KO n=3), e9.5 (WT n=6, cilia KO n=6)). **D)** *Klf4* mRNA expression as measured by Grey_Value over distance in the OFT of e9.5 cilia KO mice (green, n=3); wildtype is given for comparison in blue (n=3). **E)** Number of KLF4 protein positive endocardial cells versus distance in the OFT of e9.5 cilia KO mice (green, n=3); wildtype is given for comparison in blue (n=3). **F)** Immunofluorescence on e9.5 sections of *Ift2*$^{+/-}$*/ Ncx1*$^{+/-}$ and *Ift20*$^{-/-}$ */ Ncx1*$^{-/-}$ hearts for KLF4 (red) in endocardial cells (CD31, white). Nuclei are shown in blue (Hoescht); the AVC is outlined in white. KLF4 protein positive endocardial cells in the e9.5 **G)** AVC and **G')** OFT as percentage of all endocardial cells in *Ift2*$^{+/-}$*/ Ncx1*$^{+/-}$ (blue, AVC n=3, OFT n=3) and *Ift20*$^{-/-}$*/ Ncx1*$^{-/-}$ (brown, AVC n=4, OFT n=4). *Ift20*$^{-/-}$ (green, AVC n=4, OFT n=4) and *Ncx1*$^{-/-}$ (red, AVC n=4, OFT n=5) are given for comparison. **H)** Immunofluorescence on e9.5 wildtype and cilia KO AVC sections for Fibronectin (red). Nuclei are shown in blue (Hoescht); AVC is outlined in white. **I)** Endocardial cells with a leading edge of fibronectin as percentage of all endocardial cells in e9.5 AVCs of wildtype (blue, n=4) and cilia KO (green, n=5) mice. **J)** Ciliated endocardial cells and KLF4 protein positive endocardial cells in the OFT as percentage of all endocardial cells in e9.5 wildtype (blue) and *Dnah11*$^{-/-}$ (grey) mice (cilia: WT n=6, *Dnah11*$^{-/-}$ n=6; KLF4: WT n=6, *Dnah11*$^{-/-}$ n=6). For D) and E), distance runs left to right, Right Ventricle to Dorsal Aorta. *Statistics*: ns ($p > 0.05$), * ($p \leq 0.05$), ** ($p \leq 0.01$), *** ($p \leq 0.001$), **** ($p \leq 0.0001$). Data are represented as mean ± SEM. Abbreviations: A-Atrium, EC-Endocardial Cell, OFT-Outflow Tract, RV-Right Ventricle, WT-Wildtype.

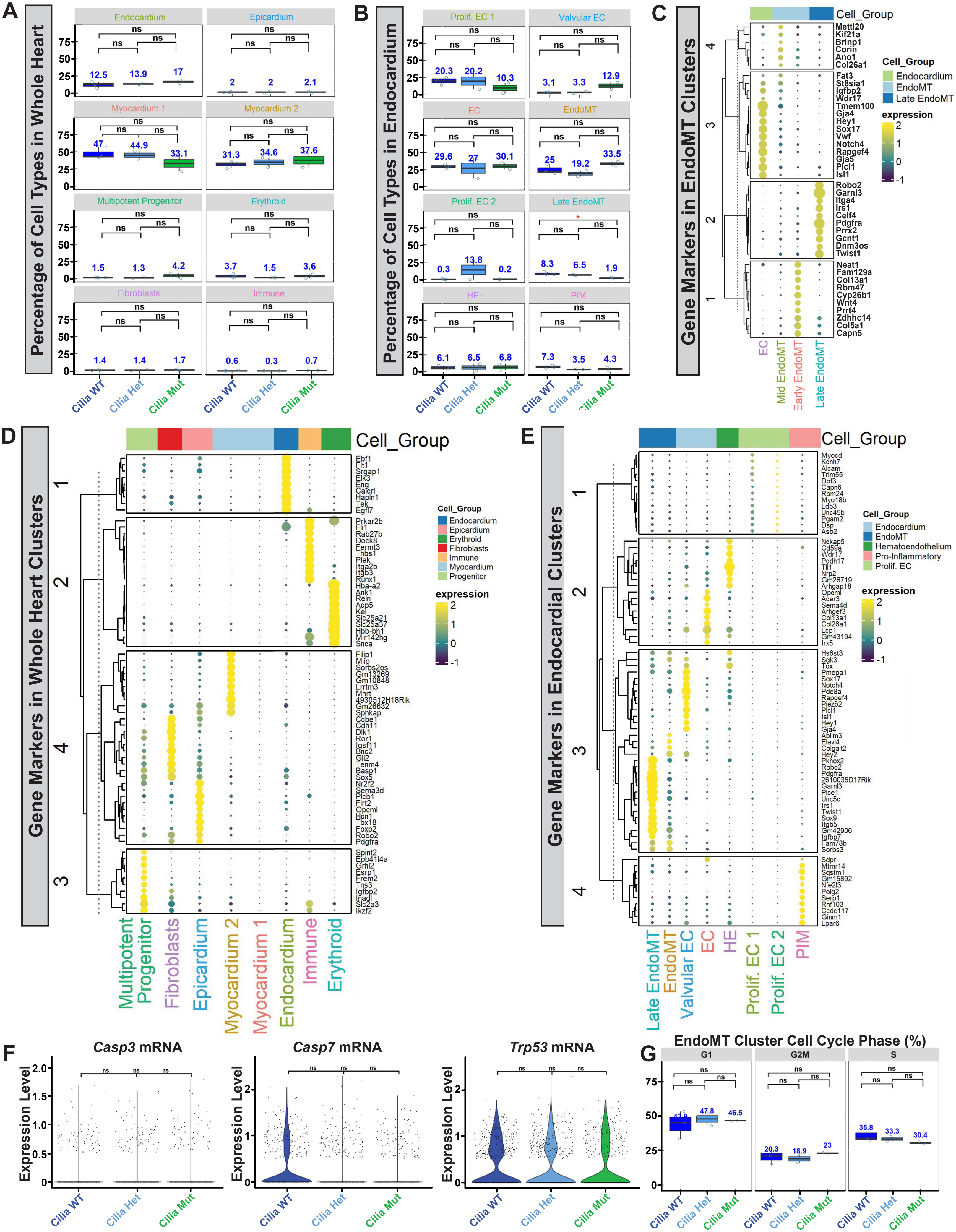
A
Percentage of Cell Types in Whole Heart
Endocardium
Epicardium
Myocardium 1
Myocardium 2
Multipotent Progenitor
Erythroid
Fibroblasts
Immune
12.5
13.9
17
2
2
2.1
47
44.9
33.1
31.3
34.6
37.6
1.5
1.3
4.2
3.7
1.5
3.6
1.4
1.4
1.7
0.6
0.3
0.7
ns
Cilia WT
Cilia Het
Cilia Mut
B
Percentage of Cell Types in Endocardium
Prolif. EC 1
Valvular EC
EC
EndoMT
Prolif. EC 2
Late EndoMT
HE
PIM
20.3
20.2
10.3
3.1
3.3
12.9
29.6
27
30.1
25
19.2
33.5
0.3
13.8
0.2
8.3
6.5
1.9
6.1
6.5
6.8
7.3
3.5
4.3
*
C
Gene Markers in EndoMT Clusters
Cell_Group
Endocardium
EndoMT
Late EndoMT
expression
Mettl20
Kif21a
Brinp1
Corin
Ano1
Col26a1
Fat3
St8sia1
Igfbp2
Wdr17
Tmem100
Gja4
Hey1
Sox17
Vwf
Notch4
Rapgef4
Gja5
Plcl1
Isl1
Robo2
Garnl3
Itga4
Irs1
Celf4
Pdgfra
Prrx2
Gcnt1
Dnm3os
Twist1
Neat1
Fam129a
Col13a1
Rbm47
Cyp26b1
Wnt4
Prrt4
Zdhhc14
Col5a1
Capn5
EC
Mid EndoMT
Early EndoMT
Late EndoMT
D
Gene Markers in Whole Heart Clusters
Cell_Group
Endocardium
Epicardium
Erythroid
Fibroblasts
Immune
Myocardium
Progenitor
expression
Ebf1
Flt1
Srgap1
Elk3
Eng
Calcrl
Hapln1
Tek
Egfl7
Prkar2b
Fli1
Rab27b
Dock8
Fermt3
Thbs1
Plek
Itga2b
Itgb3
Runx1
Hba-a2
Ank1
Reln
Acp5
Kel
Slc25a21
Slc25a37
Hbb-bh1
Mir142hg
Snca
Filip1
Mlip
Sorbs2os
Gm13269
Gm10848
Lrrtm3
Mhrt
4930512H18Rik
Gm26632
Sphkap
Ccbe1
Cdh11
Dlk1
Ror1
Igsf11
Bnc2
Gli2
Tenm4
Basp1
Sox5
Nr2f2
Sema3d
Plcb1
Flrt2
Opcml
Hcn1
Tbx18
Foxp2
Robo2
Pdgfra
Spint2
Epb41l4a
Grhl2
Esrp1
Frem2
Tns3
Igfbp2
Inadl
Slc2a3
Ikzf2
Multipotent Progenitor
Fibroblasts
Epicardium
Myocardium 2
Myocardium 1
Endocardium
Immune
Erythroid
E
Gene Markers in Endocardial Clusters
Cell_Group
Endocardium
EndoMT
Hematoendothelium
Pro-Inflammatory
Prolif. EC
expression
Myocd
Kcnh7
Alcam
Trim55
Dpf3
Capn6
Rbm24
Myo18b
Ldb3
Unc45b
Pgam2
Dsp
Asb2
Nckap5
Cd59a
Wdr17
Pcdh17
Tll1
Nrp2
Gm26719
Arhgap18
Opcml
Acer3
Sema4d
Arhgef3
Col13a1
Col26a1
Lcp1
Gm43194
Irx5
Hs6st3
Sgk3
Tox
Pmepa1
Sox17
Notch4
Pde8a
Rapgef4
Piezo2
Plcl1
Isl1
Hey1
Gja4
Ablim3
Elavl4
Colgalt2
Hey2
Pknox2
Robo2
Pdgfra
2610035D17Rik
Garnl3
Plce1
Unc5c
Irs1
Twist1
Sox9
Itgb5
Gm42906
Igfbp7
Fam78b
Sorbs3
Sdpr
Mtmr14
Sqstm1
Gm15892
Nfe2l3
Polg2
Serp1
Rnf103
Ccdc117
Ginm1
Lpar6
Late EndoMT
EndoMT
Valvular EC
EC
HE
Prolif. EC 1
Prolif. EC 2
PIM
F
Casp3 mRNA
Casp7 mRNA
Trp53 mRNA
Expression Level
G
EndoMT Cluster Cell Cycle Phase (%)
G1
G2M
S
47.8
46.5
20.3
18.9
23
35.8
33.3
30.4

**Figure S6. Defects in ciliogenesis block EndoMT**

Graphs of the percentage of cell types in the **A)** whole heart and **B)** endocardium from snRNA-seq data for each genotype (Replicates: Cilia WT n=3, Cilia Het n=2, Cilia Mut n=2). Statistically significant numbers are given in red. Dotplots for top DEGs for each cell type in the **C)** EndoMT cluster, **D)** whole heart and **F)** endocardium from snRNA-seq data. **G)** Violin plots for proliferation gene expression in each genotype. **H)** Percentage of cells in each cell cycle phase for each genotype in the snRNA-seq EndoMT cluster (Replicates: Cilia WT n=3, Cilia Het n=2, Cilia Mut n=2). *Statistics*: ns ($p > 0.05$), ** ($p \leq 0.01$). Data are represented as mean ± SEM. Abbreviations: A-Atrium, DEG-Differentially Expressed Gene, EC-Endocardial Cell, HE-Hematoendothelium, LV-Left Ventricle, PIM-Pro-inflammatory, WT-Wildtype.

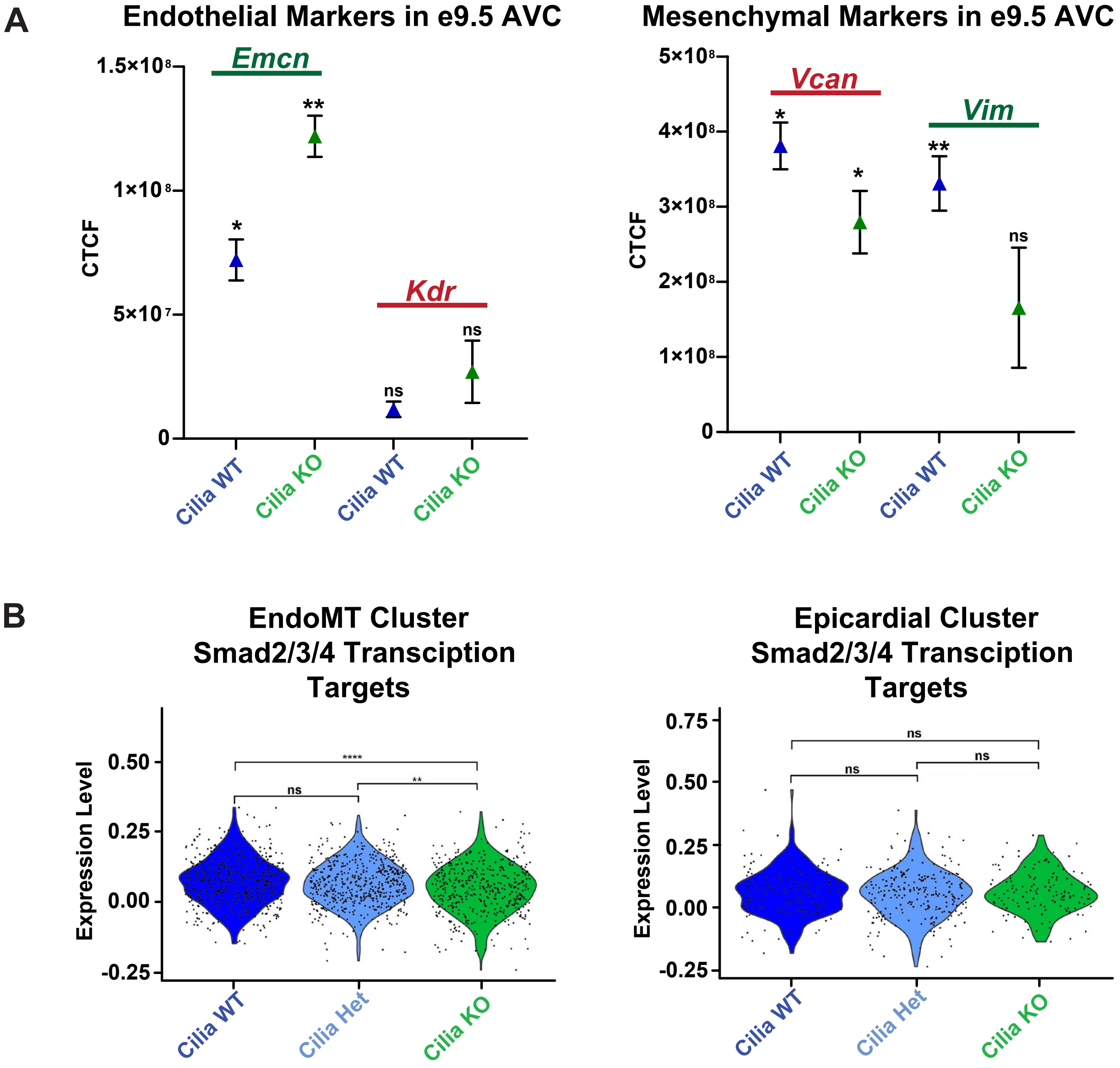
A
Endothelial Markers in e9.5 AVC
Emcn
Kdr
CTCF
Cilia WT
Cilia KO
Cilia WT
Cilia KO
Mesenchymal Markers in e9.5 AVC
Vcan
Vim
CTCF
Cilia WT
Cilia KO
Cilia WT
Cilia KO
B
EndoMT Cluster
Smad2/3/4 Transcription
Targets
Expression Level
Cilia WT
Cilia Het
Cilia KO
Epicardial Cluster
Smad2/3/4 Transcription
Targets
Expression Level
Cilia WT
Cilia Het
Cilia KO

**Figure S7. Known EndoMT molecular pathways are dysregulated in Cilia KO hearts**

**A)** Corrected total cell fluorescence (CTCF) analysis of HCR-FISH for *Emcn, Kdr, Vcan, and Vim* in the e9.5 AVC (WT n=3, Cilia KO n=3). **B)** Violin plots for GeneSet Enrichment of Smad2/3/4 transcriptional targets (GSEA-MSigDB:MM14876) in each genotype for the EndoMT cluster or the Epicardial Cluster. *Statistics*: ns ($p > 0.05$), *** ($p \leq 0.001$), **** ($p \leq 0.0001$). Abbreviations: WT-Wildtype.

#### Supplemental Table Legends:

**Table S1: KLF2 and KLF4 variants in human HLHS**

From Sierant et al. (in prep).

**Table S2: Differentially expressed genes between each pseudo-phase of EndoMT**

A) Quiescent Endocardium. B) Early EndoMT. C) Mid EndoMT. D) Late EndoMT. Generated from scRNA-seq[28] and snRNA-seq. Avg_logFC is given for the named cluster compared to the other three.

**Table S3: Differentially Expressed Genes between *Ift20*$^{+/+}$, *Ift20*$^{+/-}$ and *Ift20*$^{-/-}$ endocardium undergoing EndoMT**

A cutoff of 0.4 avg_logFC in *Ift20*$^{+/+}$ vs *Ift20*$^{-/-}$ was used to guarantee differential expression.

**Table S4: Disorganized Gene Expression in *Ift20*$^{-/-}$ endocardium between each pseudo-phase of EndoMT**

Genes that are significantly either increased or decreased between each pseudo-phase in *Ift20*$^{+/+}$ and *Ift20*$^{+/-}$ but not *Ift20*$^{-/-}$.

**STAR Methods: Key Resources Table**

| **Table S1** | KLF2/KLF4 Interactome | | | | | |
|---|---|---|---|---|---|---|
| HLHS, N=475 | Observed | | Expected | | Enrichment | p_value |
| | N | Rate | N | Rate | | |
| Total | 4 | 8.42E-03 | 0.7 | 1.47E-03 | 6 | 4.84E-03 |
| Syn | 0 | 0 | NA | NA | 0 | NA |
| D-Mis | 2 | 4.21E-03 | 0.1 | 2.11E-04 | 14.8 | 8.31E-03 |
| Mis | 2 | 4.21E-03 | 0.4 | 8.42E-04 | 4.84 | 0.07 |
| LoF | 2 | 4.21E-03 | 0.1 | 2.11E-04 | 32.9 | 1.78E-01 |
| Damaging | 4 | 8.42E-03 | 0.2 | 4.21E-04 | 20.4 | 5.23E-05 |

| Table S2A | Quiescent Endocardium | | | | | | | |
|---|---|---|---|---|---|---|---|---|
| Gene Name | avg_logFC | p_val_adj | Gene Name | avg_logFC | p_val_adj | Gene Name | avg_logFC | p_val_adj |
| Tmem100 | 2.203237 | 1.35E-64 | Tox1 | 1.106295 | 7.94E-12 | Pik3c2b1 | 0.762216 | 0.000265 |
| Rgs2 | 0.808759 | 5.52E-50 | Ptprm1 | 0.706502 | 2.22E-11 | Gm20631 | 0.350515 | 0.000276 |
| Gja4 | 1.43244 | 1.76E-49 | Eva1a | 0.757828 | 3.08E-11 | Card10 | 0.416312 | 0.000278 |
| Myct1 | 0.949421 | 4.85E-48 | Ushbp1 | 0.646227 | 3.38E-11 | Dock61 | 0.53802 | 0.000279 |
| Gja5 | 1.168792 | 2.16E-47 | Pmepa1 | 0.914887 | 6.91E-11 | Inpp5d | 0.642727 | 0.000297 |
| Prdm16 | 1.124404 | 1.72E-43 | Epas1 | 0.719116 | 1.16E-10 | St8sia4 | 0.269929 | 0.000321 |
| Acvrl1 | 0.832456 | 7.24E-43 | Csrp21 | 0.76951 | 1.44E-10 | Plcb1 | 0.858723 | 0.000325 |
| Tes | 0.857196 | 3.79E-39 | Prkaa2 | 0.562159 | 1.62E-10 | Cd40 | 0.515846 | 0.000367 |
| Vwf | 1.57904 | 4.33E-39 | Hmcn11 | 1.106647 | 3.39E-10 | Mkl21 | 0.57033 | 0.00039 |
| Rapgef4 | 1.733555 | 8.22E-36 | Mast41 | 1.101307 | 3.66E-10 | Tspan2 | 0.657846 | 0.00041 |
| Zfp69 | 1.713838 | 1.06E-32 | Efna1 | 0.877403 | 5.69E-10 | Myo10 | 0.678631 | 0.00042 |
| Slco3a1 | 2.075907 | 1.43E-32 | Irf2 | 0.879424 | 8.50E-10 | Frmd4b1 | 0.554759 | 0.000544 |
| Slc22a23 | 1.873085 | 2.08E-32 | Gm14507 | 0.447858 | 1.17E-09 | Igfbp2 | 0.318485 | 0.00062 |
| Isl1 | 1.241783 | 3.41E-32 | Ramp2 | 0.914268 | 1.25E-09 | Esam | 0.552377 | 0.00068 |
| Hey1 | 1.0768 | 9.69E-32 | Unc13b | 0.94826 | 1.37E-09 | Bmpr2 | 0.571726 | 0.000691 |
| Emcn1 | 2.015388 | 2.19E-28 | Tgfbr31 | 0.916157 | 2.16E-09 | Zfp711 | 0.591322 | 0.000823 |
| Arhgef28 | 1.649687 | 3.40E-28 | Piezo21 | 0.7774 | 2.29E-09 | Smurf2 | 0.785184 | 0.000828 |
| Usp43 | 0.73846 | 1.95E-26 | Synpo | 0.853066 | 4.92E-09 | Disc1 | 0.679008 | 0.000915 |
| Prex21 | 1.42731 | 3.37E-26 | Chn1 | 0.837305 | 5.24E-09 | Junb | 0.534123 | 0.001029 |
| Fam124b | 0.461875 | 5.98E-26 | Eya1 | 0.558966 | 7.93E-09 | Adgre5 | 0.549786 | 0.001044 |
| Nrp11 | 1.726345 | 6.13E-26 | Nxpe4 | 0.406627 | 1.76E-08 | Ddx26b | 0.69107 | 0.001052 |
| Slit3 | 2.147417 | 1.25E-23 | Dysf1 | 0.771744 | 1.92E-08 | Cd109 | 0.580562 | 0.001081 |
| Cyp26a1 | 0.52743 | 2.26E-23 | Prickle21 | 0.896921 | 5.21E-08 | Kcnc2 | 0.449978 | 0.001111 |
| Adam121 | 1.89887 | 8.23E-23 | Tjp1 | 0.646035 | 5.83E-08 | Plxnd11 | 0.672214 | 0.001169 |
| Notch4 | 0.881153 | 8.37E-22 | Vegfc | 1.168827 | 6.05E-08 | Ctsh | 0.492318 | 0.001223 |
| Mfng | 0.581894 | 1.06E-21 | Npr1 | 0.567569 | 6.45E-08 | Tbc1d1 | 0.759522 | 0.001234 |
| Tll11 | 1.419156 | 1.15E-21 | Dab2 | 0.910906 | 8.01E-08 | Sncaip2 | 0.701264 | 0.001431 |
| Ldb2 | 1.72162 | 2.80E-21 | 8030442B05Rik | 0.326315 | 1.22E-07 | Palld | 0.85224 | 0.001564 |
| St6gal11 | 1.392322 | 6.44E-21 | Tspan181 | 0.621196 | 1.50E-07 | Adgrl41 | 0.646056 | 0.001807 |
| Efnb2 | 1.312061 | 1.40E-20 | Sox18 | 0.724049 | 1.67E-07 | Gdpd5 | 0.649602 | 0.00214 |
| Egfl71 | 1.226943 | 1.09E-19 | Myzap | 0.466649 | 1.83E-07 | Nxn | 0.765374 | 0.002251 |
| Sat11 | 1.637985 | 1.27E-19 | Rprm | 0.306323 | 2.58E-07 | Arntl2 | 0.33826 | 0.002267 |
| Dll4 | 0.645392 | 4.94E-19 | Clec14a | 0.479912 | 4.13E-07 | Nbea1 | 0.430978 | 0.002515 |
| Mecom1 | 1.374594 | 5.17E-19 | Inpp4b1 | 0.948003 | 5.02E-07 | Adgrl3 | 0.772086 | 0.002572 |
| Sox17 | 1.113399 | 6.56E-19 | Nipal2 | 0.571416 | 5.09E-07 | Ahr1 | 0.653257 | 0.002643 |
| Pced1b | 1.454409 | 5.77E-18 | Sema3g | 0.586625 | 5.63E-07 | Trim46 | 0.326866 | 0.002965 |
| Exoc3l4 | 0.834858 | 6.28E-18 | Arap31 | 0.547583 | 7.36E-07 | Luzp1 | 0.601365 | 0.00298 |
| Gm43734 | 0.31204 | 1.19E-17 | Nhsl1 | 0.80455 | 9.34E-07 | Sema6d | 0.61161 | 0.003304 |
| Dach11 | 1.50231 | 1.65E-17 | Mamld1 | 0.639555 | 9.44E-07 | Ctnnd21 | 0.713424 | 0.003699 |
| Cdc42ep3 | 0.525266 | 2.30E-17 | Tie11 | 0.840795 | 1.16E-06 | Rasip1 | 0.535667 | 0.004028 |
| Zfp366 | 0.539172 | 2.88E-17 | Arhgap29 | 0.836065 | 1.54E-06 | Fmnl3 | 0.543942 | 0.004293 |
| Atxn1 | 1.291133 | 1.23E-16 | Pcdh71 | 0.920567 | 1.70E-06 | Pdlim5 | 0.709794 | 0.0047 |
| Mmrn2 | 1.020497 | 2.22E-16 | Slit21 | 0.691984 | 2.52E-06 | Ccm2l1 | 0.586731 | 0.00512 |
| Gm7580 | 0.501959 | 2.32E-16 | Ptk2 | 0.526004 | 2.55E-06 | Cnnm21 | 0.600265 | 0.005291 |
| Sost | 0.354392 | 3.49E-16 | Plxna4 | 0.728249 | 2.57E-06 | Orai1 | 0.48781 | 0.005512 |
| Trpm31 | 1.69945 | 3.90E-16 | Pde9a | 0.323347 | 2.67E-06 | Anxa3 | 0.557227 | 0.005521 |
| Pde8a | 1.256234 | 4.69E-16 | Slc43a2 | 0.654474 | 3.86E-06 | Ass1 | 0.326779 | 0.005853 |
| Ccdc60 | 0.941198 | 5.44E-16 | Pik3r3 | 0.703996 | 5.12E-06 | Arhgap27 | 0.479027 | 0.006451 |
| Mall | 0.309814 | 6.44E-16 | Khdrbs31 | 0.841501 | 7.06E-06 | Plcb4 | 0.613597 | 0.006767 |
| Hsd11b2 | 0.368612 | 6.63E-16 | Gng11 | 0.58878 | 8.34E-06 | Frmd4a2 | 0.644883 | 0.007177 |
| Lamc3 | 0.350997 | 7.13E-16 | Vegfa | 0.711558 | 9.72E-06 | Ppp1r9a1 | 0.520008 | 0.007263 |
| Ecscr | 0.825553 | 8.33E-16 | Dennd5b1 | 0.632534 | 9.78E-06 | Mmp2 | 0.518049 | 0.007304 |
| Zfpm21 | 1.214645 | 9.01E-16 | Exoc61 | 0.61374 | 1.01E-05 | 11-Sep | 0.588834 | 0.012028 |
| Ldlrad4 | 1.155756 | 9.61E-16 | Col4a21 | 0.723302 | 1.33E-05 | Fam163a1 | 0.602364 | 0.01263 |
| Gm4876 | 0.416888 | 1.50E-15 | Rbfox1 | 0.656981 | 1.88E-05 | Nod1 | 0.260912 | 0.01424 |
| Asap3 | 0.38376 | 2.17E-15 | Prickle1 | 0.750982 | 2.04E-05 | Myo18a | 0.567795 | 0.015583 |
| Flt11 | 1.049352 | 4.24E-15 | Slc6a6 | 0.87547 | 2.06E-05 | St6galnac31 | 0.443455 | 0.016346 |
| Abcg1 | 0.314408 | 4.67E-15 | Mef2c | 0.828278 | 2.38E-05 | Tcf4 | 0.534587 | 0.017575 |
| Ptpre | 0.640436 | 4.71E-15 | Kalrn | 0.769532 | 2.49E-05 | Nr6a1os1 | 0.695261 | 0.018927 |
| Gcnt2 | 0.880674 | 5.06E-15 | Nedd9 | 1.016657 | 3.43E-05 | Lpcat2 | 0.320611 | 0.020693 |
| Col23a11 | 1.643987 | 7.09E-15 | Bcar3 | 0.478977 | 4.07E-05 | Lyl1 | 0.30345 | 0.021374 |

| | | | | | | | | |
|---|---|---|---|---|---|---|---|---|
| Plcl1 | 1.04891 | 9.59E-15 | Plekhg1 | 0.683189 | 7.36E-05 | Lrrc8d | 0.577141 | 0.024121 |
| Cdh5 | 0.962351 | 1.95E-14 | Slc46a3 | 0.261314 | 7.59E-05 | Gata31 | 0.502421 | 0.024296 |
| Cdh21 | 1.224204 | 4.11E-14 | Zfp423 | 0.793076 | 8.56E-05 | Taok2 | 0.510514 | 0.025066 |
| Calcrl | 1.202321 | 5.11E-14 | 1810011O10Rik | 0.614062 | 8.96E-05 | Mtus1 | 0.460923 | 0.02507 |
| Rassf9 | 0.550986 | 2.10E-13 | Nxpe2 | 0.413988 | 9.74E-05 | Atp13a31 | 0.877319 | 0.026374 |
| Pon3 | 0.874771 | 2.11E-13 | Plekhg5 | 0.791722 | 0.000125 | Irak2 | 0.396325 | 0.027018 |
| Plvap | 0.9806 | 2.82E-13 | Ccnjl | 0.746439 | 0.000131 | Pon2 | 0.626741 | 0.027531 |
| Rapgef51 | 1.050744 | 3.01E-13 | C530008M17Rik | 0.681494 | 0.000155 | Prtg1 | 0.713022 | 0.027965 |
| Trpc61 | 1.170834 | 4.20E-13 | Kdr1 | 0.692035 | 0.000156 | Ago4 | 0.312618 | 0.030057 |
| Rab27b | 0.846446 | 6.21E-13 | Eogt | 0.618967 | 0.000165 | S1pr11 | 0.577623 | 0.031637 |
| Dnm3 | 0.594114 | 8.75E-13 | Rdx | 0.622484 | 0.000168 | Efna51 | 0.625977 | 0.035179 |
| Stmn2 | 0.948249 | 1.01E-12 | Dock91 | 0.530966 | 0.00017 | Fbxl7 | 0.526779 | 0.036034 |
| Lcp1 | 1.016993 | 1.17E-12 | Tm6sf1 | 0.599926 | 0.00017 | Dock8 | 0.559169 | 0.037883 |
| Rapgef4os1 | 0.674549 | 1.95E-12 | Stox2 | 0.747478 | 0.000177 | Pecam1 | 0.461043 | 0.038165 |
| Cyyr1 | 1.139464 | 2.78E-12 | Gab2 | 0.512917 | 0.000184 | Ss18 | 0.480551 | 0.039045 |
| Kcnq1 | 0.457638 | 3.96E-12 | Col4a11 | 0.682848 | 0.000184 | Gm27151 | 0.526269 | 0.039385 |
| Adora2a | 0.429769 | 4.03E-12 | Txnip | 0.613559 | 0.000205 | Alpk3 | 0.537268 | 0.04056 |
| Fli1 | 0.920757 | 4.34E-12 | Lifr | 0.621064 | 0.000213 | Stab1 | 0.725554 | 0.042922 |
| Car8 | 1.049448 | 5.12E-12 | Bcl6b | 0.589293 | 0.000218 | Cmklr1 | 0.284537 | 0.043261 |
| Lama3 | 0.875109 | 6.23E-12 | | | | | | |

| **Table S2B** | Early EndoMT | | | | | | | |
|---|---|---|---|---|---|---|---|---|
| Gene Name | avg_logFC | p_val_adj | Gene Name | avg_logFC | p_val_adj | Gene Name | avg_logFC | p_val_adj |
| Ccdc141 | 2.341964 | 1.43E-79 | Cobll1 | 0.605149 | 1.77E-11 | Add3 | 0.411701 | 1.53E-05 |
| Tek | 1.222403 | 6.41E-68 | Hey2 | 0.517891 | 1.97E-11 | Egfl7 | 0.296124 | 1.91E-05 |
| Erg | 1.161378 | 1.36E-51 | Stard8 | 0.351668 | 2.26E-11 | Sh3bp4 | 0.379965 | 2.07E-05 |
| Tmem108 | 1.285604 | 1.19E-48 | Igf2 | 0.481855 | 2.31E-11 | Zc3h7a | 0.368258 | 2.19E-05 |
| Eml4 | 0.976775 | 6.08E-48 | Sorbs3 | 0.497918 | 4.00E-11 | Itga5 | 0.375188 | 2.36E-05 |
| Shank3 | 1.168677 | 1.67E-47 | H19 | 0.467329 | 5.48E-11 | Gls | 0.482651 | 2.82E-05 |
| Prkch | 1.076146 | 1.03E-46 | Ccser1 | 1.037834 | 7.26E-11 | Hdgf | 0.339474 | 2.95E-05 |
| Nfatc1 | 1.08397 | 1.18E-44 | Nipal3 | 0.339919 | 8.34E-11 | F11r | 0.327855 | 3.08E-05 |
| Atp2b4 | 1.161103 | 5.98E-43 | Ptprj | 0.636016 | 1.14E-10 | Kat2b | 0.392013 | 3.25E-05 |
| Wnt4 | 1.032092 | 8.35E-41 | Igfbp5 | 0.966734 | 2.04E-10 | Pkn3 | 0.300891 | 3.59E-05 |
| Itpr1 | 1.343675 | 1.95E-39 | Pard3b | 0.675212 | 2.35E-10 | Atp13a3 | 0.478454 | 3.67E-05 |
| Heg1 | 1.343672 | 1.50E-38 | Coro1c | 0.510052 | 3.01E-10 | Ralb | 0.297607 | 3.75E-05 |
| Afap1l1 | 0.899891 | 6.04E-38 | Abcb10 | 0.383408 | 3.66E-10 | Tspan5 | 0.42591 | 3.94E-05 |
| Ppp3ca | 0.975857 | 8.33E-36 | Mapk14 | 0.465102 | 4.21E-10 | Asah2 | 0.311963 | 3.98E-05 |
| Prrt4 | 0.612918 | 3.02E-34 | Hn1 | 0.425268 | 4.91E-10 | Obsl1 | 0.385188 | 4.20E-05 |
| Ptprb | 0.981308 | 3.34E-34 | Prkd2 | 0.432691 | 5.00E-10 | Pi16 | 0.290178 | 4.30E-05 |
| Iqgap2 | 1.064367 | 1.14E-33 | Abi3 | 0.435799 | 1.00E-09 | Ext1 | 0.342321 | 4.43E-05 |
| Foxo1 | 1.114616 | 1.57E-33 | Etl4 | 0.550315 | 1.13E-09 | Gldc | 0.339359 | 4.83E-05 |
| Cyp26b1 | 1.195623 | 6.59E-33 | Mcu | 0.527292 | 1.63E-09 | 2510009E07Rik | 0.354645 | 5.85E-05 |
| Fam129a | 0.836598 | 5.92E-31 | Sorbs1 | 0.520273 | 1.76E-09 | Crip2 | 0.32718 | 7.72E-05 |
| Rasgrp3 | 0.804204 | 1.29E-30 | Igf2os | 0.42719 | 1.78E-09 | Klf3 | 0.467021 | 7.83E-05 |
| Lrig1 | 0.875261 | 3.25E-30 | Pik3c2b | 0.369526 | 2.75E-09 | Fbxo34 | 0.459565 | 8.52E-05 |
| Dysf | 0.866786 | 3.61E-30 | Dock3 | 0.434916 | 2.80E-09 | Fgf2 | 0.296869 | 8.68E-05 |
| Nav3 | 1.82983 | 1.65E-28 | Rcsd1 | 0.474643 | 2.90E-09 | Dram2 | 0.302854 | 0.000105 |
| Actn1 | 0.979743 | 2.55E-28 | Hecw1 | 0.388927 | 2.97E-09 | Tbc1d24 | 0.283215 | 0.000132 |
| Arhgef15 | 0.658115 | 5.44E-28 | Ecm1 | 0.378965 | 3.05E-09 | Vim | 0.301306 | 0.000156 |
| Rbm47 | 0.847132 | 7.67E-28 | Spon1 | 0.898879 | 3.28E-09 | Grk5 | 0.39459 | 0.000158 |
| Klf2 | 0.946975 | 9.11E-28 | Daam1 | 0.548368 | 3.38E-09 | Gphn | 0.380816 | 0.000161 |
| Mical2 | 0.669693 | 1.23E-27 | Snx5 | 0.44135 | 3.92E-09 | Slc7a1 | 0.369197 | 0.000163 |
| Lmo7 | 0.766505 | 1.58E-27 | Stim2 | 0.424182 | 5.25E-09 | Dock6 | 0.303079 | 0.000177 |
| Nos3 | 0.801915 | 1.61E-27 | She | 0.355305 | 6.13E-09 | Kcnh1 | 0.318434 | 0.00019 |
| Fgd6 | 0.847455 | 1.83E-27 | Slc7a7 | 0.352117 | 7.18E-09 | Snd1 | 0.296524 | 0.000295 |
| Col13a1 | 0.787717 | 2.08E-27 | Rin3 | 0.464618 | 9.25E-09 | Arf2 | 0.389086 | 0.000307 |
| Plk2 | 0.952031 | 5.74E-27 | Fgd5 | 0.409031 | 1.00E-08 | Itga6 | 0.435009 | 0.000352 |
| Hspg2 | 0.76211 | 1.00E-26 | Nhs | 0.785074 | 1.73E-08 | Pdlim1 | 0.401669 | 0.000419 |
| Pdzd2 | 1.032395 | 7.22E-26 | Ets2 | 0.427769 | 1.75E-08 | Racgap1 | 0.389597 | 0.000457 |
| Slc1a5 | 0.771353 | 1.08E-25 | Elmo1 | 0.473826 | 1.93E-08 | Camta1 | 0.348275 | 0.00047 |
| Tshz3 | 0.808148 | 1.66E-25 | Dock4 | 0.49604 | 2.67E-08 | Lrrc8b | 0.336857 | 0.000553 |
| Dock9 | 0.781906 | 1.83E-24 | Nostrin | 0.316414 | 3.09E-08 | Fam163a | 0.329791 | 0.00057 |
| Plpp1 | 1.044808 | 2.26E-24 | Fabp5 | 0.51439 | 3.12E-08 | Cers6 | 0.432888 | 0.000597 |
| Hecw2 | 0.737608 | 3.21E-24 | Radil | 0.285478 | 3.81E-08 | Ephb4 | 0.300947 | 0.000622 |
| Zfp521 | 0.733362 | 7.64E-24 | Tjp2 | 0.493347 | 4.04E-08 | Tsc22d3 | 0.261492 | 0.000642 |
| Ece1 | 0.757298 | 9.22E-24 | Adam19 | 0.661017 | 4.15E-08 | Arrb1 | 0.293797 | 0.000692 |
| Meis2 | 0.786048 | 7.62E-22 | Adgrf5 | 0.373532 | 4.55E-08 | Golph3 | 0.332611 | 0.000714 |
| Flt1 | 0.644233 | 8.94E-22 | Capn2 | 0.446664 | 4.83E-08 | Fbxw7 | 0.356522 | 0.000892 |
| Col5a1 | 0.73524 | 2.00E-21 | Ccm2l | 0.275951 | 4.87E-08 | Dock5 | 0.344517 | 0.000909 |
| Tspan9 | 0.626691 | 3.42E-21 | Ano6 | 0.423906 | 5.99E-08 | Plcd3 | 0.266931 | 0.000966 |
| Cbfa2t3 | 0.878752 | 3.94E-21 | Nrp2 | 0.393279 | 6.27E-08 | Kras | 0.350531 | 0.001026 |
| Ets1 | 0.742568 | 2.79E-20 | 2900026A02Rik | 0.420295 | 6.78E-08 | Tns1 | 0.380192 | 0.001069 |
| Tmcc2 | 0.536637 | 4.26E-20 | Rap1b | 0.444249 | 7.40E-08 | Mdm1 | 0.311385 | 0.001083 |
| Ubash3b | 0.82033 | 1.57E-19 | Ppp1r2 | 0.494282 | 7.65E-08 | Plxnd1 | 0.362067 | 0.001204 |
| Elavl4 | 0.884636 | 1.65E-19 | Polk | 0.346316 | 7.91E-08 | Calm2 | 0.370312 | 0.001295 |
| Ttc9 | 0.415966 | 1.73E-19 | Stk39 | 0.47975 | 8.45E-08 | Bace2 | 0.470303 | 0.001315 |
| Shroom4 | 0.894583 | 6.68E-19 | Anxa2 | 0.537319 | 1.09E-07 | Utrn | 0.284025 | 0.001386 |
| Arhgef3 | 1.03408 | 7.51E-19 | Rem1 | 0.251334 | 1.43E-07 | F2r | 0.305 | 0.001399 |
| Mapkapk3 | 0.677288 | 7.56E-19 | Tead4 | 0.385107 | 1.44E-07 | Fbn1 | 0.347137 | 0.001457 |
| Akap13 | 0.557696 | 8.14E-19 | Zc3hav1 | 0.337718 | 2.01E-07 | Elk3 | 0.299956 | 0.001637 |
| Sptbn1 | 0.589384 | 9.71E-19 | Ifitm10 | 0.2768 | 2.24E-07 | B4galt5 | 0.326809 | 0.001664 |
| Tgfbr2 | 0.7078 | 1.06E-18 | Iqsec1 | 0.494979 | 2.38E-07 | AU020206 | 0.261518 | 0.001721 |
| Mtss1 | 0.718765 | 1.16E-18 | Adam23 | 0.512111 | 2.86E-07 | Ikzf3 | 0.327869 | 0.001761 |
| Dapk1 | 0.751303 | 2.57E-18 | Klhl3 | 0.351008 | 3.28E-07 | Ghr | 0.562392 | 0.001911 |

| | | | | | | | | |
|---|---|---|---|---|---|---|---|---|
| Galnt18 | 1.176309 | 5.26E-18 | Mirg | 0.527047 | 3.86E-07 | Pdgfd | 0.441834 | 0.002064 |
| Podxl | 0.634993 | 2.03E-17 | Dmgdh | 0.275389 | 3.98E-07 | Tmem164 | 0.453664 | 0.002227 |
| Palm | 0.601418 | 3.11E-17 | Macf1 | 0.306755 | 4.86E-07 | Sema3a | 0.655732 | 0.00288 |
| Vav3 | 0.739164 | 6.57E-17 | Gap43 | 0.639616 | 5.81E-07 | Pcsk5 | 0.348679 | 0.00288 |
| Colgalt2 | 0.643449 | 7.21E-17 | Tmem131 | 0.381563 | 6.47E-07 | 9430020K01Rik | 0.319137 | 0.003621 |
| Nrg1 | 1.598708 | 8.25E-17 | Plpp3 | 0.735919 | 6.68E-07 | Esyt2 | 0.296499 | 0.00367 |
| Meg3 | 0.860245 | 1.25E-16 | Rassf8 | 0.591721 | 7.21E-07 | Col4a3bp | 0.312725 | 0.00416 |
| Vstm5 | 0.366864 | 1.51E-16 | Prkca | 0.553554 | 7.23E-07 | Pcgf5 | 0.403377 | 0.005375 |
| Kank1 | 0.66051 | 1.98E-16 | Cpne8 | 0.319577 | 7.42E-07 | Fndc3b | 0.261856 | 0.005584 |
| Smtnl2 | 0.480814 | 2.65E-16 | Kdr | 0.43344 | 7.79E-07 | Elf4 | 0.313906 | 0.006123 |
| Fam155a | 1.767978 | 3.04E-16 | Adcy4 | 0.4009 | 8.29E-07 | Apcdd1 | 0.405603 | 0.006262 |
| Uaca | 0.654491 | 4.44E-16 | Rgs3 | 0.406437 | 8.67E-07 | Taok3 | 0.3223 | 0.006346 |
| Plxnc1 | 0.556841 | 4.85E-16 | Tie1 | 0.279632 | 1.24E-06 | Adgra2 | 0.289451 | 0.006356 |
| Cd9 | 0.621394 | 7.83E-16 | Tanc1 | 0.334433 | 1.29E-06 | Tspan12 | 0.476965 | 0.006864 |
| Entpd1 | 0.74812 | 8.41E-16 | Acot11 | 0.290148 | 1.42E-06 | Whsc1 | 0.253738 | 0.006992 |
| Nab1 | 0.639612 | 1.25E-15 | Adamts19 | 0.631943 | 1.46E-06 | Chchd3 | 0.288836 | 0.007314 |
| Kank3 | 0.419935 | 2.66E-15 | Fhl2 | 0.414917 | 1.64E-06 | Lnx2 | 0.256274 | 0.008034 |
| Fam222b | 0.54542 | 5.18E-15 | Hspa12b | 0.340353 | 1.96E-06 | Thsd1 | 0.255724 | 0.008181 |
| Agap1 | 0.645957 | 6.12E-15 | Itga1 | 0.478584 | 1.97E-06 | Exoc3l2 | 0.291575 | 0.009015 |
| Jam2 | 0.429932 | 8.30E-15 | Cd93 | 0.32958 | 2.05E-06 | Inf2 | 0.278845 | 0.009271 |
| Spag9 | 0.519363 | 9.74E-15 | Gm42418 | 0.36975 | 2.13E-06 | Vezt | 0.319287 | 0.010444 |
| Scml4 | 0.289735 | 1.13E-14 | Ebf1 | 0.790448 | 2.88E-06 | Klf10 | 0.331721 | 0.01179 |
| Neat1 | 0.667106 | 1.22E-14 | Aff1 | 0.45855 | 3.13E-06 | Pik3r1 | 0.343021 | 0.012045 |
| Apba1 | 0.792856 | 1.41E-14 | Pde4d | 0.602952 | 3.16E-06 | Ggta1 | 0.330725 | 0.01217 |
| Spats2l | 0.753949 | 2.09E-14 | Rapgef1 | 0.39682 | 3.28E-06 | Ksr1 | 0.384215 | 0.012632 |
| Fam167a | 0.326269 | 3.00E-14 | Plcd1 | 0.355144 | 3.35E-06 | Dpysl2 | 0.311841 | 0.016995 |
| Lhfp | 0.68275 | 3.44E-14 | Rai14 | 0.436153 | 3.41E-06 | Gm20532 | 0.288566 | 0.018603 |
| Slc8a3 | 0.574925 | 4.30E-14 | Itpkb | 0.47169 | 3.69E-06 | Rhoa | 0.254693 | 0.020015 |
| Ppp1r16b | 0.573874 | 1.45E-13 | B4galt1 | 0.449429 | 4.12E-06 | Itsn1 | 0.279619 | 0.021553 |
| Slc9a3r2 | 0.385506 | 1.98E-13 | Tmem2 | 0.550905 | 4.64E-06 | Lrrc8c | 0.323795 | 0.022455 |
| Npr3 | 0.881119 | 2.16E-13 | Slit2 | 0.674101 | 5.06E-06 | Tbl1xr1 | 0.260348 | 0.022541 |
| Capn5 | 0.511436 | 8.69E-13 | Slc43a3 | 0.364684 | 5.12E-06 | Fam78b | 0.399581 | 0.025686 |
| Amotl1 | 0.467397 | 1.33E-12 | Ppp1r13b | 0.371075 | 5.14E-06 | Ackr3 | 0.263772 | 0.029548 |
| Sulf1 | 1.050423 | 1.93E-12 | Syndig1 | 0.52607 | 5.48E-06 | Myo1c | 0.274341 | 0.031411 |
| Piezo1 | 0.548978 | 1.97E-12 | Ddah1 | 0.451385 | 5.81E-06 | Dok4 | 0.314081 | 0.034501 |
| Tnfaip2 | 0.47904 | 2.80E-12 | Btbd9 | 0.385706 | 6.26E-06 | Npc2 | 0.286846 | 0.03601 |
| Fxyd5 | 0.420623 | 3.42E-12 | Ccdc6 | 0.396387 | 6.65E-06 | Ndrg3 | 0.254735 | 0.036069 |
| Notch1 | 0.475358 | 3.96E-12 | Arhgef1 | 0.319478 | 9.42E-06 | Cped1 | 0.587021 | 0.036445 |
| Arfgef1 | 0.553056 | 4.25E-12 | Fn1 | 0.453687 | 1.13E-05 | Arap3 | 0.274376 | 0.037725 |
| Adam15 | 0.395345 | 4.95E-12 | Ptprf | 0.358809 | 1.20E-05 | Rnf220 | 0.638189 | 0.037895 |
| Foxp1 | 0.50308 | 5.28E-12 | Palmd | 0.27265 | 1.24E-05 | Sgcd | 0.359889 | 0.038969 |
| Arhgap31 | 0.477418 | 5.66E-12 | Eng | 0.360601 | 1.25E-05 | Adgrl4 | 0.337506 | 0.045926 |
| Zdhhc14 | 0.556968 | 8.26E-12 | Rasa3 | 0.423998 | 1.49E-05 | Limch1 | 0.278861 | 0.047465 |
| Nuak2 | 0.362559 | 1.42E-11 | Parvb | 0.363494 | 1.52E-05 | | | |

| Table S2C | Mid EndoMT | | | | | | | |
|---|---|---|---|---|---|---|---|---|
| Gene Name | avg_logFC | p_val_adj | Gene Name | avg_logFC | p_val_adj | Gene Name | avg_logFC | p_val_adj |
| Trpm3 | 1.349987 | 7.79E-17 | Nr6a1os | 0.439539 | 0.000174 | Add2 | 0.33684 | 0.006023 |
| Frmd4b | 0.598034 | 1.34E-12 | St3gal5 | 0.421569 | 0.000317 | Ctnnd2 | 0.454814 | 0.0067 |
| St6gal1 | 0.751305 | 2.14E-12 | Gm15563 | 0.396543 | 0.000319 | Itpr2 | 0.51502 | 0.007175 |
| Thsd7a | 0.663261 | 2.39E-10 | Ppp1r9a | 0.480924 | 0.00035 | Efna5 | 0.369035 | 0.007311 |
| Plagl1 | 0.643032 | 3.06E-10 | Nrp1 | 0.336754 | 0.000372 | Snhg12 | 0.395882 | 0.007513 |
| Sorcs1 | 0.474017 | 1.09E-09 | Gm26733 | 0.486079 | 0.000414 | Jazf1 | 0.495432 | 0.008036 |
| Foxp2 | 0.54797 | 5.49E-09 | Lrrc58 | 0.50295 | 0.000527 | Adamts6 | 0.45449 | 0.010874 |
| Edn1 | 0.493509 | 1.38E-08 | Inpp4b | 0.431351 | 0.000561 | Stxbp6 | 0.423744 | 0.012066 |
| Bmper | 0.706467 | 7.57E-08 | Grip1 | 0.335129 | 0.000863 | Sncaip | 0.518625 | 0.012684 |
| St8sia1 | 0.470719 | 3.24E-07 | Lrrc4 | 0.457914 | 0.001044 | Adam12 | 0.381556 | 0.013314 |
| Ano1 | 1.062594 | 3.76E-07 | Gm26936 | 0.397288 | 0.001281 | Taf1d | 0.378527 | 0.014321 |
| Prickle2 | 0.536743 | 5.48E-07 | Igf2r | 0.35982 | 0.001729 | Trpc6 | 0.367476 | 0.014782 |
| Ptprm | 0.482324 | 7.69E-07 | Col23a1 | 0.551661 | 0.001961 | Rasl11b | 0.289372 | 0.02398 |
| Samd5 | 0.647571 | 1.33E-06 | Klhl4 | 0.435778 | 0.002337 | Frmd4a | 0.344305 | 0.026558 |
| S1pr1 | 0.480029 | 1.80E-06 | Aff4 | 0.295571 | 0.002971 | Cacna2d1 | 0.536355 | 0.028663 |
| Zfpm2 | 0.541495 | 5.24E-06 | Bcl2 | 0.44039 | 0.003248 | Emcn | 0.310931 | 0.032048 |
| Acer2 | 0.609833 | 5.94E-06 | Prex2 | 0.3575 | 0.003382 | Gm26883 | 0.417785 | 0.042202 |
| Ahr | 0.52502 | 8.74E-06 | Gm16599 | 0.424089 | 0.003531 | Dennd5b | 0.289722 | 0.044398 |
| Mecom | 0.418562 | 4.59E-05 | Fat3 | 0.360998 | 0.004624 | Pappa | 0.335914 | 0.048184 |
| Mast4 | 0.40726 | 0.000123 | Prtg | 0.492713 | 0.005441 | Pcdh7 | 0.533306 | 0.049117 |

| Table S2D | Late EndoMT | | | | | | | |
|---|---|---|---|---|---|---|---|---|
| Gene Name | avg_logFC | p_val_adj | Gene Name | avg_logFC | p_val_adj | Gene Name | avg_logFC | p_val_adj |
| 2610035D17Rik | 2.833626 | 1.35E-102 | Ank | 0.801023 | 1.19E-12 | Gata6 | 0.50008 | 0.000145 |
| Garnl3 | 2.1249 | 9.93E-96 | Inpp4a | 0.550713 | 1.45E-12 | Klhl13 | 0.396987 | 0.000164 |
| Sox5 | 2.193098 | 9.38E-87 | Rai1 | 0.52407 | 1.72E-12 | Tmem64 | 0.396855 | 0.000168 |
| Robo2 | 3.366691 | 1.21E-75 | Plscr1 | 0.440918 | 3.87E-12 | Vangl1 | 0.511256 | 0.000237 |
| Pdgfra | 1.800816 | 2.61E-70 | Eya4 | 0.807625 | 4.38E-12 | Nos1ap | 0.803628 | 0.000254 |
| Vcan | 1.708266 | 1.33E-60 | Tnc | 0.876627 | 9.42E-12 | Fat1 | 0.399365 | 0.000277 |
| Lmo4 | 1.389055 | 2.38E-49 | Ppfibp2 | 0.360533 | 2.41E-11 | Gm16070 | 0.257371 | 0.000278 |
| Sipa1l1 | 1.262276 | 3.99E-48 | Notch2 | 0.66432 | 3.03E-11 | Gli3 | 0.363792 | 0.000324 |
| Itga4 | 1.105348 | 7.05E-48 | Pcolce | 0.252031 | 3.13E-11 | Dnah7b | 0.27778 | 0.000326 |
| Pde7b | 1.725787 | 6.85E-47 | St6galnac51 | 0.59461 | 3.46E-11 | Msi2 | 0.405239 | 0.000332 |
| Plce1 | 1.331234 | 1.67E-43 | 5033428I22Rik | 0.595801 | 3.85E-11 | Has21 | 0.472046 | 0.000411 |
| Ncam1 | 1.45176 | 3.79E-42 | Papss2 | 1.058369 | 6.63E-11 | Adk | 0.409007 | 0.000413 |
| Itgb5 | 1.185144 | 3.24E-40 | Pdlim3 | 0.591247 | 2.12E-10 | Epb41l4a | 0.381772 | 0.000421 |
| Specc1 | 1.149487 | 1.99E-39 | Lpar1 | 0.636778 | 3.43E-10 | St3gal51 | 0.429606 | 0.000426 |
| Erbb3 | 0.720258 | 3.21E-39 | Rgma | 0.340648 | 3.45E-10 | Zswim6 | 0.49606 | 0.000484 |
| Aff3 | 1.73266 | 1.20E-38 | Arhgef40 | 0.543958 | 4.46E-10 | Fzd2 | 0.292552 | 0.000488 |
| Sox91 | 1.231544 | 5.38E-37 | Pik3cb | 0.646671 | 5.10E-10 | Trim2 | 0.293123 | 0.000495 |
| Gpc6 | 1.408888 | 5.84E-36 | Npvf | 0.312714 | 5.21E-10 | Prkdc | 0.469491 | 0.000503 |
| Twist1 | 0.89786 | 2.08E-35 | Slc25a21 | 0.739324 | 5.24E-10 | Ccnd2 | 0.473719 | 0.000514 |
| Chst11 | 1.468163 | 2.53E-35 | Etv1 | 0.610657 | 5.46E-10 | Fryl | 0.460545 | 0.000516 |
| Clmp | 0.91519 | 1.03E-34 | Mark1 | 0.648846 | 5.91E-10 | Foxp21 | 0.63091 | 0.000529 |
| Irs1 | 1.158876 | 2.79E-34 | Ptpn13 | 0.454214 | 7.63E-10 | Astn2 | 0.301128 | 0.000593 |
| Gpc3 | 2.036473 | 3.46E-34 | Ror1 | 0.842734 | 1.19E-09 | Tnfaip8 | 0.50761 | 0.00065 |
| Cdon | 1.426201 | 2.80E-33 | Zcchc11 | 0.464301 | 1.37E-09 | Zfp516 | 0.402285 | 0.000678 |
| Pknox2 | 1.352748 | 4.36E-33 | Ssbp2 | 0.675107 | 1.42E-09 | Ccny | 0.391894 | 0.000693 |
| Greb1 | 0.954237 | 1.29E-32 | Wdpcp | 0.575406 | 2.08E-09 | Gata3 | 0.462439 | 0.000716 |
| Dnm3os | 0.791632 | 1.70E-32 | Slc24a2 | 1.007482 | 2.45E-09 | Cpeb2 | 0.462668 | 0.000784 |
| Hs3st3a1 | 0.966656 | 2.84E-31 | Tbx2 | 0.365265 | 5.24E-09 | Plac1 | 1.022152 | 0.000913 |
| Col9a3 | 0.491017 | 3.25E-31 | Gm11149 | 0.530953 | 6.20E-09 | Oxct1 | 0.421595 | 0.000936 |
| Epha7 | 1.471453 | 4.01E-31 | Ankrd44 | 0.71874 | 7.59E-09 | Sh3pxd2a | 0.45613 | 0.00103 |
| Ltbp1 | 0.995384 | 2.21E-30 | Gulp1 | 0.507429 | 8.34E-09 | Gm14582 | 0.553526 | 0.00107 |
| Rap1gap2 | 0.846318 | 3.34E-30 | Me1 | 0.478643 | 8.43E-09 | Spint2 | 0.282599 | 0.001072 |
| Pde1a | 0.991887 | 5.23E-30 | Haus7 | 0.309421 | 8.98E-09 | Slc4a7 | 0.425889 | 0.00108 |
| Epha3 | 1.922081 | 1.72E-29 | Grid1 | 0.719773 | 1.47E-08 | Sesn3 | 0.688217 | 0.001207 |
| Dlc1 | 0.93553 | 1.91E-29 | Ror2 | 0.613741 | 1.78E-08 | Map4 | 0.380973 | 0.001278 |
| Unc5c | 1.459003 | 3.41E-29 | Chsy1 | 0.584657 | 2.19E-08 | Susd1 | 0.315542 | 0.001352 |
| Epb41l1 | 0.59454 | 8.56E-29 | Pde3a | 0.699801 | 2.54E-08 | Bmper1 | 0.39649 | 0.001543 |
| Sox61 | 0.968305 | 6.94E-28 | Grik2 | 0.727407 | 3.47E-08 | Csgalnact1 | 0.618002 | 0.001659 |
| Erbb4 | 1.359651 | 1.72E-27 | Enah | 0.586956 | 5.92E-08 | Med141 | 0.501719 | 0.001792 |
| Celf4 | 0.894739 | 4.97E-27 | Kitl | 0.679225 | 8.25E-08 | Frs2 | 0.389172 | 0.001827 |
| St3gal4 | 1.09368 | 7.14E-26 | Lrp1 | 0.382155 | 1.93E-07 | Kcnq5 | 0.450155 | 0.001827 |
| Zbtb20 | 0.796366 | 8.95E-26 | Gm21807 | 0.367708 | 2.22E-07 | Znrf3 | 0.408047 | 0.002168 |
| Serpine2 | 0.892532 | 1.13E-25 | Nav2 | 0.887957 | 3.04E-07 | 1010001N08Rik | 0.264251 | 0.002374 |
| Slc1a3 | 1.097389 | 6.25E-25 | Ctnna3 | 0.654036 | 4.36E-07 | Mpp6 | 0.385584 | 0.002575 |
| Gcnt1 | 0.830739 | 7.51E-25 | Dact3 | 0.29733 | 5.65E-07 | Myh10 | 0.355223 | 0.002634 |
| Ociad2 | 0.460667 | 2.30E-24 | Tiam2 | 0.637301 | 6.08E-07 | Wwp2 | 0.438261 | 0.002715 |
| Ntng1 | 0.77805 | 5.23E-24 | Sntb1 | 0.445704 | 8.15E-07 | Lrrc16a | 0.309019 | 0.002822 |
| Postn | 0.69334 | 1.48E-23 | Akr1b8 | 0.285109 | 8.77E-07 | Zfp618 | 0.395725 | 0.002965 |
| Mcc | 0.9578 | 5.84E-23 | Tenm4 | 0.330381 | 1.04E-06 | Sox5os4 | 0.26225 | 0.003051 |
| Prrx2 | 0.829954 | 1.32E-22 | Farp1 | 0.385755 | 1.20E-06 | Lef1 | 0.57703 | 0.003538 |
| Psd3 | 1.204449 | 1.61E-22 | Hdac4 | 0.51643 | 1.67E-06 | Gm42439 | 0.52919 | 0.003745 |
| Clstn2 | 1.324245 | 4.13E-22 | Crispld1 | 0.423329 | 1.83E-06 | St6galnac6 | 0.292316 | 0.004659 |
| Esrrg | 1.640133 | 1.24E-21 | Cdh11 | 0.484152 | 1.94E-06 | Cdk14 | 0.39418 | 0.004867 |
| Cttnbp2 | 0.91941 | 1.67E-20 | Twist2 | 0.533857 | 2.05E-06 | Tmcc3 | 0.546638 | 0.006149 |
| Col9a1 | 0.506117 | 1.84E-20 | Cux2 | 0.483185 | 2.21E-06 | Hsd17b7 | 0.260769 | 0.006221 |
| Fbn2 | 0.765847 | 2.13E-20 | Grb10 | 0.389169 | 2.60E-06 | Tcf7l2 | 0.346355 | 0.006367 |
| Ppfibp1 | 0.72009 | 7.66E-20 | Hs3st3b1 | 0.559289 | 3.35E-06 | Rasl11b1 | 0.34905 | 0.006452 |
| Pag1 | 0.746713 | 1.53E-19 | P3h2 | 0.707494 | 3.63E-06 | Fam171a1 | 0.34447 | 0.006527 |
| Tenm3 | 0.813528 | 2.15E-19 | Smad9 | 0.273659 | 4.20E-06 | Zfp644 | 0.301797 | 0.008271 |
| Kcnt2 | 0.83264 | 2.76E-19 | Prkce | 0.400853 | 4.38E-06 | Rnd2 | 0.365659 | 0.008464 |
| Dock7 | 0.593315 | 1.65E-17 | Cbfa2t2 | 0.494658 | 7.38E-06 | Tspan8 | 0.288647 | 0.008787 |

| | | | | | | | | |
|---|---|---|---|---|---|---|---|---|
| Antxr1 | 0.778996 | 2.37E-17 | Xpo1 | 0.371978 | 9.07E-06 | Igf1 | 0.499798 | 0.009871 |
| Id2 | 0.845618 | 2.85E-17 | Fat4 | 0.597126 | 1.16E-05 | Foxo3 | 0.335005 | 0.01014 |
| A630012P03Rik | 0.745574 | 4.94E-17 | Dyrk1a | 0.419869 | 1.34E-05 | Igf1r | 0.305188 | 0.011364 |
| Snai1 | 0.444565 | 6.49E-17 | Msx1 | 0.506359 | 1.56E-05 | Mme | 0.439861 | 0.01139 |
| Cthrc1 | 0.752383 | 7.86E-17 | Sdk2 | 0.265251 | 1.85E-05 | Galntl61 | 0.705277 | 0.011573 |
| Tgfbi | 0.602275 | 9.76E-17 | Prdm6 | 0.658388 | 2.21E-05 | Apoe | 0.474274 | 0.011712 |
| Auts2 | 0.694468 | 1.53E-16 | Hipk2 | 0.419166 | 2.30E-05 | Papss1 | 0.328294 | 0.013374 |
| Epb41l3 | 0.877174 | 2.49E-16 | Osbpl1a | 0.449389 | 2.76E-05 | Pde1c1 | 0.421638 | 0.01383 |
| Ptprd1 | 0.773595 | 4.03E-16 | Synpo2 | 0.692928 | 2.90E-05 | Man2a2 | 0.273273 | 0.014319 |
| Trpv4 | 0.299244 | 8.22E-16 | Slc4a4 | 0.390567 | 2.98E-05 | Jarid2 | 0.336907 | 0.01579 |
| Diras2 | 0.387436 | 9.27E-16 | Hmga2 | 0.509355 | 3.18E-05 | Slc25a36 | 0.284013 | 0.019115 |
| Bach2 | 0.69711 | 1.79E-15 | Fhit | 0.495831 | 3.36E-05 | Cdh10 | 0.562272 | 0.020237 |
| Setbp1 | 0.99 | 8.95E-15 | Rtn1 | 0.437602 | 4.38E-05 | E130006D01Rik | 0.254012 | 0.022992 |
| Klhl29 | 0.68105 | 1.65E-14 | Arhgap5 | 0.437685 | 4.75E-05 | Ddhd2 | 0.328518 | 0.027026 |
| Cdh19 | 0.253183 | 2.02E-14 | Flrt3 | 0.260091 | 4.77E-05 | Ndfip1 | 0.269452 | 0.03206 |
| Tbx3 | 0.446472 | 2.28E-14 | Timp31 | 0.741726 | 6.47E-05 | Msi1 | 0.343235 | 0.033086 |
| Tbx20 | 0.502674 | 2.34E-14 | Robo1 | 0.571611 | 8.13E-05 | Pgm2 | 0.492026 | 0.036258 |
| Fbln2 | 0.617375 | 3.67E-14 | Clcn5 | 0.505118 | 8.70E-05 | Plekha6 | 0.327433 | 0.038912 |
| Dpysl3 | 0.78731 | 5.60E-14 | Tox | 0.356083 | 9.27E-05 | Cpq | 0.347992 | 0.039293 |
| Ugdh | 0.67774 | 9.30E-14 | Gyg | 0.393557 | 0.000115 | Frmd4a1 | 0.277999 | 0.04375 |
| Ddr1 | 0.378689 | 9.37E-14 | Tuba1c | 0.476651 | 0.000118 | Mb21d2 | 0.342261 | 0.046038 |
| A930011G23Rik | 0.933692 | 3.34E-13 | Apbb2 | 0.427997 | 0.000129 | Fign | 0.441498 | 0.046505 |
| Sh2d4a | 0.372414 | 5.49E-13 | Rnf130 | 0.407487 | 0.000144 | Mex3b | 0.291773 | 0.047159 |
| Epha4 | 0.829233 | 7.67E-13 | | | | | | |
| Gm42906 | 0.677375 | 9.91E-13 | | | | | | |

| **Table S3** | $Ift20^{+/+}$ vs $Ift20^{-/-}$ | | $IIft20^{+/-}$ vs $Ift20^{-/-}$ | | | $Ift20^{+/+}$ vs $Ift20^{-/-}$ | | $IIft20^{+/-}$ vs $Ift20^{-/-}$ | |
|---|---|---|---|---|---|---|---|---|---|
| Gene Name | avg_logFC | p_val_adj | avg_logFC | p_val_adj | Gene Name | avg_logFC | p_val_adj | avg_logFC | p_val_adj |
| 2610035D17Rik | 1.73422 | 1.19E-19 | 1.336436 | 1.15E-12 | Myh6 | 0.598603 | 4.36E-12 | 0.704152 | 1.42E-16 |
| 9530026P05Rik | -0.41273 | 0.010654 | -0.40976 | 0.010385 | Myo6 | -0.66297 | 2.10E-10 | -0.67421 | 3.35E-11 |
| Abhd17a | -0.4808 | 0.002947 | -0.44125 | 0.014432 | Nav3 | -0.53962 | 1.25E-05 | -0.54607 | 0.014619 |
| Abhd18 | -0.41474 | 0.000171 | -0.54647 | 5.15E-11 | Ncam1 | 0.784758 | 4.10E-07 | 0.838262 | 2.66E-11 |
| Acer2 | -0.55875 | 1.40E-06 | -0.70007 | 3.52E-10 | Nfia | 0.400701 | 0.000235 | 0.329679 | 0.005872 |
| Adgrf5 | -0.47933 | 5.78E-05 | -0.51363 | 2.90E-06 | Notch4 | -0.41625 | 6.98E-05 | -0.44878 | 4.91E-05 |
| Adgrl4 | -0.99108 | 2.04E-12 | -0.78346 | 1.23E-06 | Nrp1 | -0.42604 | 6.47E-05 | -0.54609 | 1.84E-07 |
| Aff4 | -0.50891 | 1.59E-13 | -0.41749 | 1.37E-08 | P4ha1 | -0.57241 | 9.25E-09 | -0.71687 | 1.31E-14 |
| Arglu1 | -0.6072 | 3.11E-14 | -0.5413 | 3.53E-11 | Pcdh7 | -0.67194 | 0.000453 | -0.67427 | 2.82E-05 |
| Arhgap29 | -0.76583 | 1.40E-10 | -0.85403 | 9.51E-15 | Pde1a | 0.478376 | 1.39E-05 | 0.387543 | 0.000526 |
| Arhgef28 | -0.90829 | 3.02E-05 | -0.90085 | 5.30E-05 | Pde4b | -0.91852 | 3.46E-18 | -1.01171 | 6.67E-24 |
| Bcar3 | -0.79935 | 7.57E-11 | -0.78429 | 4.75E-11 | Pde8a | -0.70104 | 1.19E-05 | -0.74682 | 1.67E-07 |
| Bnip3 | -0.6728 | 3.01E-24 | -0.64401 | 1.55E-21 | Pdgfra | 0.713645 | 1.01E-07 | 0.818758 | 5.88E-09 |
| Cacna1d | 0.41885 | 0.00225 | 0.276023 | 0.036115 | Pdk1 | -0.82805 | 7.54E-24 | -0.85868 | 1.83E-28 |
| Cav1 | -0.50609 | 0.000502 | -0.50164 | 0.000247 | Pdlim3 | 0.833943 | 5.82E-16 | 0.969886 | 4.38E-23 |
| Ccdc141 | 0.493672 | 4.02E-08 | 0.651564 | 7.54E-13 | Peak1 | -0.5584 | 2.02E-09 | -0.53725 | 7.29E-09 |
| Ccdc58 | -0.543 | 1.10E-07 | -0.38971 | 0.024703 | Peg3 | -0.56039 | 1.26E-06 | -0.52393 | 1.63E-06 |
| Cd109 | -0.51917 | 0.000119 | -0.41985 | 0.001406 | Pfkfb3 | -0.59633 | 1.94E-07 | -0.72042 | 3.42E-15 |
| Cdh5 | -0.44423 | 1.11E-05 | -0.34952 | 0.006534 | Pgk1 | -1.02365 | 1.30E-31 | -0.90223 | 7.65E-23 |
| Cdkn1c | 0.405894 | 0.028819 | 0.560989 | 2.40E-07 | Piezo2 | -0.91348 | 1.74E-06 | -0.88568 | 2.68E-05 |
| Cdon | 0.806307 | 1.16E-08 | 0.671646 | 2.59E-05 | Pkm | -0.59219 | 1.61E-14 | -0.43689 | 1.68E-06 |
| Celf4 | 0.597692 | 2.18E-15 | 0.452593 | 3.12E-09 | Plagl1 | -0.40348 | 0.007438 | -0.60937 | 6.44E-07 |
| Chsy1 | 0.55069 | 3.93E-12 | 0.476944 | 5.81E-07 | Plce1 | 0.801824 | 2.99E-11 | 0.786456 | 4.50E-15 |
| Clcn3 | -0.42143 | 0.016478 | -0.51114 | 0.000771 | Plcl2 | 0.414774 | 7.82E-05 | 0.35406 | 0.01244 |
| Clvs1 | -0.86797 | 1.09E-14 | -0.78501 | 4.13E-11 | Plod2 | -0.75908 | 5.14E-14 | -0.68446 | 9.66E-12 |
| Clybl | -0.66209 | 4.93E-11 | -0.61838 | 1.74E-08 | Ppp2r3a | -0.71799 | 1.03E-06 | -0.68547 | 1.44E-05 |
| Col13a1 | 0.455505 | 4.77E-06 | 0.294049 | 0.003335 | Prelid2 | -0.6294 | 5.56E-13 | -0.69081 | 1.29E-15 |
| Col23a1 | -0.87771 | 1.14E-05 | -0.88767 | 9.71E-08 | Prex2 | -0.56694 | 2.38E-08 | -0.6318 | 1.58E-09 |
| Col3a1 | 0.537461 | 1.98E-06 | 0.6112 | 8.09E-10 | Prkca | 0.408533 | 0.000202 | 0.42083 | 1.40E-06 |
| Col4a2 | -0.41432 | 1.04E-05 | -0.68063 | 1.33E-14 | Prkce | 0.704361 | 1.16E-16 | 0.44516 | 7.91E-07 |
| Col5a1 | 0.433113 | 0.002832 | 0.295409 | 0.005247 | Prpf4b | -0.50548 | 6.09E-11 | -0.3133 | 0.001877 |
| Creb5 | 0.48342 | 2.25E-05 | 0.416038 | 0.000396 | Prrx2 | 0.500816 | 1.05E-07 | 0.445944 | 1.22E-07 |
| Ctnna3 | 0.974145 | 5.92E-17 | 0.760678 | 4.86E-12 | Psd3 | 0.563992 | 0.005674 | 0.796163 | 4.22E-10 |
| Cttnbp2 | 0.687814 | 2.83E-09 | 0.628311 | 4.63E-07 | Ptprd | 0.56069 | 0.00114 | 0.514009 | 0.000647 |
| Ddx26b | -0.82082 | 5.11E-09 | -0.79949 | 4.29E-08 | Ptpre | -0.43335 | 0.008575 | -0.4346 | 0.003805 |
| Dennd5b | -0.53334 | 6.34E-10 | -0.52025 | 7.53E-09 | Ralgps2 | -0.60845 | 8.21E-07 | -0.70845 | 1.99E-09 |
| Dock7 | 0.400159 | 7.78E-06 | 0.343493 | 0.000868 | Rapgef5 | -1.34581 | 1.71E-37 | -1.30051 | 4.66E-35 |
| Dpp6 | -0.45379 | 8.29E-05 | -0.45885 | 3.39E-05 | Rasal2 | -0.41898 | 0.001853 | -0.43975 | 0.000741 |
| Dpysl3 | 0.577011 | 1.18E-09 | 0.348367 | 0.001514 | Rasgrp3 | -0.41979 | 0.044516 | -0.46119 | 0.004282 |
| Egfl7 | -0.79785 | 9.17E-22 | -0.58507 | 1.69E-12 | Rbms3 | 0.561833 | 1.71E-08 | 0.412766 | 4.07E-05 |
| Egln3 | -0.74612 | 4.00E-24 | -0.81842 | 3.12E-30 | Rgs7bp | 0.485706 | 0.002924 | 0.308192 | 0.037041 |
| Emcn | -1.08045 | 2.17E-10 | -1.10649 | 1.22E-10 | Robo2 | 1.791456 | 5.14E-11 | 2.037856 | 4.11E-12 |
| Eno1 | -0.84936 | 4.67E-26 | -0.39744 | 2.00E-05 | Rplp1 | 0.418163 | 1.58E-08 | 0.504465 | 4.73E-10 |
| Epha3 | 0.734162 | 0.006603 | 0.977472 | 0.001616 | S1pr1 | -0.63714 | 7.96E-07 | -0.54859 | 2.27E-05 |
| Epha4 | 0.805509 | 1.13E-12 | 0.699564 | 4.01E-06 | Sept11 | -0.84005 | 3.40E-21 | -0.80607 | 1.60E-18 |
| Epha7 | 1.177162 | 5.68E-12 | 1.179521 | 1.91E-12 | Shroom2 | -0.96431 | 1.32E-18 | -0.9328 | 2.19E-17 |
| Erbb4 | 1.030288 | 1.42E-13 | 0.867711 | 2.39E-09 | Slc16a3 | -0.45274 | 2.81E-05 | -0.47771 | 8.66E-06 |
| Erdr1 | 0.767856 | 5.81E-23 | 0.39988 | 3.09E-05 | Slc22a23 | -1.04711 | 1.37E-15 | -1.16345 | 1.33E-17 |
| Exoc3l4 | -0.66346 | 1.15E-09 | -0.51001 | 9.10E-05 | Slc2a1 | -1.09147 | 1.37E-26 | -1.05687 | 1.69E-23 |
| Exoc6 | -0.71456 | 1.73E-11 | -0.87795 | 9.81E-16 | Slc2a3 | -0.88131 | 9.29E-19 | -0.92648 | 6.18E-19 |
| Eya4 | 0.609518 | 5.41E-10 | 0.480544 | 6.14E-06 | Slc8a1 | 1.07403 | 2.79E-12 | 0.622299 | 0.000229 |
| Fam162a | -0.69174 | 1.61E-13 | -0.67175 | 1.59E-12 | Slco3a1 | -0.92451 | 5.85E-14 | -0.77158 | 2.59E-08 |
| Farp1 | 0.52125 | 9.82E-12 | 0.325843 | 0.010792 | Sobp | 0.524543 | 4.33E-07 | 0.445691 | 0.012442 |
| Fbln2 | 0.492324 | 1.57E-12 | 0.427498 | 1.19E-07 | Sox17 | -0.5613 | 4.64E-07 | -0.56634 | 3.59E-07 |
| Fbn2 | 0.741168 | 1.02E-16 | 0.6699 | 1.47E-12 | Sox5 | 1.246096 | 9.36E-15 | 1.276606 | 1.14E-16 |
| Flt1 | -0.77368 | 9.78E-19 | -0.86963 | 1.42E-21 | Sox7 | -0.47332 | 1.15E-05 | -0.48898 | 8.96E-08 |
| Fndc3a | -0.50879 | 4.37E-06 | -0.56965 | 1.02E-08 | Sox9 | 0.691842 | 5.96E-05 | 0.724249 | 6.95E-06 |
| Frmd4a | -0.42664 | 0.013439 | -0.5365 | 4.24E-06 | Sparcl1 | -0.43831 | 2.17E-06 | -0.45279 | 3.11E-10 |
| Galk1 | -0.53922 | 1.66E-07 | -0.4952 | 7.34E-06 | Spats2l | 0.598856 | 1.34E-08 | 0.444128 | 2.24E-05 |
| Garnl3 | 1.185759 | 2.45E-12 | 0.979652 | 4.17E-14 | Spon1 | 0.839078 | 1.76E-08 | 0.700573 | 1.95E-06 |
| Gata4 | 0.516105 | 4.54E-14 | 0.333078 | 1.41E-05 | St3gal4 | 0.708271 | 4.27E-12 | 0.637164 | 8.75E-08 |

| | | | | | | | | | |
|---|---|---|---|---|---|---|---|---|---|
| Gcnt1 | 0.405314 | 0.008909 | 0.437135 | 0.017426 | St6gal1 | -0.79201 | 1.77E-12 | -0.88967 | 1.24E-13 |
| Gja5 | -0.50089 | 0.039036 | -0.62048 | 2.32E-05 | St6galnac3 | -0.63508 | 2.89E-14 | -0.80294 | 3.09E-22 |
| Gm26883 | -1.3845 | 2.78E-28 | -0.91887 | 5.23E-14 | St8sia1 | -0.52058 | 0.002937 | -0.48069 | 0.045055 |
| Gm26917 | 0.58031 | 3.51E-13 | 0.273175 | 0.000496 | Sulf1 | 0.842585 | 1.89E-05 | 0.753874 | 2.07E-05 |
| Gm42906 | 1.240348 | 2.43E-17 | 0.972796 | 1.19E-16 | Tbx20 | 0.887918 | 1.38E-29 | 0.708572 | 8.95E-20 |
| Gng11 | -0.43937 | 0.002628 | -0.42967 | 0.00011 | Tcf4 | -0.56382 | 1.07E-10 | -0.61887 | 2.53E-13 |
| Greb1 | 0.572709 | 4.41E-11 | 0.395614 | 0.000355 | Tfpi | -0.94722 | 1.21E-15 | -0.90217 | 6.99E-15 |
| Grk5 | 0.755902 | 1.07E-17 | 0.692142 | 3.10E-14 | Tgfb2 | 0.448067 | 0.02515 | 0.506783 | 0.001586 |
| Hapln1 | 0.414958 | 5.48E-05 | 0.602512 | 4.10E-19 | Tgfbr3 | -0.42673 | 4.35E-05 | -0.59755 | 1.61E-08 |
| Hey1 | -0.42427 | 9.06E-06 | -0.43414 | 1.11E-05 | Tie1 | -0.42308 | 1.49E-05 | -0.47103 | 2.10E-08 |
| Hk2 | -1.03908 | 9.71E-20 | -0.96031 | 2.00E-16 | Tjp1 | -0.50569 | 2.19E-08 | -0.50981 | 3.27E-08 |
| Hmcn1 | -0.87899 | 0.00079 | -0.95266 | 7.62E-08 | Tmem100 | -1.3278 | 8.48E-32 | -1.35803 | 2.49E-32 |
| Id2 | 0.41505 | 0.045097 | 0.523165 | 0.000128 | Tmem108 | 0.750212 | 8.03E-15 | 0.611078 | 5.37E-08 |
| Ier3 | -0.48821 | 0.001315 | -0.51776 | 1.80E-07 | Tmem164 | -0.4721 | 2.28E-06 | -0.58666 | 4.01E-09 |
| Igfbp2 | -0.61499 | 1.71E-12 | -0.6107 | 1.59E-12 | Tnc | 0.568146 | 0.018096 | 0.679014 | 1.22E-06 |
| Igfbp3 | -0.84056 | 3.17E-15 | -0.92749 | 1.83E-19 | Tnfaip2 | -0.91127 | 2.03E-20 | -0.74445 | 1.48E-11 |
| Irs1 | 0.628031 | 2.31E-06 | 0.662198 | 2.57E-09 | Tnfaip8 | 0.416904 | 0.002703 | 0.350621 | 0.008947 |
| Kdm3a | -0.46865 | 0.000195 | -0.47889 | 1.61E-05 | Trpc6 | -0.53752 | 0.004702 | -0.65296 | 1.03E-05 |
| Kdm7a | -0.60511 | 4.37E-08 | -0.57763 | 2.93E-09 | Tshz2 | -1.2127 | 8.97E-22 | -1.02136 | 7.49E-17 |
| Kdr | -0.65572 | 2.16E-08 | -0.70446 | 5.70E-07 | Ttn | 0.404814 | 0.007608 | 0.39653 | 4.69E-06 |
| Kit | -0.96774 | 2.10E-18 | -1.00311 | 3.83E-20 | Twist1 | 0.478588 | 2.35E-11 | 0.562294 | 3.74E-09 |
| Kitl | 1.00753 | 5.19E-21 | 0.850278 | 2.59E-15 | Unc5c | 0.718209 | 4.27E-07 | 0.714466 | 0.000134 |
| Klhl2 | -0.62083 | 5.48E-06 | -0.69204 | 1.42E-07 | Uty | -0.8123 | 1.15E-10 | -0.54248 | 0.001777 |
| Klhl4 | -0.765 | 1.77E-11 | -0.7418 | 2.73E-09 | Vcan | 1.224913 | 3.63E-20 | 1.17047 | 2.92E-19 |
| Lama3 | -0.63442 | 1.01E-07 | -0.62578 | 4.07E-10 | Vegfa | -0.78546 | 9.96E-15 | -0.76405 | 1.32E-13 |
| Lcp1 | -0.50557 | 0.000753 | -0.4928 | 4.44E-06 | Vegfc | -0.79271 | 2.22E-07 | -0.8452 | 1.13E-07 |
| Ldb2 | -1.39044 | 9.73E-21 | -1.47873 | 1.50E-22 | Vwf | -0.81181 | 7.25E-06 | -0.80611 | 2.98E-06 |
| Lmo7 | 0.565289 | 7.03E-11 | 0.48526 | 9.85E-10 | Wdr17 | -0.74345 | 4.57E-11 | -0.56884 | 0.000181 |
| Malat1 | -0.75316 | 2.21E-29 | -0.33237 | 0.001725 | Wnt4 | 0.420794 | 0.002232 | 0.419833 | 0.000601 |
| Mast4 | -0.76369 | 7.94E-23 | -0.71759 | 3.73E-18 | Xist | 0.822021 | 6.69E-11 | 0.591109 | 5.86E-05 |
| Mest | -0.46468 | 2.00E-06 | -0.53318 | 6.65E-08 | Zfp711 | -0.62262 | 3.62E-05 | -0.59207 | 0.00276 |
| Mpped2 | 0.541571 | 3.23E-12 | 0.516696 | 3.00E-09 | | | | | |
| Msx1 | 0.539758 | 2.15E-09 | 0.459123 | 3.35E-08 | | | | | |
| mt-Atp6 | 0.441724 | 6.00E-11 | 0.577415 | 3.45E-11 | | | | | |
| mt-Co3 | 0.541115 | 2.28E-15 | 0.517791 | 3.31E-05 | | | | | |

| Table S4 | | | | | | |
|---|---|---|---|---|---|---|
| Quiescent Endocardium to Early EndoMT | | Early EndoMT to Mid EndoMT | | Mid EndoMT to Late EndoMT | | |
| Cilia KO don’t increase | Cilia KO don’t decrease | Cilia KO don’t increase | Cilia KO don’t decrease | Cilia KO don’t increase | Cilia KO don’t decrease | |
| Colec12 | Slc46a3 | Trpm3 | Rem1 | Sox5 | Gata2 | Tspan18 |
| Chst11 | Vamp1 | Bmp6 | Fus | Gpc3 | Nipal2 | Tie1 |
| Rgs7bp | Mapk4 | Tox | Gas6 | Esrrg | Rab27a | Fn1 |
| Adamts19 | 4930412O13Rik | Sox9 | Fam167a | Cdon | Eva1a | Klhl6 |
| Pappa | Mctp1 | Tll1 | Ralb | Erbb4 | Smagp | Ptprb |
| Tmtc2 | Abcg1 | Ptprd | Nostrin | Aff3 | Cmtm8 | Col4a2 |
| Rps6ka2 | Mall | Col23a1 | Nuak2 | Chst11 | Dusp3 | Ablim3 |
| Igfbp5 | Rprm | Sox6 | Anxa7 | Irs1 | Sema3g | Iqgap2 |
| Itpr1 | Pde9a | Thsd7a | Ifitm10 | Clstn2 | Notch4 | Chn1 |
| Gas7 | Gm43734 | St3gal5 | Slc7a7 | Psd3 | Myct1 | St8sia1 |
| Ror2 | Lamc3 | Galntl6 | Add1 | Vcan | Gm20532 | Tnfaip2 |
| Epha4 | Lpcat2 | Frmd4a | Fgd5 | Gpc6 | Vwf | Pced1b |
| Shroom4 | Sost | Asb4 | Sh3tc2 | Pknox2 | Ccm2l | Plvap |
| Edil3 | Hsd11b2 | Prickle2 | Hn1 | Lmo4 | Clvs1 | Prickle2 |
| Rcsd1 | Gm20631 | Sat1 | Mical2 | Papss2 | Adgrf5 | Rasip1 |
| Pde4d | Ier3 | Slco3a1 | Jam2 | Pde7b | Aff4 | Corin |
| 6030498E09Rik | Nxpe4 | Timp3 | Ccdc6 | Ncam1 | Col18a1 | Cped1 |
| Hs6st2 | Myzap | Fbn2 | Smtnl2 | Epha4 | Nhsl1 | Dysf |
| Tiparp | Kcnq1 | Inpp4b | 2510009E07Rik | Pde1a | Gldc | Rgs3 |
| Ddah1 | Asap3 | Specc1 | Tmcc2 | Hs3st3a1 | Ide | Foxo1 |
| Mdfic | Abca1 | St6galnac5 | Gm42418 | Epha7 | Usp53 | Pdzd2 |
| Esrrg | Gria3 | Grik2 | Plekha6 | Setbp1 | Ecscr | Sox7 |
| Rgl1 | Tspan18 | Cacna2d1 | Ano4 | Clmp | Rapgef4 | Lrig1 |
| Mtss1 | Nxpe2 | Pde1c | Btbd9 | Mcc | Hecw2 | Slco3a1 |
| Bach2 | Gm4876 | Fat3 | Fam129b | Sipa1l1 | Rnf138 | Pde8a |
| Bcor | Adora2a | Ltbp1 | Hey2 | Dlc1 | Lrba | Exoc3l4 |
| Ankrd44 | Gab2 | Emcn | Prkd2 | Itgb5 | Mmrn2 | Plxnd1 |
| 2610035D17Rik | Npr1 | Gm26733 | Stim2 | Cthrc1 | Icam2 | Atp2b4 |
| Arhgap26 | Clec14a | Ahr | Tspan9 | Kcnt2 | Fam163a | Zfp69 |
| Snx5 | Fam124b | Ctnnd2 | Rapgef1 | St3gal4 | Ext1 | Itga1 |
| Daam1 | Gm14507 | Notch3 | Sptan1 | Serpine2 | Taf1d | Calcrl |
| Msx1 | Ptk2 | Gm16599 | Arf2 | Ctnna3 | Nos3 | Ahr |
| Man1a | Gm15738 | Rasl11b | Ptprf | Ank | Esam | Stab1 |
| Sgcd | Osbpl6 | RP24-112I14.3 | Mapk14 | Celf4 | Ptprk | Ppp3ca |
| Tanc1 | Ctsh | Gm26936 | Kank1 | Dnm3os | Gja4 | Exoc6 |
| Mapk14 | Rassf9 | Dpysl3 | Lhfp | Gcnt1 | Abhd17a | Sorbs1 |
| Ano4 | Gm7580 | Slc2a13 | Add3 | Prrx2 | Gimap6 | Ppp1r9a |
| Ctdspl | Cdc42ep3 | Cdk14 | Notch1 | Cdh11 | F11r | Lrrc58 |
| Mthfd1l | Zfp366 | Adamts6 | Rasgrp3 | Twist1 | Palm | Nrp1 |
| Lpar1 | Gm26742 | Gm15563 | Dock9 | Antxr1 | Tjp1 | Zfpm2 |
| Tmcc2 | Nipal2 | Arhgef28 | Piezo1 | Prdm6 | Tmem100 | Thsd7a |
| Stk39 | Sema3g | Sipa1l1 | Boc | Kitl | Msn | Plpp1 |
| Rere | Bin1 | Acer2 | Iqgap1 | Ror2 | Dock5 | St6galnac3 |
| Smtnl2 | Slc43a2 | Plxna4 | Plxnc1 | Eya4 | Igf2r | Mast4 |
| Arhgap10 | Dnm3 | Efna5 | Lrig1 | Gas7 | Disc1 | Itpr1 |
| Eps15 | Tm6sf1 | Ralgapa2 | Prkch | Epb41l3 | Srgap1 | Rasgrp3 |
| Mpped2 | Dll4 | Bcl2 | Ppp1r16b | Boc | Thsd1 | Col4a1 |
| Jam2 | Mmp2 | Mast4 | Tns1 | Notch2 | Ksr1 | Efnb2 |
| Map3k1 | Gm26936 | Lrrc58 | Ednrb | A630012P03Rik | Uvrag | Hmcn1 |

| | | | | | | |
|---|---|---|---|---|---|---|
| Polk | Mfng | Zfp69 | Slc8a3 | Cpeb2 | Bcl6b | Slc22a23 |
| Akap13 | Prkaa2 | Nbea | Arfgef1 | Fbln2 | Lcp1 | Acer2 |
| Phlda1 | Asxl3 | Gm26699 | Nab1 | Tanc1 | Hdac7 | Pcsk5 |
| Dpysl2 | Ptprm | Khdrbs3 | Etl4 | Specc1 | Tspan9 | Klhl4 |
| Arhgef26 | 1810011O10Rik | Pkia | B4galt1 | Zbtb20 | Rasal2 | Adgrl4 |
| Tbl1xr1 | Epas1 | Piezo2 | Ets1 | Hdac4 | Inpp5d | Erg |
| Nkd1 | Cdk14 | Slc18a2 | Ece1 | Dock7 | Inpp4b | Tmem2 |
| | Eya1 | Atxn1 | Adam19 | Ugdh | Klf10 | Dennd5b |
| | Gm27151 | Ppp1r9a | Iqgap2 | Zmiz1 | Entpd1 | Pcdh7 |
| | Ptpre | Csrp2 | Fam129a | 5033428I22Rik | Lrrc8b | Itpr2 |
| | Ushbp1 | Ankrd28 | Shroom4 | Postn | Ccnjl | Cdh5 |
| | Bmpr2 | 4930444A19Rik | Hspg2 | Tmcc3 | Peli2 | Sat1 |
| | Mvb12b | Sorcs1 | Spats2l | Id2 | Pvt1 | Tmem164 |
| | Lifr | Lrrc4 | Ppp3ca | Ssbp2 | 5330416C01Rik | Trpc6 |
| | Plxna4os3 | Hmga2 | Arhgef15 | Greb1 | Sorcs1 | Npr3 |
| | Rdx | Tenm3 | Atp2b4 | Plekha6 | Myo6 | Elmo1 |
| | Zfp608 | Gata3 | Mtss1 | Ednrb | Fli1 | Dock9 |
| | Plekhg1 | Igf1 | Ptprj | Rin2 | Pik3r1 | Shank3 |
| | Eva1a | St8sia1 | Pdzd2 | Tiam2 | Limch1 | Prex2 |
| | Tspan2 | 1110004F10Rik | Elavl4 | Map4 | Piezo2 | Afap1l1 |
| | Cnnm2 | Frmd4b | Ubash3b | Cttnbp2 | Tspan6 | Edn1 |
| | Vegfa | Prpf4b | Col13a1 | Slc1a3 | Tspan5 | Pcgf5 |
| | Fbn2 | Add2 | Meis2 | Auts2 | Cd34 | Vav3 |
| | Sox18 | RP23-184K1.3 | Erg | Ltbp1 | Pecam1 | S1pr1 |
| | Fbxl7 | Gm28376 | Arhgef3 | Tgfbi | Pard3b | Col23a1 |
| | C530008M17Rik | Aff4 | Plk2 | Ppfibp1 | Arap3 | Plagl1 |
| | Id1 | Slc16a7 | Itpr1 | Me1 | Atxn1 | Ptprm |
| | Ecscr | Edn1 | Galnt18 | Chsy1 | Ctnnd2 | Zfp521 |
| | Tgfbr3 | Myct1 | Sulf1 | Rai1 | F2r | Emcn |
| | Smurf2 | | Nrg1 | Tenm4 | Pik3c2b | Tgfbr3 |
| | Plekhg5 | | | Epb41l1 | Stxbp6 | Samd5 |
| | Cdh5 | | | Wdpcp | Lamb1 | Frmd4b |
| | Efna1 | | | Sh3tc2 | Nfatc1 | Rapgef5 |
| | Irf2 | | | Col9a3 | Dock6 | Ano1 |
| | Chn1 | | | Fat1 | Fbn1 | St6gal1 |
| | Mamld1 | | | Ptpn13 | Sh2d3c | Egfl7 |
| | Synpo | | | Farp1 | Galnt1 | Prkch |
| | Usp43 | | | Arhgef40 | Jazf1 | Flt1 |
| | Ahr | | | Prkce | Lrrc4 | Slit2 |
| | Myo10 | | | Inpp4a | | Nav3 |
| | Apbb2 | | | Grb10 | | Trpm3 |
| | Asb4 | | | Snai1 | | |
| | Rgs2 | | | Lrp1 | | |
| | Stmn2 | | | Ociad2 | | |
| | Car8 | | | Anxa2 | | |
| | Notch3 | | | Ddr1 | | |
| | Ramp2 | | | Acadsb | | |
| | Unc13b | | | Plscr1 | | |
| | Kalrn | | | E130006D01Rik | | |
| | Plvap | | | Haus7 | | |
| | Acvrl1 | | | Tbx20 | | |
| | Gata3 | | | Npvf | | |

| | | | | |
|---|---|---|---|---|
| | Ctnnd2 | | | Pcolce |
| | Rab27b | | | |
| | Lama3 | | | |
| | Adgrl3 | | | |
| | Gcnt2 | | | |
| | Pon3 | | | |
| | Stox2 | | | |
| | Zfp423 | | | |
| | Ccdc60 | | | |
| | Bmp6 | | | |
| | Prtg | | | |
| | Nr6a1os | | | |
| | Plxna4 | | | |
| | Khdrbs3 | | | |
| | Pmepa1 | | | |
| | Plcb1 | | | |
| | Palld | | | |
| | Mmrn2 | | | |
| | Abtb2 | | | |
| | Nedd9 | | | |
| | Prickle2 | | | |
| | Cyyr1 | | | |
| | Efnb2 | | | |
| | Prdm16 | | | |
| | Calcrl | | | |
| | Sncaip | | | |
| | Tll1 | | | |
| | Trpm3 | | | |

**Key resources table**

| REAGENT or RESOURCE | SOURCE | IDENTIFIER |
|---|---|---|
| Antibodies | | |
| Mouse monoclonal anti-Arl13b | Neuromab | Cat#N295B/66; RRID: AB_2877361 |
| Rabbit polyclonal anti-Arl13b | ProteinTech | Cat#17711-1-AP; RRID: AB_2060867 |
| Goat polyclonal anti-KLF4 | R&D Systems | Cat#AF3158; RRID: AB_2130245 |
| Rat monoclonal anti-CD31 | BD Biosciences | Cat#553370; RRID: AB_396660 |
| Rabbit polyclonal anti-FN | Sigma Aldrich | Cat#F3648 |
| Alexa 488 anti-mouse | Invitrogen | Cat#A21202; RRID: AB_141607 |
| Alexa 488 anti-rabbit | Invitrogen | Cat#A21206; RRID: AB_2535792 |
| Alexa 594 anti-goat | Invitrogen | Cat#A21468; RRID: AB_2535871 |
| Alexa 647 anti-rat | Invitrogen | Cat#A21472; RRID: AB_2535875 |
| Sheep polyclonal anti-DIG-AP | Roche | Cat#11093274910; RRID: AB_2734716 |
| Bacterial and virus strains | | |
| pCX-Klf2 | Barak Cohen | RRID: Addgene_66655 |
| pCX-Klf4 | Barak Cohen | RRID: Addgene_66656 |
| Biological samples | | |
| Chemicals, peptides, and recombinant proteins | | |
| 1-phenyl 2-thiourea (PTU) | Sigma | Cat#P7629 |
| Tricaine | Sigma | Cat#E10521 |
| BM-Purple | Roche | Cat#11442074001 |
| Critical commercial assays | | |
| HCR RNA-FISH (v3.0) | Choi et al.[105] | https://www.molecularinstruments.com/ |
| Chromium iX | 10x Genomics | RRID: SCR_024536 |
| Chromium Single Cell Reagent Kit (v3.1) | 10x Genomics | PN-1000268 |
| NovaSeq 6000 | Illumina | RRID: SCR_016387 |
| Deposited data | | |
| Single Nuclei RNA-seq | This paper | GEO: GSE252341 |
| Single Cell RNA-seq | De Soysa et al.[28] | GEO: GSE126128 |
| Experimental models: Cell lines | | |
| Experimental models: Organisms/strains | | |
| Mouse: *Dnah11*$^{-/-}$(*lrd*$^{\Delta neo/\Delta neo}$): C57BL/6 | McGrath et al.[51] | MGI: 3623682 |
| Mouse: *Ift20*$^{-/-}$: C57BL/6 | Jonassen et al.[98] | MGI:3817269 |
| Mouse: *Kif3a*$^{-/-}$: C57BL/6 | Marszalek et al.[49] | MGI: 1861960 |
| Mouse: *Ncx1*$^{-/-}$: C57BL/6 | Koushik et al.[39] | MGI: 2384543 |
| Zebrafish: Tg(arl13b:EGFP,myl7:EGFP): ya201Tg | Yuan et al.[101] | ZFIN: ZDB-TGCONSTRCT-150602-7 |
| Zebrafish: Tg(Kdrl:mCherry): s916Tg | Hogan et al.[100] | ZFIN: ZDB-ALT-090506-2 |
| Oligonucleotides | | |
| *Klf2* (NM_008452.2) Amplifier: B1 | Molecular Instruments | https://www.molecularinstruments.com/ |
| *Klf4* (NM_010637.3) Amplifier: B3 | Molecular Instruments | https://www.molecularinstruments.com/ |
| *Kdr* (NM_010612.3) Amplifier: B1 | Molecular Instruments | https://www.molecularinstruments.com/ |
| *Vim* (NM_011701.4) Amplifier: B4 | Molecular Instruments | https://www.molecularinstruments.com/ |
| *Emcn* (NM_001163522.1) Amplifier: B1 | Molecular Instruments | https://www.molecularinstruments.com/ |
| *Vcan* (NM_001081249.1) Amplifier: B4 | Molecular Instruments | https://www.molecularinstruments.com/ |
| Recombinant DNA | | |
| Software and algorithms | | |
| Scrublet v0.2.3 | Wolock et al.[112] | RRID: SCR_018098 |
| Python v3.8.8 | Rossum and Drake[113] | RRID: SCR_008394 |
| Cell Ranger v3.0.1 | 10x Genomics | RRID: SCR_017344 |
| R v4.1.2 | R Core Team[114] | RRID: SCR_001905 |

| | | |
|---|---|---|
| Seurat v4.3.0 | Hao and Hao et al.[116], Stuart and Butler et al.[118], Butler et al.[115], Satija and Farrell et al.[117] | RRID: SCR_016341 |
| Harmony v0.1.1 | Korsunsky et al.[119] | RRID: SCR_022206 |
| Enrichr | Chen et al.[120], Kuleshov et al.[121] | RRID: SCR_001575 |
| MEME Suite v5.5.5 | Bailey et al.[61] | RRID: SCR_001783 |
| ImageJ (Fiji) | Schindelin et al.[108] | RRID: SCR_003070 |
| Fluorender | Wan et al.[109] | RRID: SCR_014303 |
| Imaris | Oxford Instruments | RRID: SCR_007370 |
| Other | | |
| Hoescht 33342 (1:2000) | Thermo Fisher | Cat#62249 |
| UCSC Cell Browser | Spier et al.[122] | https://cells.ucsc.edu/ |
